# Recycling upstream redox enzymes expands the regioselectivity of cycloaddition in pseudo-aspidosperma alkaloid biosynthesis

**DOI:** 10.1101/2022.09.13.507834

**Authors:** Mohamed O. Kamileen, Matthew D. DeMars, Benke Hong, Yoko Nakamura, Christian Paetz, Benjamin R. Lichman, Prashant D. Sonawane, Lorenzo Caputi, Sarah E. O’Connor

## Abstract

Nature uses cycloaddition reactions to generate complex natural product scaffolds. Dehydrosecodine is a highly reactive biosynthetic intermediate that undergoes cycloaddition to generate several alkaloid scaffolds that are the precursors to pharmacologically important compounds such as vinblastine and ibogaine. Here we report how dehydrosecodine can be subjected to redox chemistry, which in turn allows cycloaddition reactions with alternative regioselectivity. By incubating dehydrosecodine with reductase and oxidase biosynthetic enzymes that act upstream in the pathway, we can access the rare pseudo-aspidosperma alkaloids, pseudo-tabersonine and pseudo-vincadifformine, both *in vitro* and by reconstitution in the plant *Nicotiana benthamiana* from an upstream intermediate. We propose a stepwise mechanism to explain the formation of the pseudo-tabersonine scaffold by structurally characterizing enzyme intermediates, and by monitoring the incorporation of deuterium labels. This discovery highlights how plants use redox enzymes to enantioselectively generate new scaffolds from common precursors.

---

Alkaloid producing plants in the Apocynaceae family have evolved cyclases that catalyze the cycloaddition of a highly reactive substrate, dehydrosecodine (**1**), into distinct alkaloid scaffolds.^1,2^ We and others recently discovered and characterized the only known enzymes that catalyze cycloaddition of dehydrosecodine (**1**): tabersonine synthase (*Ti*TabS and *Cr*TS), which catalyzes the formation of (–)-tabersonine (**2**) (precursor to anti-cancer drugs vinblastine and vincristine), catharanthine synthase (*Cr*CS), which catalyzes formation of (+)-catharanthine (**3**) (precursor to vinblastine and vincristine), and coronaridine synthase (*Ti*CorS), which catalyzes formation of (–)-coronaridine (**4**) (precursor to anti-addiction agent ibogaine) (**Figure 1**).^2–5^ *Ti*TabS/*Cr*TS and *Cr*CS directly yield (–)-tabersonine (**2**) and (+)-catharanthine (3) from dehydrosecodine (1) by a formal [4+2] cycloaddition,^2,6^ while *Ti*CorS initially forms a hitherto uncharacterized unstable intermediate, which is then enzymatically reduced to (–)-coronaridine (**4**). The dehydrosecodine (**1**) substrate could, in principle, undergo alternative cycloaddition reactions to yield additional scaffolds but extensive mutagenesis of these cyclases did not result in an expansion of the enzymatic product profile.^2^ Here we show that the cyclase *Ti*CorS can, in addition to generating (–)-coronaridine (**4**), also produce the alternative pseudo-aspidosperma (Ψ-aspidosperma) type alkaloid pseudo-tabersonine (Ψ-tabersonine) (**5**) (**Figure 1**). We show the mechanistic basis behind this transformation by first characterizing the unstable intermediate produced by the cyclase *Ti*CorS. This intermediate can be intercepted by a reductase to generate (–)-coronaridine (**4**), or by both a reductase and an oxidase to generate the alternative scaffold Ψ-tabersonine (**5**). Deuterium labeling provides evidence for the mechanism of these enzymatic transformations. In short, the chemical reactivity of dehydrosecodine (**1**) is exploited by both a cyclase, as well as a pair of redox enzymes that can isomerize the alkene moieties, which in turn facilitates new cyclization regioselectivity.

**Figure 1.**
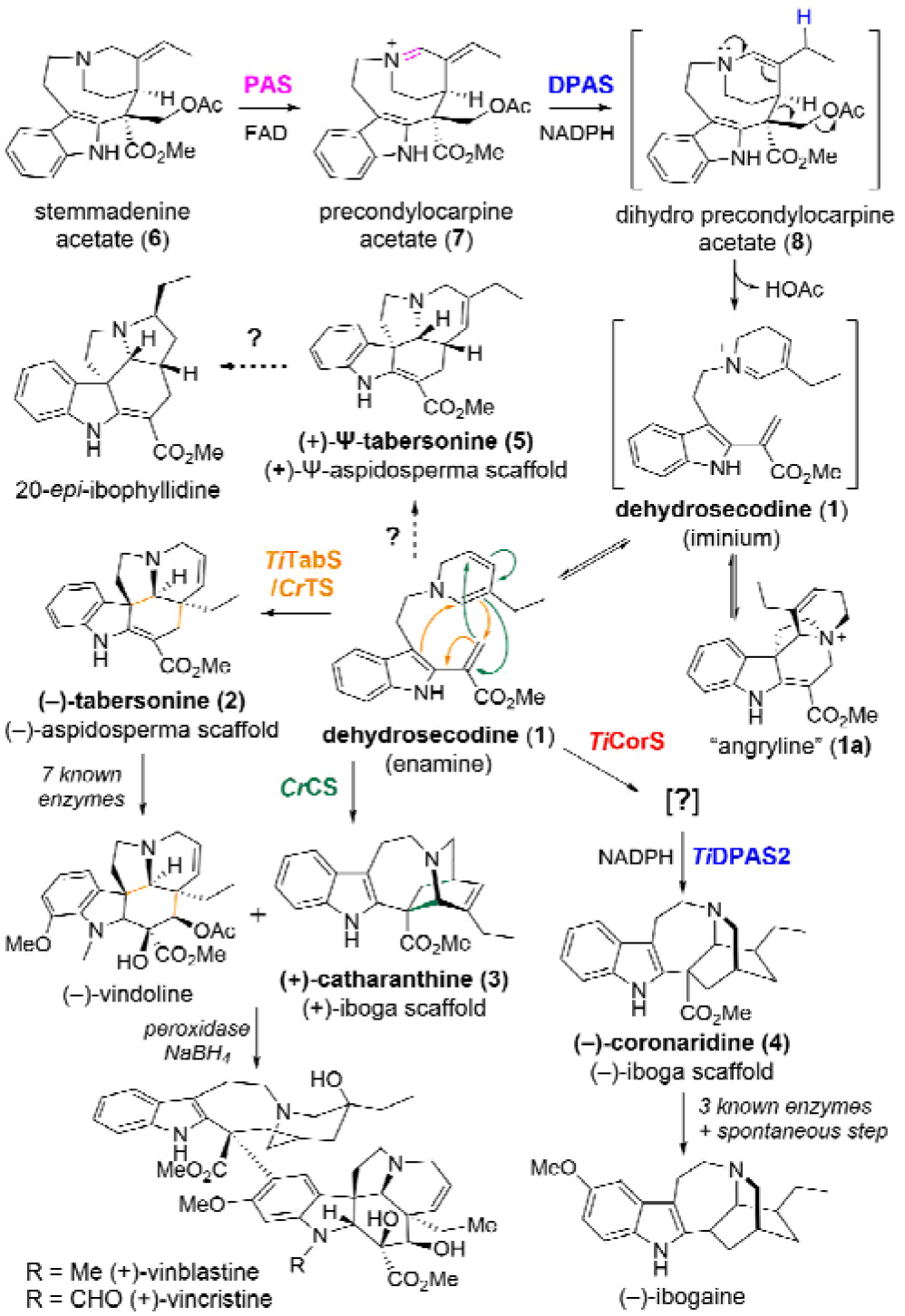
Dehydrosecodine (**1**)-derived alkaloids produced by the Apocynaceae family of plants.

*Tabernanthe iboga*, a plant that produces (–)-tabersonine (**2**) and (–)-coronaridine (**4**) via enzymes *Ti*TabS and *Ti*CorS, respectively,^3^ also produces Ψ-aspidosperma-type alkaloid 20-epi-ibophyllidine (**Figure 1**).^7–9^ We hypothesized that (+)-Ψ-tabersonine (**5**) would be the precursor to 20-epi-ibophyllidine.^10^ The natural product (+)-Ψ-tabersonine (**5**), rarely observed in nature, has only been isolated from *Tabernaemontana calcarea*, a species closely related to *T. iboga*.^11,12^ To identify *T. iboga* enzymes that could form (+)-Ψ-tabersonine (**5**), we performed coupled *in vitro* biochemical assays in which the unstable dehydrosecodine (**1**) substrate is enzymatically generated from the upstream intermediate stemmadenine acetate (**6**). Stemmadenine acetate (**6**) is first oxidized b the flavin dependent enzyme precondylocarpine acetate synthase (PAS)^5^ to generate precondylocarpine acetate (**7**), and then undergoes a 1,4-iminium reduction by the medium chain alcohol dehydrogenase dihydroprecondylocarpine acetate synthase (DPAS) (**Figure 1**).^13^ The resulting reduced unstable product, dihydroprecondylocarpine acetate (**8**), undergoes elimination of an acetoxy group to yield dehydrosecodine (**1**), which is then captured by one of the cyclases (**Figure 1**).

Our initial hypothesis was that a dedicated cyclase in *T. iboga* would catalyze isomerization and cyclization of dehydrosecodine (1) to Ψ-tabersonine (**5**). The *T. iboga* transcriptome, which contains the two previously identified cyclases *Ti*TabS and *Ti*CorS (81% sequence identity, Figure S1) does not harbor any additional cyclase homologues that might have alternative cyclization specificity. However, to our surprise, we observed that under certain conditions, Ψ-tabersonine (**5**), rather than (–)-coronaridine (**4**), was formed in assays using *Ti*CorS (**Figure 2a**). Therefore, *Ti*CorS appears to be a multi-functional cyclase, capable of producing both (–)-coronaridine (**4**) and Ψ-tabersonine (**5**).

**Figure 2.**
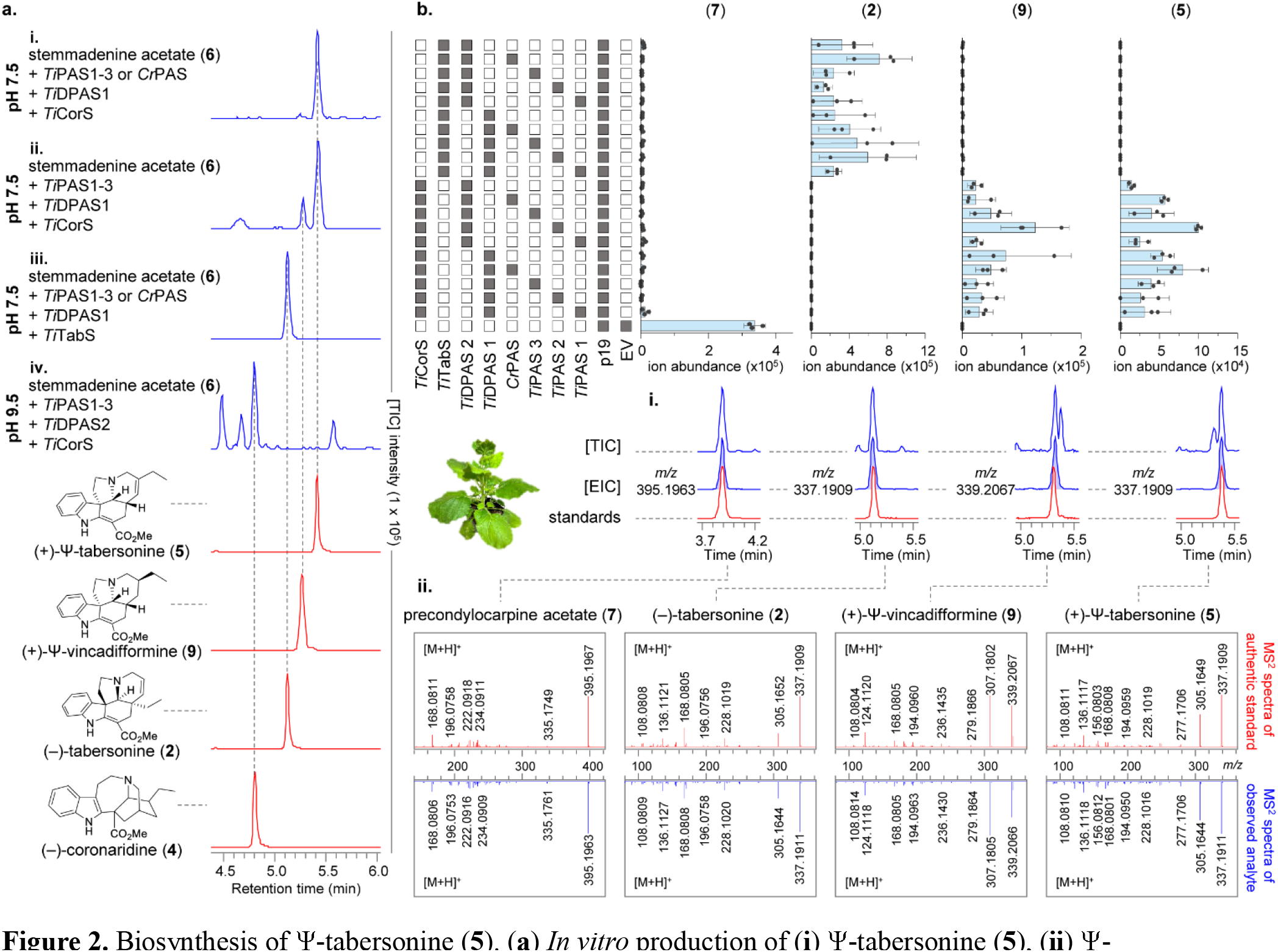
Biosynthesis of Ψ-tabersonine (**5**). (**a**) *In vitro* production of (**i**) Ψ-tabersonine (**5**), (**ii**) Ψ-vincadifformine (**9**), (**iii**) (–)-tabersonine (**2**) and (**iv**) (–)-coronaridine (**4**). (**b**) Reconstitution of Ψ-tabersonine (**5**) biosynthesis in *N. benthamiana* from stemmadenine acetate (**6**). Pathway enzyme combinations (filled grey boxes) transiently expressed in *N. benthamiana*, and harvested leaf disks fed with stemmadenine acetate (**6**) and the resulting levels of precondylocarpine acetate (**7**), (–)-tabersonine (**2**), Ψ-vincadifformine (**9**) and Ψ-tabersonine (**5**) (Bars represent ± S.E.). (**i**) Representative TIC and EIC along with authentic standard for products observed. (**ii**) MS^2^ fragmentation spectra of the product observed in the reconstitution alongside authentic standard.

*In vitro* assays using heterologously produced proteins (Figure S2) were used to probe the conditions that led to a switch in product selectivity. We first noted that the use of specific homologues of reductase DPAS and oxidase PAS with *Ti*CorS led to changes in product profile. *T. iboga* has two homologues of the reductase DPAS (*Ti*DPAS1, *Ti*DPAS2, Figure S3), and three homologues of PAS (*Ti*PAS1, *Ti*PAS2, *Ti*PAS3, Figure S4).^3^ Additionally, a PAS homologue (*Cr*PAS)^5^ from the taxonomically related plant, *Catharanthus roseus*, which produces vinblastine, is also available and was tested. Most importantly, the pH of the reaction was critical, with Ψ-tabersonine (**5**) formation being observed at pH 7.5, and (–)-coronaridine (**4**) production observed at pH 9.5 (Figure S5 and S6). Overall, Ψ-tabersonine (**5**) formation was favored at pH 7.5, with *Ti*PAS1-3 or *Cr*PAS, *Ti*DPAS1 and *Ti*CorS, (–)-tabersonine (**2**) production was favored at pH 7.5, with *Ti*PAS1-3 or *Cr*PAS, *Ti*DPAS1and *Ti*TabS, and (–)-coronaridine (**4**) formation was favored at pH 9.5 with *Ti*PAS1-3, *Ti*DPAS2 and *Ti*CorS (**Figure 2a** and Figure S5 and S6). Additionally, we also could tune the reaction to produce the over-reduced version of Ψ-tabersonine (**5**), pseudo-vincadifformine (Ψ-vincadifformine) (**9**), by using *Ti*PAS1-3, *Ti*DPAS1 and *Ti*CorS (**Figure 2a** and Figure S7).

To further substantiate the results obtained *in vitro*, we reconstituted the biosynthetic enzymes reported here leading to the production of (–)-tabersonine (**2**), Ψ-tabersonine (**5**), and Ψ-vincadifformine (**9**) in *Nicotiana benthamiana*. Enzymes were transiently expressed in *N. benthamiana* leaves, disks were excised from the transformed leaf tissue, and these disks were placed in buffer containing stemmadenine acetate (**6**) (**Figure 2b**). We observed the production of (–)-tabersonine (**2**), Ψ-tabersonine (**5**), and Ψ-vincadifformine (**9**) in the extracts of the leaf disks using this expression system (**Figure 2b**). (–)-Coronaridine (**4**) was not detected in any of the enzyme combinations tested in *N. benthamiana*, presumably because of the higher pH conditions required for formation of this product. Additionally, the selectivity observed for the PAS homologues was not observed *in planta*, because *N. benthamiana* harbors an endogenous enzyme that is able to oxidize substrate stemmadenine acetate (**6**) (**Figure 2b**).^5^

An acid-stable isomer of dehydrosecodine, angryline (**1a**), can also be isolated and directly used in cyclization assays (**Figure 1**).^2^ Angryline (**1a**) must be used at a pH value above 8.5, where it will open to generate the reactive dehydrosecodine (**1**). When angryline (**1a**) was used in enzymatic assays in place of stemmadenine acetate (pH 9.5), we could observe formation of Ψ-tabersonine (**5**), (–)-tabersonine (**2**), (–)-coronaridine (**4**) and Ψ-vincadifformine (**9**) (Figure S8). Notably, both PAS (*Ti*PAS1-3, *Cr*PAS) and *Ti*DPAS1 were required for the formation of Ψ-tabersonine (**5**) and Ψ-vincadifformine (**9**), indicating that these enzymes are required for the formation of the Ψ-aspidosperma scaffold. PAS was not required for (–)-coronaridine (**4**) production, which was favored when reductase *Ti*DPAS2 was used (Figure S9 and S10).

To investigate the mechanism by which *Ti*CorS can act as a bi-functional cyclase, we first set out to characterize the initial, unstable product that is released from *Ti*CorS in the absence of reductase (**Figure 1**). We optimized conditions under which this intermediate could be isolated, and reductively trapped this compound with NaBH4. NMR analysis showed that the compound was 16-carbomethoxycleavamine (**10**), suggesting that the initial cyclization product of *Ti*CorS is 16-carbomethoxycleaviminium (**11**) (**Figure 3** and Figure S11). When the *Ti*CorS product was reduced with NaBD4, the deuterium label was incorporated at carbon 21, which would be expected for 1,2-reduction of 16-carbomethoxy-cleaviminium (**11**) (**Figure 3**). Notably, the crystal structure of related cyclase *Cr*CS (70.8% sequence identity, Figure S1) showed that CS initially forms (+)-16-carbomethoxycleaviminium (**11a**),^14^ which then subsequently cyclizes to (+)-catharanthine (**3**), though unlike *Ti*CorS, the intermediate is not released from the active site.^2^ Moreover (+)-catharanthine (**3**) can open to form (+)-16-carbomethoxycleaviminium (**11a**) under acidic conditions (Figure S12).^1,2,15^ We used CD spectroscopy to show that *Ti*CorS generates (–)-16-carbomethoxycleavamine (**10**), which is the opposite enantiomer that is generated by *Cr*CS (Figure S13).

**Figure 3.**
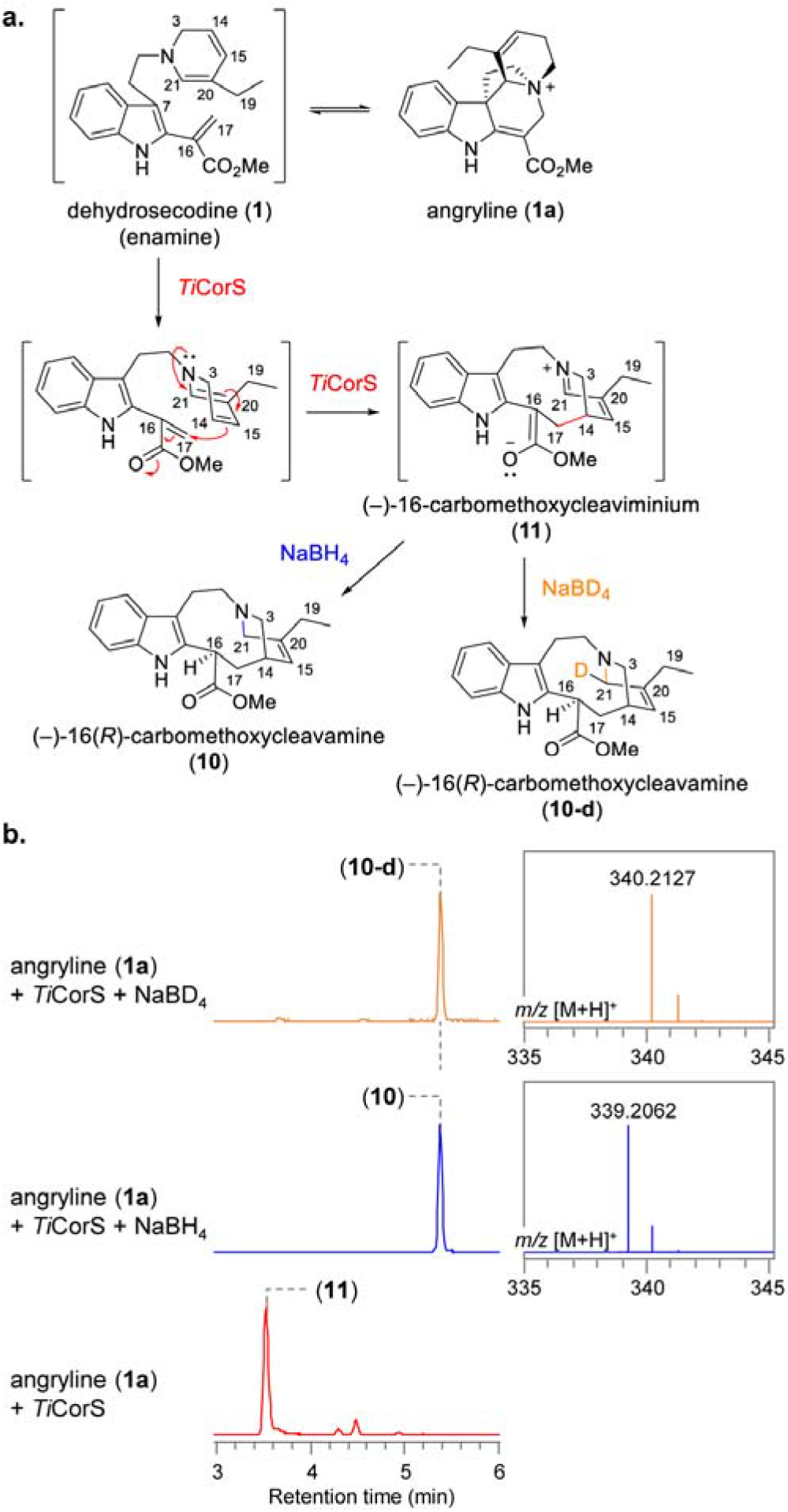
Formation of 16-carbomethoxycleavamine (**10**). (**a**) *Ti*CorS generates 16-carbomethoxycleaviminium (**11**) which could be trapped by NaBH4 to generate 16-carbomethoxy-cleavamine (**10**) or NaBD4 to generate 16-carbomethoxy-cleavamine-*d* (**10-d**). (**b**) TIC and MS spectra representing the formation of 16-carbomethoxycleaviminium (**11**) and 16-carbomethoxycleavamine (**10**) or 16-carbomethoxycleavamine-*d* (**10-d**).

With knowledge of the cyclization product of *Ti*CorS, we proposed a mechanism for the formation of (–)-coronaridine (**4**). After release from the active site of *Ti*CorS, (–)-16-carbomethoxycleaviminium (**11**) undergoes a 1,4-reduction by *Ti*DPAS2, which in turn would facilitate a second cyclization to form (–)-coronaridine (**4**) (**Figure 4**). We speculate that at higher pH values, *Ti*DPAS2 favors 1,4-reduction, which when followed by tautomerization,^16^ primes the substrate to cyclize to (–)-coronaridine (**4**).

**Figure 4.**
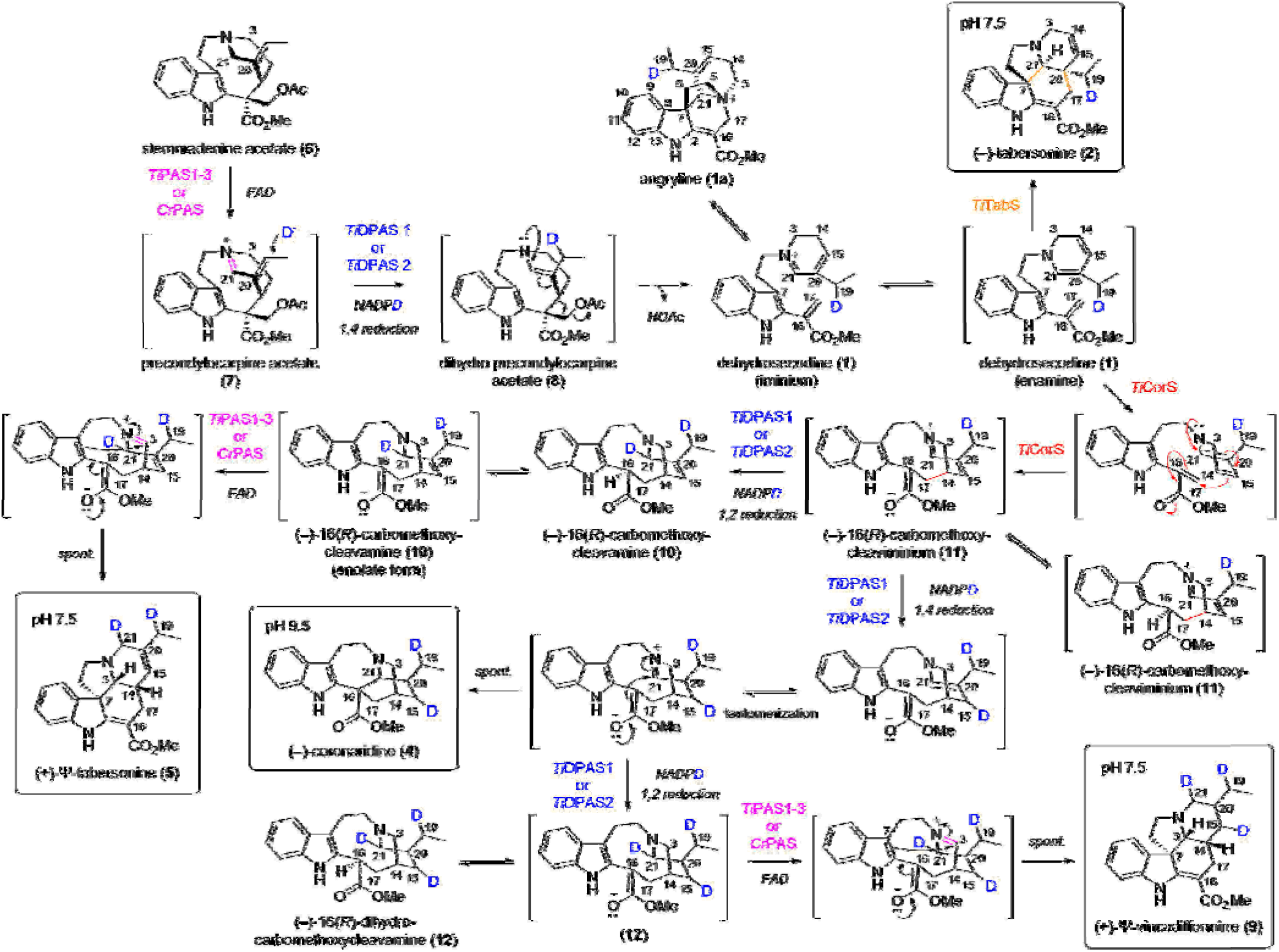
Proposed mechanism for the formation of (–)-iboga and (+)-Ψ-aspidosperma alkaloids. Biosynthesis of (–)-iboga alkaloid (–)-coronaridine (**4**), and Ψ-aspidosperma alkaloids (+)-Ψ-tabersonine (5), and (+)-Ψ-vincadifformine (9) with deuterium (blue) tracers using isotopically labelled NADPD.

However, the mechanism by which (–)-16-carbomethoxycleaviminium (**11**) could be transformed into the Ψ-tabersonine (**5**) scaffold was still not clear. Therefore, we performed the reaction in the presence of isotopically labelled NADPH (pro-(*R*)-NADPD) to determine where the deuteride is incorporated in Ψ-tabersonine (**5**).^13,17,18^ We incubated stemmadenine acetate (**6**) with *Cr*PAS, *Ti*DPAS1, and *Ti*CorS or *Ti*TabS, along with pro-(*R*)-NADPD, which is required for DPAS reduction (Figure S14). With *Ti*TabS, we saw formation of the (–)-tabersonine (**2**) product with a mass consistent with incorporation of one deuterium, as expected from the action of DPAS acting on precondylocarpine acetate (**7**) (**Figure 4**).^13^ However, when *Ti*CorS was substituted for *Ti*TabS, the resulting Ψ-tabersonine (**5**) product had a mass consistent with incorporation of two deuterium atoms (**Figure 4** and Figure S14), clearly demonstrating that formation of Ψ-tabersonine (**5**) from stemmadenine acetate (**6**) requires two reduction steps. Furthermore, the Ψ-vincadifformine (**9**) product showed a mass incorporation of three deuterium atoms (**Figure 4** and Figure S14). To corroborate these results, we also incubated the trapped isomer of dehydrosecodine (**1**), angryline (**1a**) with PAS, *Ti*DPAS1, the labeled NADPD cofactor, and *Ti*TabS or *Ti*CorS. The (–)-tabersonine (**2**) product produced by *Ti*TabS had no deuterium incorporation, as expected; formation of Ψ-tabersonine (**5**) and Ψ-vincadifformine (**9**) from angryline (**1a**) showed incorporation of one and two deuterium atoms respectively and was strictly dependent upon the presence of both reductase, *Ti*DPAS1, and oxidase, PAS (Figure S15).

We isolated isotopically labeled (–)-tabersonine (**2**), Ψ-tabersonine (**5**), and Ψ-vincadifformine (**9**) and showed that the deuterium labels were incorporated at carbon 19 for (–)-tabersonine (**2**) as expected,^13^ carbons 19 and 21 for Ψ-tabersonine (**5**), carbons and 19, 21, 15 for Ψ-vincadifformine (**9**) (**Figure 4)**. We performed CD spectroscopy of these isolated products and assigned the stereochemistry as (+)-Ψ-tabersonine (**5**) and (+)-Ψ-vincadifformine (**9**), which is the expected stereochemistry based on the downstream ibophyllidine products (Figure S16). Finally, we incubated the chemically 1,2-reduced *Ti*CorS product, (–)-16-carbomethoxyclevamine (**10**) with the oxidase PAS and observed formation of (+)-Ψ-tabersonine (**5**) (**Figure 4** and Figure S17). This result suggests that after PAS-catalyzed oxidation, the cyclization occurs non-enzymatically.

Together, this evidence strongly supports a mechanism for the formation of (+)-Ψ-tabersonine (**5**) (**Figure 4**). *Ti*CorS cyclizes dehydrosecodine (**1**) to (–)-16-carbomethoxycleaviminium (**11**), and releases it from the active site, where it is reduced to (–)-16-carbomethoxycleavamine (**10**) via 1,2-reduction by *Ti*DPAS1 and the re-oxidized by PAS. The resulting intermediate is then primed to spontaneously cyclize to form (+)-Ψ-tabersonine (**5**). (+)-Ψ-vincadifformine (**9**) forms by PAS oxidation of the doubly reduced (–)-16-dihydrocarbomethoxycleavamine (**12**) (**Figure 4** and Figure S7). The switch between (–)-coronaridine (**4**) and (+)-Ψ-tabersonine (**5**) is ultimately controlled by whether DPAS catalyzes a 1,4-reduction or 1,2-reduction. The changes in the assay pH conditions or protein-protein interactions may be responsible for favoring 1,2-reduction over 1,4-reduction. Alternatively, an as yet undiscovered reductase that generates (–)-coronaridine (**4**) at physiological pH may be responsible for the biosynthesis of this compound in in *T. iboga*.

We hypothesized that the (+)-16-carbomethoxycleavamine (**10a**) intermediate generated from opening of (+)-catharanthine (**3**) could serve as a precursor to (–)-Ψ-tabersonine (**5a**).^19^ However, oxidation of (+)-16-carbomethoxycleavamine (**10a**) by PAS yielded only a small amount of product, that, although having a mass and retention time consistent with (+)-Ψ-tabersonine (**5**), could not be fully characterized (Figure S17). Thus, PAS may only recognize one 16-carbomethoxycleavamine enantiomer.

Here we show how redox transformations of dehydrosecodine (**1**) enable cycloaddition reactions with alternative regioselectivity to form (–)-coronaridine (**4**), (+)-Ψ-tabersonine (**5**) or (+)-Ψ-vincadifformine (**9**). Notably, these redox enzymes, DPAS and PAS, which transform stemmadenine acetate (**6**) into dehydrosecodine (**1**), are recruited from upstream in the biosynthetic pathway. Therefore, this discovery highlights how plants can recycle enzymes for use in more than one pathway step. Future studies are required to establish how the recruitment of these upstream enzymes is controlled. Nevertheless, the detailed chemical analyses described here provide a compelling hypothesis for the mechanism by which these redox reactions and subsequent cyclizations expand the number of scaffolds produced from the versatile dehydrosecodine (**1**) intermediate.

## Author Contributions

S.E.O., M.O.K. and L.C. conceived experiments. M.O.K. per-formed cloning, protein purification, enzyme assays, product isolation, and pathway reconstitution. M.D.D. isolated CorS products. B.H. synthesized substrates and authentic standards. Y.N. and C.P. performed NMR analysis, compound characterization ECD experiments. B.R.L. and B.H advised on mechanistic hypothesis. P.D.S. designed the binary-vector (3Ω1) and provided advise on pathway reconstitution. S.E.O., M.O.K., M.D.D. and L.C. formulated mechanistic hypothesis. S.E.O., M.O.K. and L.C. wrote the manuscript with contributions of all authors.

## ACKNOWLEDGMENT

We would like to thank Chloe Langley for useful discussions on this work; Delia Ayled Serna Guerrero, Sarah Heinicke, and Maritta Kunert for assistance with mass spectrometry. Members of the Max Planck Institute for Chemical Ecology Research Green House for providing and taking care of Nicotiana benthamiana plants. We gratefully acknowledge the Max Planck Society and the European Research Council (788301) for funding. B.H. acknowledges the Humboldt Foundation.

