## Supporting Information for "Recycling upstream redox enzymes expands the regioselectivity of cycloaddition in pseudo-aspidosperma alkaloid biosynthesis"

**Table of Contents**


| | |
|---|---|
| Materials and Methods | S02-S08 |
| Supplementary Tables | S09-S13 |
| Supplementary Figures | S14-S33 |
| Synthesis of compounds | S34 |
| NMR spectra | S35-S62 |
| Supplementary References | S63 |

### Materials and Methods

#### Chemicals and Reagents.

All chemicals, kits and standard molecular biology reagents were purchased from commercial vendors. Stemmadenine was isolated by Professor Ivo J. Curcino Vieira as previously described.[1] Stemmadenine acetate (**6**) was synthesised as previously reported from stemmadenine.[1] Angryline (**1a**) was prepared as previously described from stemmadenine acetate (**6**).[2] Coronaridine (**4**) was synthesized by Professor Corey Stephenson and Rory McAtee (University of Michigan, USA), and obtained as described previously.[3] (+)-16(*S*)-Carbomethoxycleavamine (**10a**) and (+)-16(*R*)-carbomethoxycleavamine (**10a'**) were generated as previously described.[2,4] (+)-Catharanthine (**3**) was purchased from Abcam (cat no. 141419, CAS 2468-21-5). (–)-Tabersonine (cat no. T003900, CAS 4429-63-4) and (–)-vincadifformine (cat no. V314035, CAS 3247-10-7) were purchased from Toronto Research Chemicals. Pseudotabersonine (Ψ-tabersonine) (**5**) and (Ψ-vincadifformine) (**9**) were synthesized as described below (see section Synthesis of compounds).

#### Plant Growth.

*Tabernanthe iboga* plants were obtained and maintained as previously described[5], in a controlled environment with periodic watering. Cuttings were used to propagate new plants for RNA isolation and cDNA synthesis. The *Nicotiana benthamiana* plants utilized for transient gene expression were grown in a greenhouse on standard soil mix under a 16-h-light/8-h-dark photoperiod at 22 °C and 60% relative humidity. Plants were grown for 3-4 weeks before *Agrobacterium* infiltration, with periodic watering, as needed.

#### Cloning methods.

All genes described in this study (**Table S1**) were amplified by PCR from a cDNA library generated from *T. iboga* as previously described[5] using the primers listed in **Table S2**. P19-TBSV was amplified from clone GB0038 obtained by GB cloning website (https://gbcloning.upv.es)[6]. *Cr*PAS was amplified from a previously reported construct[1]. Overhangs compatible with the appropriate vector were included on the 5' and 3' end of the primers used for PCR amplification (**Table S2**).

High-Fidelity Q5 Hot-Start 2X Master Mix (New England Biolabs) was used for gene amplification according to the manufacturer's instructions. Following PCR amplification, DNA products were analyzed on 1% agarose gel, and the excised gel fragments were purified using Zymoclean Gel DNA Recovery Kit (Zymo Research). In-Fusion HD cloning kit (Takara Bioscience) was used to ligate all amplified genes into target vectors according to the manufacturer's instructions.

For heterologous expression of proteins in *Escherichia coli*, purified PCR amplicons of *Ti*DPAS1, *Ti*DPAS2, *Ti*CorS, and *Ti*TabS were cloned into pOPINF plasmid (Carb$^R$)[7]. The pOPINF plasmid was linearized using *HindIII* and *KpnI* restriction enzymes (New England Biolabs) to create *N*-terminal $His_6$-tagged construct suitable for affinity purification. The In-Fusion assembly products were used to transform *E. coli* TOP10 cells (ThermoFisher) by heat-shock. Positive transformants were selected on LB agar (100 µg/mL carbenicillin) and colonies were screened by colony PCR (pOPINF sequencing primers, **Table S2**) using Phire Hot Start II 2X Master Mix (ThermoFisher) according to the manufacturer's instructions. Plasmid DNA was isolated from positive colonies using Wizard Plus SV Minipreps DNA Purification Kit (Promega) and verified by Sanger sequencing. Sequence verified constructs were then used to transform *E. coli* BL21 (DE3) (ThermoFisher) expression cells. Positive transformants were confirmed by colony PCR as described above and were then used to inoculate 10 mL liquid LB medium (100 µg/mL carbenicillin), which were grown overnight at 37 °C, 200 rpm. These cultures were then used to make 25% glycerol stocks to be stored at -80 °C for further use.

For transient expression in *N. benthamiana*, purified PCR amplicons were cloned into a modified 3Ω1[6] binary plasmid (Spec$^R$) and transferred into *Agrobacterium tumefaciens* GV3101 cells by electroporation. The 3Ω1 binary plasmid modification includes a *CCDB* gene cassette between ubiquitin-10 promoter and ubiquitin-10 terminator instead of *LacZ* (present in classical 3Ω1 plasmid). For cloning with In-Fusion HD assembly kit

(Takara Bioscience), the modified 3Ω1 was linearized using *BsaI* restriction enzyme (New England Biolabs) and the assembly was carried out with the PCR amplicons as per manufacturer's instructions. The In-Fusion reaction product was transferred into *E. coli* TOP10 (ThermoFisher) cells. Positive transformants were identified by using colony PCR as described above (3Ω1 sequencing primers, **Table S2**) and then grown overnight in liquid LB medium (200 µg/mL spectinomycin) at 37 °C, 200 rpm. Plasmid DNA was isolated and sequence verified as described above. Sequence verified constructs were used to transform electrocompetent *A. tumefaciens* GV3101 cells (Goldbio) by electroporation. Cells were plated on LB agar medium (20 µg/mL rifampicin, 30 µg/mL gentamicin, 200 µg/mL spectinomycin) and grown at 28 °C for 2 days. Positive colonies were identified by colony PCR and a single transformant for each construct was grown in liquid LB medium (20 µg/mL rifampicin, 30 µg/mL gentamicin, 200 µg/mL spectinomycin) at 28 °C, 250 rpm overnight, after which 25% glycerol stocks were prepared and stored at -80 °C for further use. PAS (FAD-dependent berberine bridge enzyme) enzymes *Cr*PAS, *Ti*PAS1, *Ti*PAS2, and *Ti*PAS3 were overexpressed in *N. benthamiana* and then purified by affinity chromatography. A C-terminal $His_6$-tag together with a GSGSS linker was introduced into the cloning primers and then cloned into the modified 3Ω1 plasmid as described above (**Table S2**).

**Transient expression of candidate genes in *N. benthamiana.***

The *Agrobacterium* strains harboring gene constructs of interest were grown from glycerol stocks in 10 mL liquid LB medium (20 µg/mL rifampicin, 30 µg/mL gentamicin, 200 µg/mL spectinomycin) overnight at 28 °C, 250 rpm. Cells were harvested by centrifugation at 2,000 *g* for 15 min and the depleted media was discarded. The cell pellet for each strain was gently resuspended in *Agrobacterium* infiltration buffer (10 mM MES, pH 5.6, 10 mM $MgCl_2$, 200 µM acetosyringone) and washed to remove residual media and antibiotics. Washed cells were harvested by centrifugation at 2,000 *g* for 15 min and the buffer discarded. Cells were resuspended in 10 mL of *Agrobacterium* induction buffer and incubated in the dark at room temperature with gentle rocking for 2 h. When multiple gene constructs were tested in combinations, strains were pooled in equal cell density so that each strain had an $OD_{600}$ = 0.4 and infiltrated into the abaxial side of 3-week-old *N. benthamiana* leaves covering the entire leaf (approximately 1.5 mL per leaf) using a 1-mL needleless syringe. Each pooled combination consisted of three biological replicates, where an individual leaf (fully expanded) belonging to three different plants was infiltrated to randomize batch effects. Leaves were harvested 3 days post-*Agrobacterium* infiltration for *in vitro N. benthamiana* leaf disk assays (see section *In vitro* assays with *Agrobacterium* infiltrated *N. benthamiana* leaf disks) to test for enzymatic activity.

For the overexpression of PAS enzymes with a *C*-terminal $His_6$-tag, the corresponding *Agrobacterium* strain was prepared as described above and co-infiltrated with P19-TBSV strain at an equal cell density corresponding to a final $OD_{600}$ = 0.3. Prepared *Agrobacterium* suspension was infiltrated (covering the entirety of the leaves) into the abaxial side of 20 3-week-old *N. benthamiana* plants using a 1-mL needleless syringe and three independent leaves per plant were selected to generate sufficient biomass for protein extraction. Leaves were harvested 3 days post-*Agrobacterium* infiltration; the central vascular tissue was excised using a razor blade and the plant material was snap frozen in liquid $N_2$ and stored at -80 °C for protein purification (see section Purification of $His_6$-tagged proteins from *N. benthamiana*).

**Expression and purification of proteins in *E. coli.***

For recombinant protein production in *E. coli*, starter cultures of BL21(DE3) were prepared from glycerol stocks (see section cloning methods) in 10 mL LB medium (100 µg/mL carbenicillin) and grown at 37 °C, 200 rpm for 16 h. The starter cultures (10 mL) were used to inoculate 1 L of 2xYT medium (100 µg/mL carbenicillin) and the resulting large cultures were grown to an optical density at 600 nm ($OD_{600}$) of 0.7 at 37 °C, 200 rpm. Cultures were then chilled on ice for 20 min, induced with 0.25 mM isopropyl β-D-1-thiogalactopyranoside (IPTG) and incubated overnight (18 h) at 18 °C, 200 rpm for protein production. Cells were harvested by centrifugation at 3,000 *g*, 4 °C, for 10 min and resuspended in 50 ml of pre-cooled buffer A (50 mM Tris-HCl pH 7.4, 50 mM glycine, 500 mM NaCl, 5% glycerol, 20 mM imidazole) with 10 mg/mL lysozyme and an EDTA-free protease inhibitor (Roche Diagnostics). All remaining steps were performed on ice and all buffers were pre-cooled at 4 °C. Cells were lysed by sonication for a total of 5 min (2 sec on, 3 sec off cycle) on ice. The resulting crude protein lysate was clarified by centrifugation at 35,000 *g*, 4 °C, for 20 min and filtered through a 0.45-µm glass filter. $His_6$-tagged proteins were purified on an ÄKTA Pure 25 fast protein liquid chromatography (FPLC) system (Cytiva) using a 5 mL-HisTrap-HP column (Cytiva) equilibrated with buffer A. Sample was applied at a flow rate of 2 mL/min and step-eluted using buffer B (50 mM Tris-HCl pH 7.4, 50 mM glycine, 500 mM NaCl, 5% glycerol, 500 mM imidazole) and collected in 1 mL fractions. The principal peaks collected were verified by SDS-PAGE with Coomassie Blue staining. Fractions containing the protein of interest were pooled and concentrated using 15 mL-Amicon-Ultra 10 kDa molecular weight cutoff filters (Millipore) by centrifugation at 3,000 *g*, 4 °C, for 10 min intervals with gentle resuspension till the sample was concentrated to 1 mL. Samples were dialyzed in the same filter by the addition of 12 mL buffer C (20 mM HEPES pH 7.4, 150 mM NaCl) and centrifuged as described above. Dialysis was repeated 2 more times and the protein was concentrated to a final volume of 0.5 mL. Protein concentrations were measured using a nano-photometer (Implen) based on the absorbance at 280 nm using the theoretical extinction coefficients. Proteins were aliquoted (50 µL), snap-frozen in liquid $N_2$ and stored at -80 °C for *in vitro* enzyme assays.

**Purification of $His_6$-tagged proteins from *N. benthamiana*.**

To purify *C*-terminal $His_6$-tag PAS (oxidase) enzymes from *N. benthamiana* leaves, the harvested tissue (see section Transient expression of candidate genes in *N. benthamiana*) was ground to fine powder in liquid $N_2$ using a mortar and pestle. The powdered tissue (ca. 60 g) was extracted with 360 mL of ice-cold buffer A supplemented with 1 mM PMSF, EDTA-free protease inhibitor (Roche Diagnostics), 2% w/v polyvinylpolypyrrolidone (PVPP). The extract was mixed by inversion and incubated at 4 °C for 30 min with stirring to maximize homogeneity and extraction efficiency. All remaining steps were performed on ice and all buffers were pre-cooled at 4 °C along with consumables. The extract was clarified by centrifugation at 3,000 *g*, 4 °C, for 10 min, and filtered through miracloth (Millipore). The flow through was centrifuged at 35,000 *g*, 4 °C, for 20 min and the cleared protein extract was passed through 0.45-µm glass filters before Ni-affinity purification. $His_6$-tagged crude protein lysate was purified on an ÄKTA Start protein purification system (Cytiva) using a 5 mL-HisTrap-HP column (Cytiva) conditioned with buffer A. Sample (ca. 300 mL) was applied at a flow rate of 2 mL/min, step-eluted with buffer B, and collected as 1 mL fractions. The fractions were verified for the protein of interest using SDS-PAGE with Coomassie Blue staining. The pooled fractions of interest were dialyzed in buffer C using PD-10 desalting columns (Cytiva) and concentrated to a final volume of 1 mL using 15 mL-Amicon-Ultra 10 kDa molecular weight cutoff filters (Millipore) by centrifugation in 10 min cycles at 3000 *g*, 4 °C with gentle resuspension at the end of each round. Protein concentration was normalized to 8 mg/mL and prepared for storage as described (see section Expression and purification of proteins in *E. coli*).

***In vitro* assays with *Agrobacterium* infiltrated *N. benthamiana* leaf disks.**

At three days post-*Agrobacterium* infiltration (see section Transient expression of candidate genes in *N. benthamiana*), 10 mm leaf disks were cut using a borer and placed into 48-well-plates (10 mm diameter well). For each experiment, disks were collected from three independent plants. The wells contained 250 µL of 50 mM HEPES buffer at pH 7.5 and the leaves were placed (1 leaf disk per well) either on the abaxial or on the adaxial side and submerged in the buffer immediately upon harvest. Stemmadenine acetate (**6**) was used as substrate to a final concentration of 50 µM into the wells containing the leaf disks. Controls of wild-type, empty-vector, and P19-TBSV infiltrated *N. benthamiana* incubated with substrate were also included for each experiment. The plates were closed with the covers provided and sealed with parafilm to avoid excessive evaporation of the buffer. The plates were incubated in a growth chamber at 22 °C, 60% relative humidity and 16-h-light/8-h-dark photoperiod overnight (16-18h). After overnight incubation, individual leaf disks were collected and placed in a 2 mL safe-lock tube and snap-frozen in liquid nitrogen. The leaf disk was pulverized using a TissueLyser II (Qiagen) homogenizer using 3 mm tungsten beads, with shaking at 25 Hz for 1 min. To extract metabolites, 250 µL of extraction solution (70% methanol in water acidified with 0.1% formic acid) was added to the powdered tissue, mixed well by vortex and sonicated for 10 min at room temperature. The samples were then centrifuged at 20,000 *g* for 10 min to pellet cell debris and the extract was diluted 1:2 with extraction solution. Samples were filtered through 0.22 μm PTFE syringe filters before metabolite analysis by LC-MS.

***In vitro* enzymatic assays.**

The conditions used for analytical scale production of (+)-Ψ-tabersonine (**5**), (+)-Ψ-vincadifformine (**9**), (–)-tabersonine (**2**), and (–)-coronaridine (**4**) from substrates stemmadenine acetate (**6**) and angryline (**1a**) are described below. Substrates were prepared in methanol and added at a final concentration of 40 μM into a 100 μL reaction. For *in vitro* coupled enzyme assays with stemmadenine acetate (**6**) as substrate, reaction mixtures with combinations of purified enzymes were assembled to investigate the production of desired final products as described in **Table S3**. For *in vitro* coupled enzyme assays with angryline (**1a**) as substrate, reaction mixtures with combinations of purified enzymes were assembled to investigate the production of desired final products as described in **Table S4**. In experiments examining the effect of pH for the production of (–)-coronaridine (**4**) from angryline (**1a**) (**Table S4)**, appropriate buffers[8] from pH 7.5 to 9.5 at 50 mM were used in 0.5 pH increments. For all *in vitro* reactions, appropriate control assays omitting enzymes were additionally performed to test the validity of reaction products observed. Optimal pH and enzyme combination for the desired final products of (+)-Ψ-tabersonine (**5**), (+)-Ψ-vincadifformine (**9**), (–)-tabersonine (**2**), and (–)-coronaridine (**4**) from stemmadenine acetate (**6**) (**Table S3)** and angryline (**1a**) (**Table S4**) was established. The same conditions were used for stereo-specific deuterium labelling with pro-(*R*)-NADPD by replacing the NADPH co-factor with the isotopically labelled NADPD analogue. Pro-(*R*)-NADPD was generated *in situ* by the addition of reaction components described below. For 100 μL reactions, 1 mM $NADP^+$, 10 mM isopropanol-$d_8$ (Sigma-Aldrich, 175897), 1% (v/v) alcohol dehydrogenase (Sigma-Aldrich, 49641) were added, and the optimized reactions were assembled as described above excluding NADPH. Substrates, angryline (**1a**) and stemmadenine acetate (**6**) were used for deuterium labeling reactions.

*In vitro enzymatic assays for the production of (+)-Ψ-tabersonine (**5**) from (–)-16(R)-carbomethoxycleavamine (**10**).*

For assays initiated with (–)-16(*R*)-carbomethoxycleavamine (**10**) to produce (+)-Ψ-tabersonine (**5**) from PAS (oxidase) enzymes, 100 μL reactions were performed in 50 mM HEPES pH 7.5 buffer with 0.2 mM FAD, 1 μM PAS enzyme, and 40 μM substrate (–)-16(*R*)-carbomethoxycleavamine (**10**). Reactions were incubated at 37 °C, 600 rpm for 60 min and quenched. To attempt to produce (–)-Ψ-tabersonine (**5**), the above conditions were used with 40 μM substrate (+)-16(*S*)-carbomethoxycleavamine (**10a**).

To produce precondylocarpine acetate (**7**) from stemmadenine acetate (**6**), 100 μL reactions were performed. PAS (oxidase) enzymes (1 μM) were incubated in 50 mM HEPES pH 7.5 with 0.2 mM FAD, and 40 μM of substrate stemmadenine acetate (**6**). Reactions were incubated at 37 °C, 600 rpm for 60 min, quenched and subjected to LC-MS analysis (Figure S18).

All analytical scale reactions were quenched by the addition of 100 μL quenching solution (70% methanol in water + 0.1% formic acid). Quenched reactions were vortexed, centrifuged at 20,000 *g* for 10 min and filtered with 0.22 μm PTFE syringe filters for subsequent LC-MS analysis. All reactions were independently replicated 3 times to validate reproducibility.

**Preparative scale *in vitro* reaction workup for product isolation.**

*Large scale production of deuterium labeled (–)-tabersonine (**2**), (+)-Ψ-tabersonine (**5**), and (+)-Ψ-vincadifformine (**9**)*

Deuterium labeling assays with pro-(*R*)-NADPD analogue (described in the section *In vitro* enzyme assays) were scaled up to 200 μL reaction volumes. Stemmadenine acetate (**6**) at 200 μM was used as the substrate to produce the desired deuterium labeled product, with optimal pH, enzyme combinations, and assay conditions listed in **Table S3**. Multiple reactions were assembled in batches (24 reactions) to yield sufficient product for preparative HPLC isolation. Reactions were quenched with 200 µL of quenching solution, vortexed, and centrifuged at 20,000 *g* for 10 min. Clarified crude mixtures were pooled and filtered into a single vial using 0.22 µm PTFE syringe filters. Dilution (1:1000) of the crude mixture was analyzed by LC-MS and the desired deuterium labeled product was confirmed by *m/z* and retention time. The pooled mixture was evaporated to dryness using a rotoevaporator and the dried crude product was stored at -20 °C for isolation using preparative HPLC (see section Isolation of products using preparative HPLC).

*Workup for the production of (–)-coronaridine (**4**)*

Production of (–)-coronaridine (**4**) was performed using the optimal pH, enzyme combinations, and assay conditions listed in **Table S3** from stemmadenine acetate (**6**). Reactions were scaled up and performed as described above. The dried crude product was stored at -20 °C for isolation using preparative-HPLC (see section Isolation of products using preparative HPLC).

*Workup for the production of (+)-16(S)-carbomethoxycleavamine (**10a**) and (+)-16(R)-carbomethoxycleavamine (**10a'**)*

(+)-16(*S*)-carbomethoxycleavamine (**10a**) and 16(*R*)-carbomethoxycleavamine (**10a'**) were generated chemically using a modified method previously described[2,4]. Briefly, (+)-catharanthine (**3**) (5 mg) was dissolved in 400 µL of TFA and stirred at room temperature for 3 h. Conversion of (+)-catharanthine (**3**) to (+)-16-carbomethoxycleavamininum (**11a**) intermediate (*m/z* 337.19) was observed. Then, the iminium intermediate (**11a**) was reduced by the addition of an excess $NaBH_4$ in methanol (10 mg in 400 µL). The reaction was stirred at room temperature for 30 min and the 16-carbomethoxycleavamine diastereomers, 16(*S*) (**10a**) and 16(*R*) (**10a'**) were observed by LC-MS. Acetone (500 µL) was added and stirred for 10 min at room temperature. The reaction mixture was evaporated to dryness using a rotoevaporator and the dried product was stored at -20 °C for isolation using preparative HPLC (see section Isolation of products using preparative HPLC).

*Workup for the production of (–)-16(R)-carbomethoxycleavamine (**10**) and deuterium labeled (–)-16(R)-carbomethoxycleavamine (**10-d2**)*

To generate (–)-16(*R*)-carbomethoxycleavamine (**10**), a 4 mL reaction was performed with the substrate angryline (**1a**) at 2 mM final concentration in a reaction mixture consisting of 50 mM CHES buffer at pH 9.5, and 40 µM *Ti*CorS enzyme. The reaction mixture was incubated at 37 °C for 30 min, with shaking at 150 rpm prior to quenching by the addition of 4 mL quench solution to generate the iminium product. Subsequently, the iminium was reduced by the addition of 4 mM $NaBH_4$ sodium borohydride (Sigma Aldrich) (in methanol) to the quenched reaction, and incubation at 37 °C for 30 min, with shaking at150 rpm. Upon completion of the reaction, the precipitated protein was removed by centrifugation at 4000 *g*, for 10 min at room-temperature. Dilution (1:1000) of the crude reaction mixture subjected to LC-MS analysis confirmed the presence of the desired product, (–)-16(*R*)-carbomethoxycleavamine (**10**). The reaction mixture was dried in a rotoevaporator and was stored at -20 °C until purification by preparative HPLC. Generation of deuterium-labeled (–)-16(*R*)-carbomethoxycleavamine-$d_2$ (**10-d2**) was performed as described for (–)-16(*R*)-carbomethoxycleavamine (**10**). The iminium generated was reduced by the addition of 4 mM $NaBD_4$ (sodium borodeuteride) (Sigma Aldrich) in methanol. All subsequent steps were performed as described above. The crude mixture containing (–)-16(*R*)-carbomethoxycleavamine-$d_2$ (**10-d2**) was dried using a rotoevaporator and was stored at -20 °C until purified by preparative HPLC (see section Isolation of products using preparative HPLC).

**Isolation of products using preparative HPLC.**

Preparative-scale *in vitro* reactions (see section Preparative scale *in vitro* reaction workup for product isolation) were subjected to preparative HPLC for compound isolation. A preparative HPLC system (Agilent 1260 Infinity II) coupled to a multiple wavelength detector and fraction collector connected to a Phenomenex Kinetex XB-C18 (250 x 10 mm, 5 µm, 100 A) was used. The mobile phases used for separation were A (water containing 0.1% formic acid) and B (acetonitrile). The flow rate was set at 6.0 mL/min and a gradient of 10-40% B for 15 min, 90% B for 7 min, 10% B for 5 min. Samples prepared for preparative-HPLC analysis (see section Preparative scale *in vitro* reaction workup for product isolation) were resuspended in quenching solution to 1 mg/mL and filtered using a 0.22 µm PTFE syringe filter. Manual injections (200 µL per injection) were performed, and the separation was monitored using specific wavelengths. For (–)-tabersonine-*d* (**2-d**), (+)-Ψ-tabersonine-$d_2$ (**5-d2**), and (+)-Ψ-vincadifformine-$d_3$ (**9-d3**), 328 nm was used. For (+)-16(*S*)-carbomethoxycleavamine (**10a**), (+)-16(*R*)-carbomethoxycleavamine (**10a'**), (–)-16(*R*)-carbomethoxycleavamine (**10**), (–)-16(*R*)-carbomethoxycleavamine-$d_2$ (**10-d2**), and (–)-coronaridine (**4**), detection at 225 nm and 280 nm wavelengths were used. Eluted fractions were analyzed by LC-MS (1:1000 dilution) and desired fractions were pooled and evaporated to dryness. The isolated compounds were then submitted for NMR analysis (see section NMR analysis).

**LC-MS data acquisition and analysis of metabolites.**

Samples were analyzed using a Thermo Scientific UltiMate 3000 ultra-high performance liquid chromatography (UHPLC) system coupled to an Impact II UHR-Q-ToF (Ultra-High Resolution Quadrupole-Time-of-Flight) mass spectrometer (Bruker Daltonics). Metabolites were separated by reversed-phase liquid chromatography using a Phenomenex Kinetex XB-C18 (100 x 2.1mm, 2.6 µm; 100 Å) column at 40 °C. The mobile phases for metabolite separation were water with 0.1% formic acid (A) and acetonitrile (B). A flow rate of 0.6 mL/min was used for the chromatography. The injection volume was 2 µL. The chromatographic separation was performed starting at 10% B for 1 min, linear gradient from 10% to 30% B in 6 min, 90% B for 1.5 min, 10% B for 2.5 min. Authentic standards were prepared as 20 µM solutions in methanol and 2 µL was injected under the chromatographic conditions described above. Mass spectrometry acquisition was performed in positive electrospray ionization mode with a capillary voltage of 3500 V and an end plate offset of 500 V; a nebulizer pressure of 2.5 bar was used, with nitrogen at 250 °C and a flow of 11 L/min as the drying gas. Acquisition was done at 12 Hz in the mass range from 80 to 1000 *m/z*, with data dependent $MS^2$ and an active exclusion window of 0.2 min. For tandem mass spectroscopy ($MS^2$) fragmentation was triggered on an absolute threshold of 400 and limited to a total cycle time range of 0.5 s. For collision energy, the stepping option model (from 20 to 50 eV) was used. At the beginning of each sample run, a sodium formate-isopropanol calibration solution was directly infused to the source at 0.18 ml/hour using a syringe pump to calibrate MS spectra recorded. To avoid injection peak and salt contamination of the MS, the initial 1 min of the active chromatographic gradient of each run was discarded to waste.

Data analysis was performed using Bruker Compass Data Analysis (Version 5.3) software. For *N. benthamiana* leaf disk experiments, data acquired was analyzed using Bruker Compass MetaboScape 2021b software (version 7.0.1). The non-targeted metabolomics workflow was used to generate an output containing a list of mass signatures with retention times, along with qualitative peak intensities by automated integration of extracted ion chromatograms with 5 ppm tolerance, molecular formula determined by accurate mass of precursors and fragments and statistical test outputs (*P* value and *t*-tests). All metabolites in this study were verified and assigned using authentic standards by validation of retention time and $MS^2$ spectra (Figure S19). Figures for *N. benthamiana in vitro* leaf disk assays were prepared using the scientific graphing and data analysis program OriginPro 2019 (version 9.6.0) from the processed data obtained by MetaboScape analysis.

**NMR analysis.**

NMR spectra were measured on a 700 MHz Bruker Advance III HD spectrometer (Bruker Biospin GmbH, Rheinstetten, Germany). For the synthetic (±)-Ψ-tabersonine (**5a**) standard, NMR spectra were measured on a 500 MHz Bruker Advance III HD spectrometer (Bruker Biospin GmbH, Rheinstetten, Germany). For spectrometer control and data processing Bruker TopSpin (version 3.6.1) was used. MeOH-$d_3$ was used as a solvent and all NMR spectra were referenced to the residual solvent signals at $\delta_H$ 3.31 and $\delta_C$ 49.0, respectively.

**Electronic circular dichroism (ECD) spectra measurements of isolated products.**

ECD spectra were measured at 25 °C on a JASCO J-810 spectropolarimeter (JASCO cooperation, Tokyo, Japan) using a 350 µL cell. Spectrometer control and data processing was accomplished using JASCO spectra manager II. All the isolated products were measured in methanol at 0.2 mg/mL concentration.

**Electronic circular dichroism (ECD) spectra calculations.**

ECD spectra calculations were performed for (+)-Ψ-tabersonine (**5**), (+)-Ψ-vincadifformine (**9**), (–)-tabersonine (**2**), and (–)-16(*R*)-carbomethoxycleavamine (**10**). Based on the structure determined from NMR analysis a molecular model was created in GaussView ver.6 (Semichem Inc., Shawnee, Kansas, USA) and optimized using the semi-empirical method PM6 in Gaussian (Gaussian Inc., Wallingford, Connecticut, USA). The resulting structure was used for conformer variation with the GMMX processor of the Gaussian program package. Resulting structures were DFT-optimized with Gaussian ver.16 (APFD/6-31G(d) level). A cut-off level of 4 kcal/mol was used to select conformers which were subjected to another DFT optimization on a higher level (APFD/6-311G+(2d,p)). The structure of (–)-16(*R*)-carbomethoxycleavamine (**10**) was optimized using the semi-empirical method PM6 and the resulting structure was DFT-optimized (B3LYP/6-311G+(d,p)). All structures up to a deviation of 2.5 kcal/mol from the lowest energy conformer were used to determine the ECD-frequencies in a TD-SCF calculation on the same level as the former DFT optimization. The ECD curve was calculated from the Boltzmann-weighed contributions of all conformers with a cut-off level of two percent. Experimentally measured CD spectra and calculated ECD data were compared using SpecDis[9] (version 1.71).

### Supplementary Tables

**Table S1.** Nucleotide sequences for genes cloned and described in this study.

| **Gene Name (GenBank Accession))** | **Nucleotide sequence** |
|---|---|
| *Ti*DPAS 1 (MK840855.1) | **ATG**GCTGTAAAATCACCTGAAGCAGAGCACCCAGTGAAGGCTGTAAAATCACCTGAAGAAGAG<br>CACCCAGTGAAGGCATACGGATGGGCTATCAAAGACAGAACATCTGGCATTCTTTCCCCCTTCA<br>AGTTTTCCAGAAGGGCAACAGGAGATGAAGACGTTCGAATAAAGATCCTCTGTTGCGGAGTTTG<br>TCACACGGATCTCACGTCTACCAAGAATGAATACGAGTTTCTTTCATATCCTCTAGTGCCTGGGT<br>TGGAGACTGTCGGAATAGCGACAGAGGTCGGAAGCAAAGTCACAAAAGTAAAAGTTGGTGAAA<br>AAGTAGCAGTGGCAGCCTATTTGGGCACTTGTGGCAAATGCCACAATTGTCTAAATGACCAAGA<br>GAATTACTGTCCCGAAGTGATCATTAGCTACGGCACACCATATCACGACGGAACAATCAACTAC<br>GGAGGCTTCTCGAATGAGACGGTCGTAAATGAGCGCTTCGTTCTTCATTTTCCTGAAAAGCTTTC<br>ACTTTCTGGTGGTGCACCGCTACTCAGCGCAGGAAGCACCGCTTACAGTGCAATAAGAAATCAA<br>GGCCTTGACAAACCCGGTATCCACTTGGGAGTCGTCGGCCTTGGTGGACTTGGTCATCTGGCTGT<br>GAAGTTTGCCAAGGCTTTTGGTGTCAAGGTGACAGTGATTAGTTCCACTCCCAGCAAGAAGGAT<br>GAAGCCATCAAGAGCCTTGGTGCCGATGCGTTCTTGTTCAGTCGTGATGATGAACAAATGAAGG<br>CCGCTATTGGAACTTTCGATGCAATCATAGATACTATTGCAGTCGCTCATCCTCTTGCGCCATTAC<br>TTGATCTACTAAGGAGTCATGGAAAAATTATTTGGTCGGGGCACCGACCACCCCACTTGAGGTG<br>CCAGTTATTCCTTTAGTAGCAGGTGGGAAATCGATTACTGGATGCGTAGTTGGAAATTTGAAGCA<br>AACTCAAGAAATGCTTGAATTTGCTGCAGAACACAACATCACTGCAAACGTTGAGGTTATTTCA<br>ATGGATTACATAAACACTGCAATGGAACGTTTAGAAAAAGGTGATGTTAGATATAGATTTGTAA<br>TTGATATTGGAAACACACTAACTCCACCGGAATAG |
| *Ti*DPAS 2 (MK840856.1) | **ATG**GCAGGAAAATCACCAGAAGAGGAACACCCAGTGAAGGCATATGGATGGGCTGTCAAGGAC<br>AGAACAACTGGGATTCTTTCTCCCTTCAAGTTTTCGAGAAGGGCAACAGGAGACAATGACATCC<br>GAATCAAGATCCTCTATTGTGGAATTTGTCATACAGACCTAACATCTGTCAAGAACGAGTACGA<br>GTTTCTTTCATATCCTCTTGTGCCTGGGATGGAGATTGTAGGGATAGCGACAGAGGTCGGAAGCA<br>AAGTCACAAAAATAAAAGTTGGTGAAAAAGTAGCAGTAGCAGCCTATTTGGGTACTTGTGGAAA<br>ATGCTACAATTGTGTAAATGACCTTGAGAACTACTGTCCTGAAGTCATCATTGGTTATGGTACGC<br>CATATCACGATGGAACAATTAACTACGGAGGCCTCTCAAACGAGACGGTCGTAAATGAGCGCTT<br>TGTTCTTCGTTTTCCTGAAAAACTTTCACCTGCTGGTGGTGCCCCGCTACTCAGTGCTGGAATCAC<br>CGCGTACAGTGCAATGAGGAATCATGGCCTCGACAAGCCCGGAATCCAATTGGGAGTCGTCGGT<br>CTTGGTGGACTTGGTCATCTGGCTGTGAAGTTTGCCAAGGCTTTTGGCGTCAGAGTGACTGTGAT<br>TAGTACCACTCCTAGCAAGAAGGATGAAGCTATAAATAATCTTGGTGCTGATGCCTTTTTGTTCA<br>GCCGTGACGATAAGCAAATGAGGGCTGCCATTGGAACGTTTGATGCAATCATCGACACACTAGC<br>GGTTGTTCATCCTATTGCGCCCTTACTCGATCTATTGAGGAGTCATGGAAAACTTGTTTTGGTTGG<br>AGCCCCATCTAAGCCACTTGAGCTACCAACTATTCCTTTATTATCAGGAGGAAAATCATTGATCG<br>GTAGTGCAGCTGGAAATGTGAAGCAAACTCAGGAAATGCTTGATTTTGCAGCAGAACACGATAT<br>TACTGCAAACATTGAGGTTATACCAATAGATTATATAAACACTGCAATGGAACGTTTAGATAAA<br>GGTGATATCCGATTTAGGTTTGTGGTTGATATTGAAAATACCTTAACTCCTCCGCCAGAACCGTA<br>A |
| *Ti*TabS (MK840853.1) | **ATG**GCTTCTTCAACTGAAAGCTCTGATGAGATTATTTTTGATCTTCCTCCATACATTAGAGTCTTT<br>AAGGATGGAAGAGTAGAGAGACTCCACTCCTCACCATATGTTCCACCATCACTAGATGATCCCG<br>CAACCGGCGTATCCTGGAAAGACGTCCCAATTTCATCAGAGGTTTCGGCTAGAATCTACCTCCCA<br>AAGATAAGCCAAAAGGAAAAGGAAAAGCTTCCCATTGTGGTCTATTTCCATGGTGCAGGCTTCT<br>GTCTGGAATCCGCCTACAAGTCATTTTTCCACACTTATGTCAAGCACTTTGCAGCCGAGGCCAAA<br>GCAATTGCAGTTTCGGTTGAGTTCAGGCTCTCCCCAGAGCACCACCTGCCTGCAGCTTATGAAGA<br>TTGCTGGACTGCCCTTCAGTGGGTGGCTTCACATGTAGATGTTGACAACTCCAGCCTCAAGAATG<br>CTATAGATAAAGAGCCTTGGATAATCAACCATGGAGACTTTGACAAGATCTACTTATGGGGTGA<br>CAGTACGGGTGCCAATATTGTGCACAACGTACTCATCAGAGCTGGTAATGAGAGCTTGCATGGC<br>GGAGTGAAAATCGTGGGTGCAATTCTTTATTACCCATATTTCTTGATCAGGACAAGCTCCAGACA<br>GAGCGATTATATGGAGAACGAGTACAGAGCATACTGGAAGCTGGCTTATCCATCTGCTCCAGGT<br>GGGAACGACAACCCGATGATAAACCCCGTAGCTGAGAACGCTCCTGATTTGGCTGGATATGGAT<br>GTTCGAGGCTGCTGGTATCCATGGTGGCAGACGAGGCCAGAGACATAACCCTTCTCTACATCGA<br>GGCAGTGAAGAAGAGTGGGTGGAAAGGTGAATTGGAGGTGGCTGATTTCGAAGGAGATTACTTT<br>GAAATATTCAGCCCAGAAACTGAGACAGGCAAGAACAAGGTCAAACGTTTAACGTCTTTCATCA<br>ACAAGGAGTAA |
| *Ti*CorS (MK840854.1) | **ATG**GCTAATTCAACTGCAAACTCTGATGAGATTGTTTTCGATCTTCATCCATACATCAGAGTCTT<br>TAAAAACGGCAAGGTAGAAAGACTTCACGACACCCCATATGTTCCGCCATCACTTGAAGATCCA<br>GCCACCGGTGTATCCTGGAAAGACGTCCCAATTTCATCCGACGTTTCAGCTAGAGTCTACCTCCC<br>GAAGATCAGCGAAGCGGAAAAGAAAAAGCTCCCCATTTTCGTCTATTTCCATGGTGCAGGCTTC<br>TGTCTGGAATCAGCCTTCAAATCATTTTTCCATACTTATGTTAAGCACGTTGTTGCCGAAACCAA<br>AGCTGTCGGAGTTTCGGTTGAGTACAGACTCGCCCCCGAGCACCCTTTACCTGCGGCTTATGAAG<br>ATTGCTGGACTGCCCTTCAGTGGGTGGCTTCCCATGTTGGTCTTGACAACTCCAGCCTCAAGAAT |

| | |
|---|---|
| | GCTATTGATAAAGAGCCTTGGATAATCAACCATGGCGACCTCAATAAGCTTTACTTGGGTGGTGA<br>CAGTCCTGGTGGAAATATTGTGCACAACGTACTGATTAGAGCTGGTAAGGAGAGCTTGCATGGC<br>GGAGTGAAAATCCGGGGTGCAATTCTTTATTACCCATATTTCTTGATCAGGACAAGCAAAAGAC<br>AGAGTGATTATATGGAGATTGACTATAGAGGCTACTGGAAGTTGGCTTATCCATCTGCTCCTGGC<br>GGCACTGACAACCCAATGATAAACCCTGTAGCTAAGAATGCTCCTGATTTGGCCGGATATGGAT<br>GTTCGAGGCTGCTTGTTTCCATGGTTTCGGACGAGACCAGAGATATAACCCTTCTCTACCTTGAG<br>GCATTGAAGAAGAGTGGGTGGAAAGGTGAATTGGAAGTGGGTGACTACGAAGCACATTTCTTTG<br>ATTTGTTCAGCCCTGAAAATGAAGTTGGCAAGACTTGGATCAAACGTTCAAGCGATTTCATCAAC<br>AAGGAGTAA |
| *Ti*PAS 1<br>(MK840850.1) | ATGTATACTACTGAAGTTCGCAAAGTTTCATATCTTACTGCTTTCTCCTTCTTGTCTCACAATGC<br>CACTGCTTCATTCCTGAATCTTTCATCAGTTGTCTTTCCAAGAAATTTCCAACGGATGAACCCATA<br>TTCAGCGTTCTGCATGATCGCCGTAATGCTTCATATCAATCTGTCCTGGAATCTAATCTTCAGAAT<br>CTTAGATTTCTCAAATCAGCAAAACCATTGGCTATCATCACTCCTCTCCTTTACTCCCATGTCCAA<br>GCTGCTGTTGTCTGTTGCAAACGAGCTGGACTACAAATCAGAATCCGCAGTGGTGGCAATGACT<br>ATGAAGGCTTGTCATATCGTTCTGAGGTTCCATATATCATTCTAGACCTTCAAAATCTTAGGTCA<br>ATCATGGTTGACATTGAAGACAACAGTGCTTGGGTTGAGTCAGGAGCAACTATTGGTGAACTGT<br>ATTACGAGATAGCTGAGAAGAGTCCTGTTCATGCCTTTCCTGCTGGTGTCCATCCGACCGTTGGC<br>GTTGGCGGGCACTTAAGTGGTGGTGGGTTTGGTACTATGCTCAGAAAATATGGACTTGCTGCCGA<br>TAACATCCTTGATGCTCATATTGTTGATGCTGAAGGAAGACTTCTCAATAGGGAATCCATGGGAA<br>CAGATTTGTTTTGGGCCATCAGAGGAGGTGGAGGAGCAAGTTTTGGTGTCATAGTTGCCTGGAA<br>AATCAAACTTGTGCATGTTCCTCCAGTAGTCACAGTATTTGATTTGGCCATGACTTTGGAGGAAG<br>GAGCCATAGATCTTATTCACAAATGGCAAACCGTCGGACCCAATCTCAATGAAGATGCATTTCTT<br>GCCGCTAGTATTATGGCAGATCCATCAAGTGAAAACATAACACTCCTGGCAGGTTTCTTTTCATT<br>GTTCCTTGGTACAGCTGACCAACTCCTGAAAGAAATGGGGGAAAGCTTCCCCGAGCTCGGCCTA<br>CGAAAAGAACATTGTTTGGAGATGAGTTGGATCAAAGCAGCACTGCATTTTTCAGGGTATGAAT<br>CTGGTGAAACACTATATGCACTTAAAAATCGAAAGCCTCCTCAACCGAAGCATTGCATCACAGT<br>AAGGTCAGATTTCATTCAAGAACCTCTATCCCTGCCCGCATTAGACAGGTTGTGGAAGTTTTTAT<br>CAGAGGAAGAGAATCCTCCCATAATTGCCATGCTTCCTCATGGCGGAATGATGAGCAAAATATC<br>AGAAACGGAAATTCCATACCCCTACAGAGAAGGTGTGATATATAGCTTTTGTACGAGTTAATTT<br>GGGATTGTGAGGACGATTCCTTGTCTGAAAGATACGTTAGTGCATTGACAAGGCTCTATGACCAT<br>ATGACTCCTTATGTGCTGAAACATCCAAGAGGTAGTTTTTGAACATGAGAAGCCTCGAAATTGG<br>GAAGAACGATGATTATGGGACAACTTATTCAGAAGCTGAGGAATGGGGATTGAAGTATTTCAAG<br>AATAACTTCAAGAGGTTGGCCATTACCAAAGGTGTAGTTGATCCAGATAACTTCTTTTACTTCGA<br>GCAGAGCATTCCCCCTCTTGCTTCCAAAGACGAACTTTAA |
| *Ti*PAS 2<br>(MK840851.1) | **ATG**GTTGAAGTCTCTAAAGTTCTTTCGTGTTTCTGCTTTCTTCTTCTCCTCTCATTGTCTCATGGCT<br>TGATTCCTGAGGCTTTTATCAGTTGTATTTCCCTGGAATTTGAGTCATATGAATCCATTCTTAGGG<br>TTCTGCATGATCCCGGCAATTCTTCCTACCATTCAGTGCTGCAATCAAGAATTCAAAATCTCAGA<br>TTTCTGAAGTCACCCAAACCATTGGCTATTATCACTCCTCTACTTTACTCCCATGTCCAGGCTGCT<br>ATTGTTTGTAGCAAACGGGTGGGATTACAAATTAGAATCCGCAGTGGAGGAAGTGACTATGAAG<br>GGTTGTCGTATCGTTCTGAGGTTCCATTTATCCTTCTAGACCTCCATAACCTTAGATCCATTGTGG<br>TTGACATTGAAGACAACAGCGCATGGGTTGAGTCAGGAGCAACCATTGGCGAACTGTATTATGA<br>GATTGCCCAGAAAAGTCCTATTCATGCCTTTCCTGCGGGGGTGTATGCAACCGTTGGCGTTGGCG<br>GGCACTTCAGCGGTGGTGGTTTTGGTACGATGCTCAGGAAATATGGACTAGCAGCTGACAATGT<br>GATTGACGCTTATATTGTTGATGCCAAAGGTAGACTTCTGGATAGGGAATCAATGGGAGAAGAT<br>TTGTTTTGGGCCATCAGGGGAGGTGGAGGAGCAAGTTTTGGCGTCATAGTTTCATGGAAGATCA<br>AACTTGTGCATGTTCCTCCAGTGGTCACTGTTTTTGACATGCCCAAGACTTTAGAGCAAGGAGCC<br>TTGGATCTTCTGAACAAATGGCAATATATAGGACACAAGCTCAGTGAAGATCTATTCCTTGCTGT<br>AAGTATCATGGCAGATACATCTGGTGGAAATAAAACGCTTATGGCAGGTTTCAACTCGTTGTTTC<br>TCGGCACAGCTGACCAGCTCTTGAAAGAAATGGCAGAAAGCTTCCCCGAGCTAGGGTTGAGAAA<br>AGAACATTGCTCCGAGATGGGTTGGATCAAGGCAGCAATGCATTTTTCTGGATATCCAAGTGTA<br>GAAACCACATCTCCACTTAAAAGTCGAGAACCTCCTCTACCCAAGACTTGCATTGCGACCAAGT<br>CAGACTTCATTCAAGAACCTCTATCCCTGCCCGCATTAGAGAAGTTGTGGAAGTTGTTATGGGAG<br>GAAGAGAATACTCCCATCATTCTCATGCTTCCTCATGGCGGAATGATGAGCAAAATATCAGAAT<br>GGGAACTTCCATTCCCATACAGACAAGATGTGATATACAGCATGATCGATGAAGTAGTATGGGA<br>TTGTGAAGACGATTACTCCTCCGAAGAGCACATTAGTGGACTGAGAAAGCTCTACGATCTTATG<br>ACGCCTTATGTGTCGAAACACCCAAGAGGTACTTTTCTAAACATGAGAAACCTTGACACGGGTA<br>GGAACGGTGATTATTATGGCACAACTTATTCAAAAGCCAAGGAATGGGGATTGAAGTATTTCAA<br>GAACAACTTTCGAAAGTTGGCCATTACCAAGGGTGCAGTTGATCCAGATAACTTCTTTTACTTCG<br>AGCAGAGCATTCCGCCTCTCACTTCACAAGACGAATCATCGTAA |
| *Ti*PAS 3<br>(MK840852.1) | **ATG**TTAGCAGAAGTCTCCAAAGTTCTTTCATGTTTCTACTTTCTTGTTCTACTCTCAACATCGCAT<br>GGCTTGATTCCTGAAGCTTTTATTAATTGTGTTTCCCAGAAATTTGTGTCGGATCAATCTATTTTT<br>AGCATTCTGCATGGTCCCGGAAATTCTTCCTATCATTCTGTGCTGCAATCTAGAATTCAAAATCTT<br>AGATTTCTCAAGTCACCCAAACCACTGGCTATTATCACTCCGCTACTTTACTCTCATGTCCAAGC<br>CGCTGTCGTTTGTAGCAAACAGGTCGGACTACAAATTAGAATCCGCAGTGGAGGGGCCGACTAT<br>GAAGGGTTGTCATATCGTTCTGAGGTTCCTTTTGTCCTACTAGACCTCCAAAATCTTAGATCCATT<br>GCGGTTGACATTGAGGACAACAGCGCTTGGGTTGAGTCAGGAGCAACCCTTGGCGAACTGTATT<br>ATGAGATCGCTGAGAAAAGTCCGATTCATGCCTTTCCTGCAGGGGCCTCTGCAACTGTAGGCGTT |

| | |
|---|---|
| | GGCGGGCACTTCAGTTGTGGTGGTTTTGGTACGATGCTCAGAAAATATGGACTGGCATCTGACA<br>ATGTAATTGATGCTTATATCGTTGATGCCACAGGTAGACTTCTGGATAAGGAATCAATGGGAGA<br>AGATTTGTTTTGGGCCATCAGAGGAGGTGGAGGAGAAAGTTTTGGCGTCATAGTTTCGTGGAAG<br>ATCAAACTTGTGTATGTTCCTCCAGTGGTCACTGTTTTTGACTTGCCCAAGACTTTGGAGCAGGG<br>AGCCTTAGATCTTCTTAACAAATGGCAGTATATAGGATACAAGCAAAGTGAAGATCTATTCCTTG<br>CTGTAAGTATCATGGCAGATACATCTGCTGGAAATAAAACACTTATGGCAGGTTTCACCTCGTTG<br>TTTCTGGGTACAGCTGACCAGCTTCTCAAGGAAATGCATGAGAGCTTCCCAGAGCTTGGCTTGAG<br>GAAAGAACATTGCTCCGAGATGAGTTGGATCAAGGCAGCAATGCATTTTTCTGGATATCCAAGT<br>GCAGAAACCATATCTGCACTGAAAAATCAAGATCCTCCTCTACCCAAGACTTGCATTGCGACCA<br>AGTCAGACTTCATTCAAGAACCTCTACCCCTGGCAGCATTAGAGAAGTTGTGGAAGTTCTCATCG<br>GACGAAGAGAATACTCCCATAATTCTCATGCTTCCTCATGGCGGAATGATGAGCAAAATATCAG<br>ATTCTGAAACTCCATTCCCATACAGACAAGGTGTGATCTACAGCATGATCGAAGAAGTAGTTTG<br>GGATTGTCAAGACGATTACTCCTCCGAAGAGCACGTTAGTGGACTGAGAAGGCTGTATGACCTT<br>ATGGCGCCTTATGTGTCGAAACAGCCAAGAGGTACTTTTCGAACCATCAGAAACCTCGATACCG<br>GTAGAAATAATGATTCCGGCACAACTTATTCAGCGGCAAAGGAATGGGGTTTGAAGTATTTCAA<br>GAACAACTTTCGAAAGCTAGCCATTACCAAGGGTGCAGTTGATCCAGAAAACTTCTTTTATAATG<br>AACAAAGCATTCCGCCTCTAACTTTACATGATGAATTGTAA |
| *Cr*PAS<br>(MH213134.1) | **ATG**ATAAAAAAAGTCCCAATAGTTCTTTCAATTTTCTGCTTTCTTCTTCTACTCTCATCATCCCAT<br>GGCTCAATTCCTGAAGCTTTTCTCAATTGTATTTCCAATAAATTTTCATTAGATGTATCCATTTTA<br>AACATTCTTCATGTTCCCAGCAATTCTTCCTATGATTCTGTTCTCAAATCTACTATCCAAAATCCA<br>AGATTCCTCAAATCACCCAAGCCCTTAGCTATAATCACCCCAGTACTTCACTCCCATGTCCAATC<br>TGCTGTTATCTGTACCAAACAAGCCGGTTTACAAATTAGAATCCGAAGCGGAGGAGCTGATTAC<br>GAGGGCTTATCCTATCGTTCTGAGGTTCCCTTTATTCTGCTAGATCTCCAGAATCTTCGATCAATT<br>TCCGTTGATATTGAAGACAACAGCGCTTGGGTCGAATCAGGAGCAACAATTGGTGAATTCTATC<br>ATGAGATAGCTCAGAACAGCCCTGTTCATGCGTTTCCAGCTGGGGTCTCTTCCTCTGTTGGAATT<br>GGCGGCCATTTGAGTAGCGGCGGTTTTGGTACATTGCTTCGGAAATATGGATTAGCAGCCGATA<br>ATATAATCGATGCAAAAATTGTTGATGCCAGAGGCAGAATTCTTGATAGGGAATCAATGGGAGA<br>AGATCTATTTTGGGCTATTAGAGGAGGAGGAGGAGCTAGTTTTGGTGTTATAGTTTCTTGGAAGG<br>TTAAACTTGTAAAAGTCCCTCCGATGGTAACTGTTTTCATCTTGTCCAAGACTTATGAAGAAGGA<br>GGTTTAGATCTTCTACACAAATGGCAATATATAGAACACAAACTCCCTGAAGATTTATTCCTTGC<br>TGTAAGCATCATGGATGATTCATCTAGTGGAAATAAAACACTTATGGCAGGTTTTATGTCTCTGT<br>TTCTTGGAAAAACAGAGGACCTTCTGAAAGTAATGGCGGAAAATTTCCCACAACTTGGATTGAA<br>AAAGGAAGATTGCTTAGAAATGAATTGGATTGATGCAGCAATGTATTTTTCAGGACACCCAATT<br>GGAGAATCCCGATCTGTGCTTAAAAACCGAGAATCTCATCTTCCAAAGACATGCGTTTCGATCAA<br>ATCAGACTTTATTCAAGAACCACAATCCATGGATGCATTGGAAAAGTTATGGAAGTTTTGTAGG<br>GAAGAAGAAAATAGTCCCATAATACTGATGCTTCCACTGGGGGGAATGATGAGTAAAATATCAG<br>AATCAGAAATCCCATTTCCTTACAGAAAAGATGTGATTTACAGTATGATATACGAAATAGTTTGG<br>AATTGTGAAGACGATGAATCATCGGAAGAATATATCGATGGATTGGGAAGGCTTGAGGAATTAA<br>TGACTCCATATGTGAAACAACCAAGAGGTTCTTGGTTCAGCACCAGAAACCTTTATACCGGTAA<br>AAATAAAGGTCCAGGAACAACTTATTCCAAAGCTAAAGAATGGGGATTTCGGTATTTTAATAAT<br>AATTTCAAAAAGTTGGCCCTTATCAAAGGACAAGTTGATCCAGAAAACTTCTTCTACTATGAACA<br>AAGCATTCCCCCTCTCCATTTACAAGTCGAACTTTGA |
| p19-TBSV<br>(M21958.1) | **ATG**GAACGAGCTATACAAGGAAACGATGCTAGGGAACAAGCTTATGGTGAACGTTGGAATGGA<br>GGATCAGGAAGTTCCACTTCTCCCTTCAAACTTCCTGACGAAAGTCCGAGTTGGACTGAGTGGCG<br>GCTACATAACGATGAGACTATTTCGAATCAAGATAATCCCCTTGGTTTCAAGGAAAGCTGGGGTT<br>TCGGGAAAGTTGTATTTAAGAGATATCTCAGATACGACGGGACGGAAACTTCACTGCACAGAGT<br>CCTTGGATCTTGGACGGGAGATTCGGTTAACTATGCAGCATCTCGATTTCTCGGTTTCGACCAGA<br>TCGGATGTACCTATAGTATTCGGTTTCGAGGAGTTAGTGTCACCATTTCTGGAGGGTCGCGAACT<br>CTTCAGCATCTCAGTGAAATGGCAATTCGGTCTAAGCAAGAACTGCTACAGCTTACCCCAGTCA<br>AAGTGGAAAGTGATGTATCAAGAGGATGCCCTGAAGGTGTTGAAACCTTCGAAGAAGAAAGCG<br>AGTAA |

Notes

1. Start codon is highlighted in **bold**.
2. Stop codon is underlined.

**Table S2.** Primers used in this study. Primer sequences. Primer sequences with homology to the destination plasmid for DNA assembly are in **bold**.

| Gene | Plasmid | Primer direction | Sequence (5‘-3‘) |
|---|---|---|---|
| *Nicotiana benthamiana* transient expression | | | |
| *Cr*PAS | 3Ω1 | Forward | **TTTATGAATTTTGCAGCTCG**ATGATAAAAAAAGTCCCAATAGTTCTTT |
| | | Reverse | **GACAACCACAACAAGCACCG**TCAAAGTTCGACTTGTAAATGGA |
| *Ti*PAS 1 | 3Ω1 | Forward | **TTTATGAATTTTGCAGCTCG**ATGTATACTACTGAAGTTCGCAAAGTTT |
| | | Reverse | **GACAACCACAACAAGCACCG**TTAAAGTTCGTCTTTGGAAGCAAGA |
| *Ti*PAS 2 | 3Ω1 | Forward | **TTTATGAATTTTGCAGCTCG**ATGGTTGAAGTCTCTAAAGTTCTTTCG |
| | | Reverse | **GACAACCACAACAAGCACCG**TTACGATGATTCGTCTTGTGAAG |
| *Ti*PAS 3 | 3Ω1 | Forward | **TTTATGAATTTTGCAGCTCG**ATGTTAGCAGAAGTCTCCAAAGTTC |
| | | Reverse | **GACAACCACAACAAGCACCG**TTACAATTCATCATGTAAAGTTAGAG |
| *Ti*DPAS 1 | 3Ω1 | Forward | **TTTATGAATTTTGCAGCTCG**ATGGCTGTAAAATCACCTGAAGC |
| | | Reverse | **GACAACCACAACAAGCACCG**CTATTCCGGTGGAGTTAGTGTG |
| *Ti*DPAS 2 | 3Ω1 | Forward | **TTTATGAATTTTGCAGCTCG**ATGGCAGGAAAATCACCAGAAGA |
| | | Reverse | **GACAACCACAACAAGCACCG**TTACGGTTCTGGCGGAGGAG |
| *Ti*TabS | 3Ω1 | Forward | **TTTATGAATTTTGCAGCTCG**ATGGCTTCTTCAACTGAAAGCTCTG |
| | | Reverse | **GACAACCACAACAAGCACCG**TTAGTCCTTGTTGATGAAAGACGTTAG |
| *Ti*CorS | 3Ω1 | Forward | **TTTATGAATTTTGCAGCTCG**ATGGCTAATTCAACTGCGAACTCT |
| | | Reverse | **GACAACCACAACAAGCACCG**TTACTCCTTGTTGATGAAATCGCTTG |
| *Cr*PAS-C-6xHis | 3Ω1 | Forward | **TTTATGAATTTTGCAGCTCG**ATGATAAAAAAAGTCCCAATAGTTCTTT |
| | | Reverse | **GACAACCACAACAAGCACCG**TCA*GTGATGGTGATGGTGATG*AGAAGAACCAGAACCAAGTTCGACTTGTAAATGGA |
| *Ti*PAS 1-C-6xHis | 3Ω1 | Forward | **TTTATGAATTTTGCAGCTCG**ATGTATACTACTGAAGTTCGCAAAGTTT |
| | | Reverse | **GACAACCACAACAAGCACCG**TCA*GTGATGGTGATGGTGATG*AGAAGAACCAGAACCAAGTTCGTCTTTGGAAGCAAGA |
| *Ti*PAS 2-C-6xHis | 3Ω1 | Forward | **TTTATGAATTTTGCAGCTCG**ATGGTTGAAGTCTCTAAAGTTCTTTCG |
| | | Reverse | **GACAACCACAACAAGCACCG**TCA*GTGATGGTGATGGTGATG*AGAAGAACCAGAACCCGATGATTCGTCTTGTGAAG |
| *Ti*PAS 3-C-6xHis | 3Ω1 | Forward | **TTTATGAATTTTGCAGCTCG**ATGTTAGCAGAAGTCTCCAAAGTTC |
| | | Reverse | **GACAACCACAACAAGCACCG**TCA*GTGATGGTGATGGTGATG*AGAAGAACCAGAACCCAATTCATCATGTAAAGTTAGAG |
| p19-TBSV | 3Ω1 | Forward | **TTTATGAATTTTGCAGCTCG**ATGGAACGAGCTATACAAGGAAA |
| | | Reverse | **GACAACCACAACAAGCACCG**TTACTCGCTTTCTTCTTCGAAGG |
| **Gene** | **Plasmid** | **Primer direction** | **Sequence (5‘-3‘)** |
| *Escherichia coli* (DE3) expression | | | |
| *Ti*DPAS 1 | pOPINF | Forward | **AAGTTCTGTTTCAGGGCCCG**GCTGTAAAATCACCTGAAGC |
| | | Reverse | **ATGGTCTAGAAAGCTTTA**TTCCGGTGGAGTTAGTGTG |
| *Ti*DPAS 2 | pOPINF | Forward | **AAGTTCTGTTTCAGGGCCCG**GCAGGAAAATCACCAGAAGA |
| | | Reverse | **ATGGTCTAGAAAGCTTTA**CGGTTCTGGCGGAGGAG |
| *Ti*TabS | pOPINF | Forward | **AAGTTCTGTTTCAGGGCCCG**GCTTCTTCAACTGAAAGCTCTG |
| | | Reverse | **ATGGTCTAGAAAGCTTTA**GTCCTTGTTGATGAAAGACGTTAG |
| *Ti*CorS | pOPINF | Forward | **AAGTTCTGTTTCAGGGCCCG**GCTAATTCAACTGCGAACTCT |
| | | Reverse | **ATGGTCTAGAAAGCTTTA**CTCCTTGTTGATGAAATCGCTTG |
| **Gene** | **Plasmid** | **Primer direction** | **Sequence (5‘-3‘)** |
| Primers used for sequencing plasmid constructs | | | |
| | 3Ω1 | Forward | GATGAAAAAGCCCTAAAATTGGAG |
| | | Reverse | ATTATTCACAAATGAGAAACAGAATGG |
| | pOPINF | Forward | TAATACGACTCACTATAGGG |
| | | Reverse | TAGCCAGAAGTCAGATGCT |

Notes:

1. GSGSS(Gly-Ser-Gly-Ser-Ser)-linker: sequence underlined.
2. C-terminal 6 x His ($His_6$-tag): sequence *italicized.*

**Table S3.** Reaction components for the production of final products using stemmadenine acetate (**6**) as substrate.

<table>
<tr><th>Desired final product</th><th>Buffer used [50 mM]</th><th>Oxidase used [1 µM]</th><th>Cyclase used [5 µM]</th><th>Reductase used [1 µM]</th><th>Cofactors used</th><th>Incubation method</th><th>Optimal pH and enzyme combination</th></tr>
<tr><td>(+)-Ψ-tabersonine (5)</td><td rowspan="4">HEPES<br>pH 7.5<br>or<br>CHES<br>pH 9.5</td><td rowspan="4">CrPAS<br>or<br>TiPAS 1<br>or<br>TiPAS 2<br>or<br>TiPAS 3</td><td rowspan="3">TiCorS</td><td rowspan="4">TiDPAS 1<br>or<br>TiDPAS 2</td><td rowspan="4">FAD<br>[0.2 mM]<br>and<br>NADPH<br>[1 mM]</td><td rowspan="4">Method*</td><td>pH 7.5<br>+ CrPAS<br>+ TiDPAS1<br>+ TiCorS</td></tr>
<tr><td>(+)-Ψ-vincadifformine (9)</td><td>pH 7.5<br>+ TiPAS3<br>+ TiDPAS1<br>+ TiCroS</td></tr>
<tr><td>(–)-coronaridine (4)</td><td>pH 9.5<br>+ TiPAS3<br>+ TiDPAS2<br>+ TiCorS</td></tr>
<tr><td>(–)-tabersonine (2)</td><td>TiTabS</td><td>pH 7.5<br>+ CrPAS<br>+ TiDPAS1<br>+ TiTabS</td></tr>
</table>

Notes:

Method*: Stepwise reaction, incubate stemmadenine acetate (**6**) (substrate) with oxidase and FAD co-factor in reaction buffer at 37 °C, 600 rpm for 15 min. Then add cyclase, reductase, and co-factor NADPH in sequential order, and further incubate at 37 °C, 600 rpm for 60 min. Reactions were quenched after incubation was complete.

**Table S4.** Reaction components for the production of final products using angryline (**1a**) as substrate.

<table>
<tr><th>Desired final product</th><th>Buffer used [50 mM]</th><th>Oxidase used [1 µM]</th><th>Cyclase used [5 µM]</th><th>Reductase used [1 µM]</th><th>Cofactors used</th><th>Incubation method</th><th>Optimal enzyme combination</th></tr>
<tr><td>(+)-Ψ-tabersonine (5)</td><td rowspan="4">CHES<br>pH 9.5</td><td rowspan="3">CrPAS<br>or<br>TiPAS 1<br>or<br>TiPAS 2<br>or<br>TiPAS 3</td><td rowspan="2">TiCorS</td><td rowspan="4">TiDPAS 1<br>or<br>TiDPAS 2</td><td rowspan="3">FAD<br>[0.2 mM]<br>and<br>NADPH<br>[1 mM]</td><td rowspan="4">Method**</td><td>+ CrPAS<br>+ TiDPAS1<br>+ TiCorS</td></tr>
<tr><td>(+)-Ψ-vincadifformine (9)</td><td>+ TiPAS 3<br>+ TiDPAS1<br>+ TiCroS</td></tr>
<tr><td>(–)-tabersonine (2)</td><td>TiTabS</td><td>+ TiTabS<br>[no cofactors required]</td></tr>
<tr><td>(–)-coronaridine (4)</td><td>-</td><td>TiCorS</td><td>NADPH<br>[1 mM]</td><td>+ TiCorS<br>+ TiDPAS2</td></tr>
</table>

Notes:

Method**: One-pot reaction, assemble reaction mixture in buffer and initiate the reaction by the addition of angryline (**1a**). Incubate at 37 °C, 600 rpm for 90 min and quench reaction.

### Supplementary Figures

a.

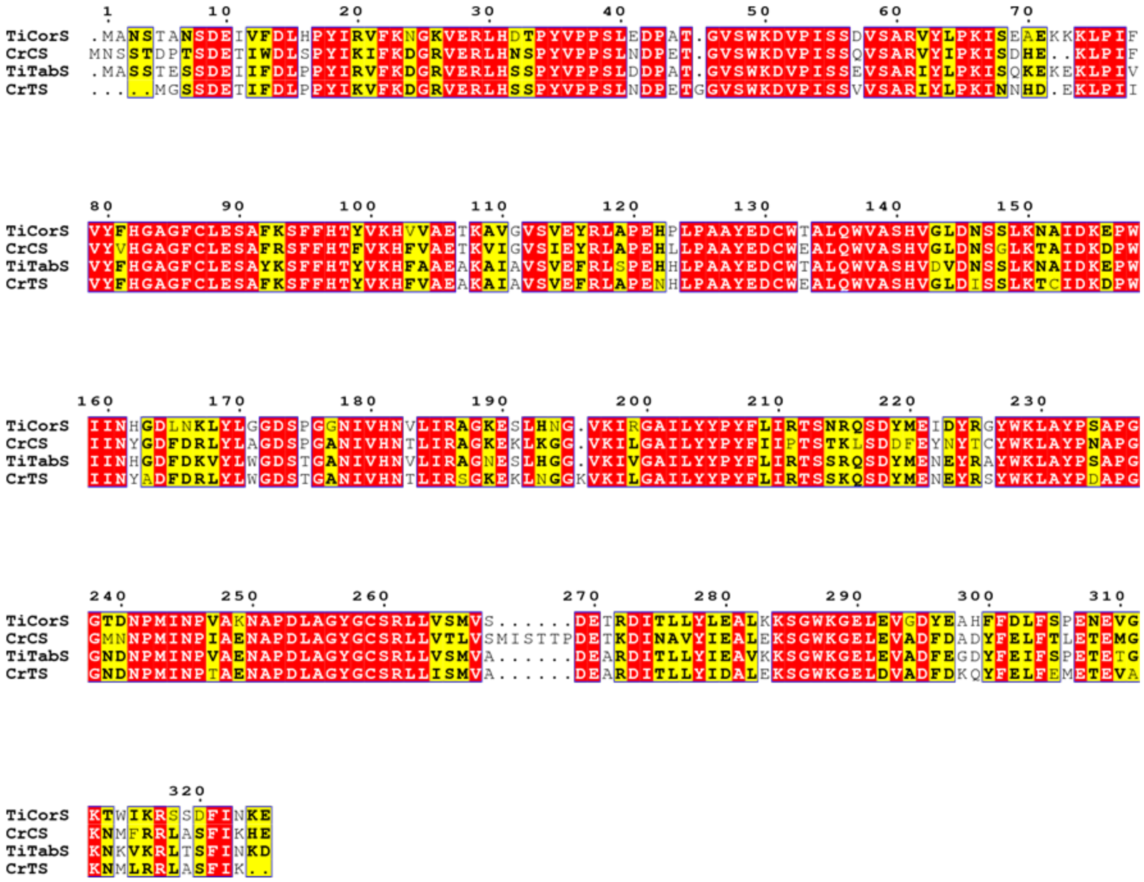

b.

| Enzyme | TiCorS | CrCS | TiTabS | CrTS |
|---|---|---|---|---|
| TiCorS | | 70.8 | 81,0 | 73.3 |
| CrCS | 70.8 | | 71.7 | 77.4 |
| TiTabS | 81,0 | 71.7 | | 81.7 |
| CrTS | 73.3 | 77.4 | 81.7 | |

**Figure S1. Amino-acid alignments of cyclases. a)** Protein sequence alignment of α-β hydrolases was generated using Geneious Prime (version 2022.0.1) Muscle v5 algorithm[10]. Aligned sequences were visualized using ESPript V3[11]. Genbank ID of proteins; *Ti*CorS (QED20593.1)[2,3], *Cr*CS (A0A2P1GIW2.1)[2,12], *Ti*TabS (QED20592.1)[3], *Cr*TS (A0A2P1GIW3.1)[2,12]. **b)** Amino-acid similarity matrix of cyclases. Abbreviations; *Cr*CS, *Catharanthus roseus* catharanthine synthase; *Cr*TS, *Catharanthus roseus* tabersonine synthase; *Ti*TabS, *Tabernanthe iboga* tabersonine synthase; *Ti*CorS, *Tabernanthe iboga* coronaridine synthase.

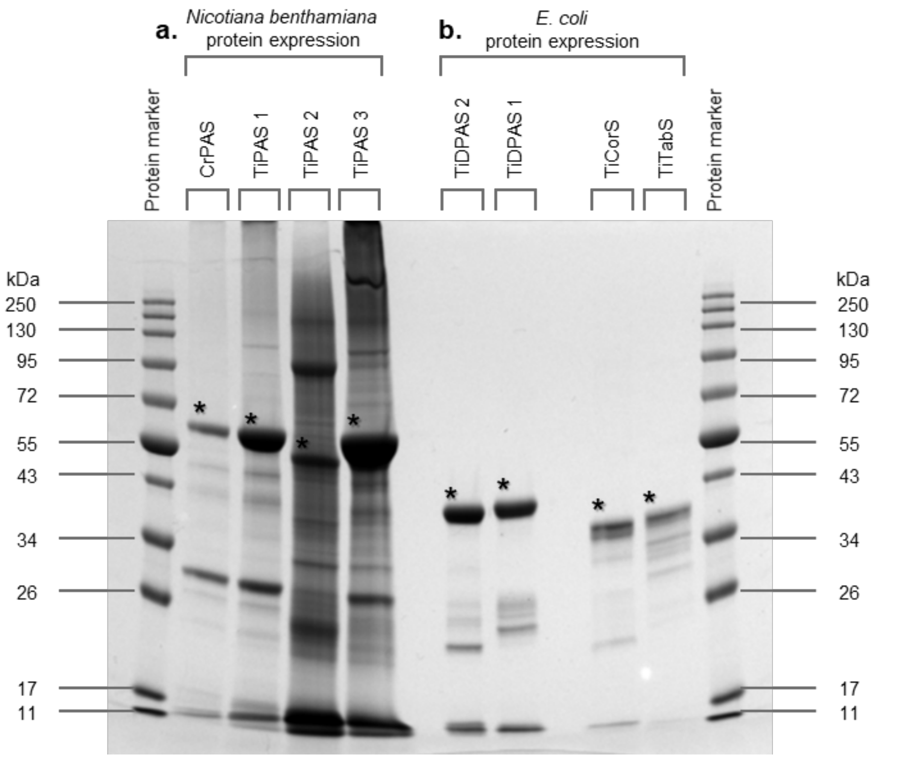


**Figure S2.** SDS-PAGE gel image of proteins used for *in vitro* enzymatic assays. **a**) Proteins with C-terminal $His_6$-tag expressed heterologously in *N. benthamiana*. **b**) Proteins with N-terminal $His_6$-tag expressed heterologously in *E. coli*. Size estimates by ProtParam[13] of the target proteins (*) are as follows: *Cr*PAS – 59647 Da; *Ti*PAS 1 – 59410 Da; *Ti*PAS 2 – 56318 Da; *Ti*PAS 3 – 58751 Da; *Ti*DPAS 2 – 39225 Da; *Ti*DPAS 1 – 40151 Da; *Ti*CorS – 36309 Da; *Ti*TabS – 36502 Da. Abbreviations; *Cr*PAS, *Catharanthus roseus* precondylocarpine acetate synthase; *Ti*PAS1, *Tabernanthe iboga* precondylocarpine acetate synthase 1; *Ti*PAS2, *Tabernanthe iboga* precondylocarpine acetate synthase 2; *Ti*PAS3, *Tabernanthe iboga* precondylocarpine acetate synthase 3; *Ti*DPAS1, *Tabernanthe iboga* dihydro-precondylocarpine acetate synthase 1; *Ti*DPAS2, *Tabernanthe iboga* dihydroprecondylocarpine acetate synthase 2; *Ti*TabS, *Tabernanthe iboga* tabersonine synthase; *Ti*CorS, *Tabernanthe iboga* coronaridine synthase.

**a.**

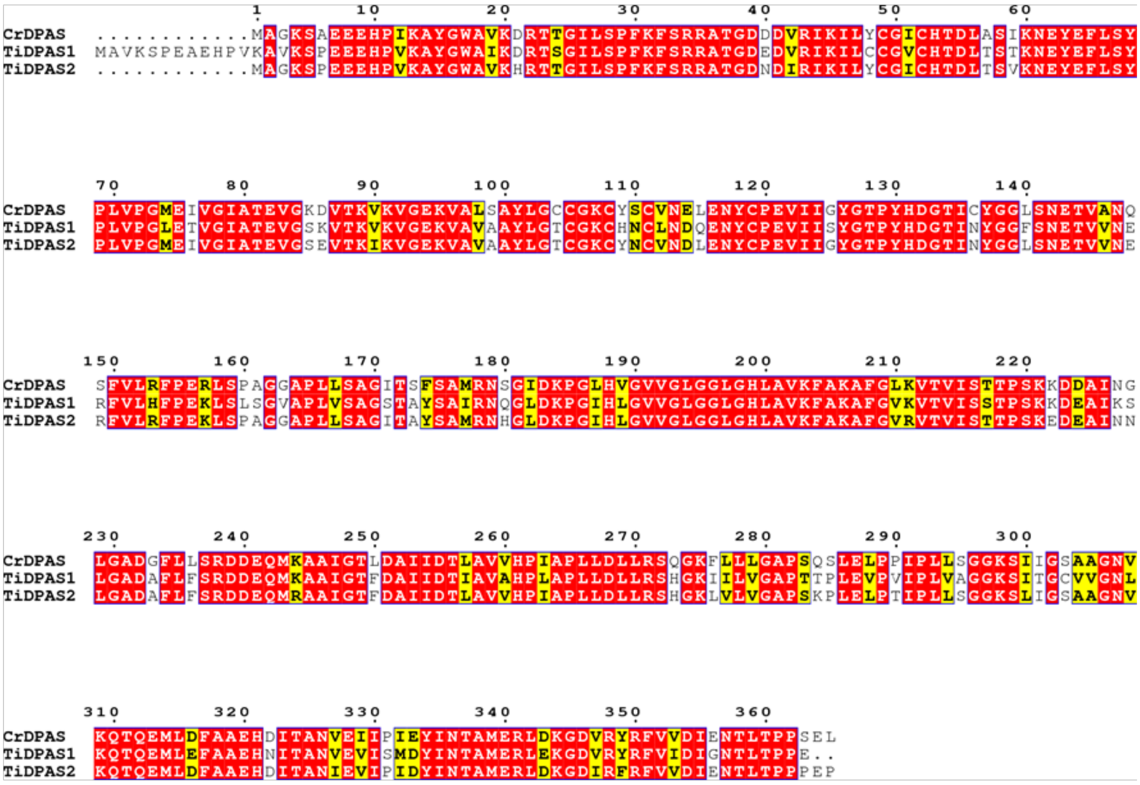


**b.**

| Enzyme | CrDPAS | TiDPAS1 | TiDPAS2 |
|---|---|---|---|
| CrDPAS | | 77.7 | 86.3 |
| TiDPAS1 | 77.7 | | 83.0 |
| TiDPAS2 | 86.3 | 83.0 | |

**Figure S3. Amino-acid alignments of reductases. a)** Protein sequence alignment of medium-chain alcohol dehydrogenase (ADH). Protein sequence alignment was generated using Geneious Prime (version 2022.0.1) Muscle v5 algorithm[10]. Aligned sequences were visualized using ESPript V3[11]. Genbank ID of proteins; *Cr*DPAS (A0A1B1FHP3)[1], *Ti*DPAS1 (A0A5B8XAH0)[3], *Ti*DPAS2 (A0A5B8X8Z0)[3]. **b)** Amino-acid similarity matrix of reductases. Abbreviations; *Cr*DPAS, *Catharanthus roseus* dihydroprecondylocarpine acetate synthase; *Ti*DPAS1, *Tabernanthe iboga* dihydroprecondylocarpine acetate synthase 1; *Ti*DPAS2, *Tabernanthe iboga* dihydroprecondylocarpine acetate synthase 2.

a

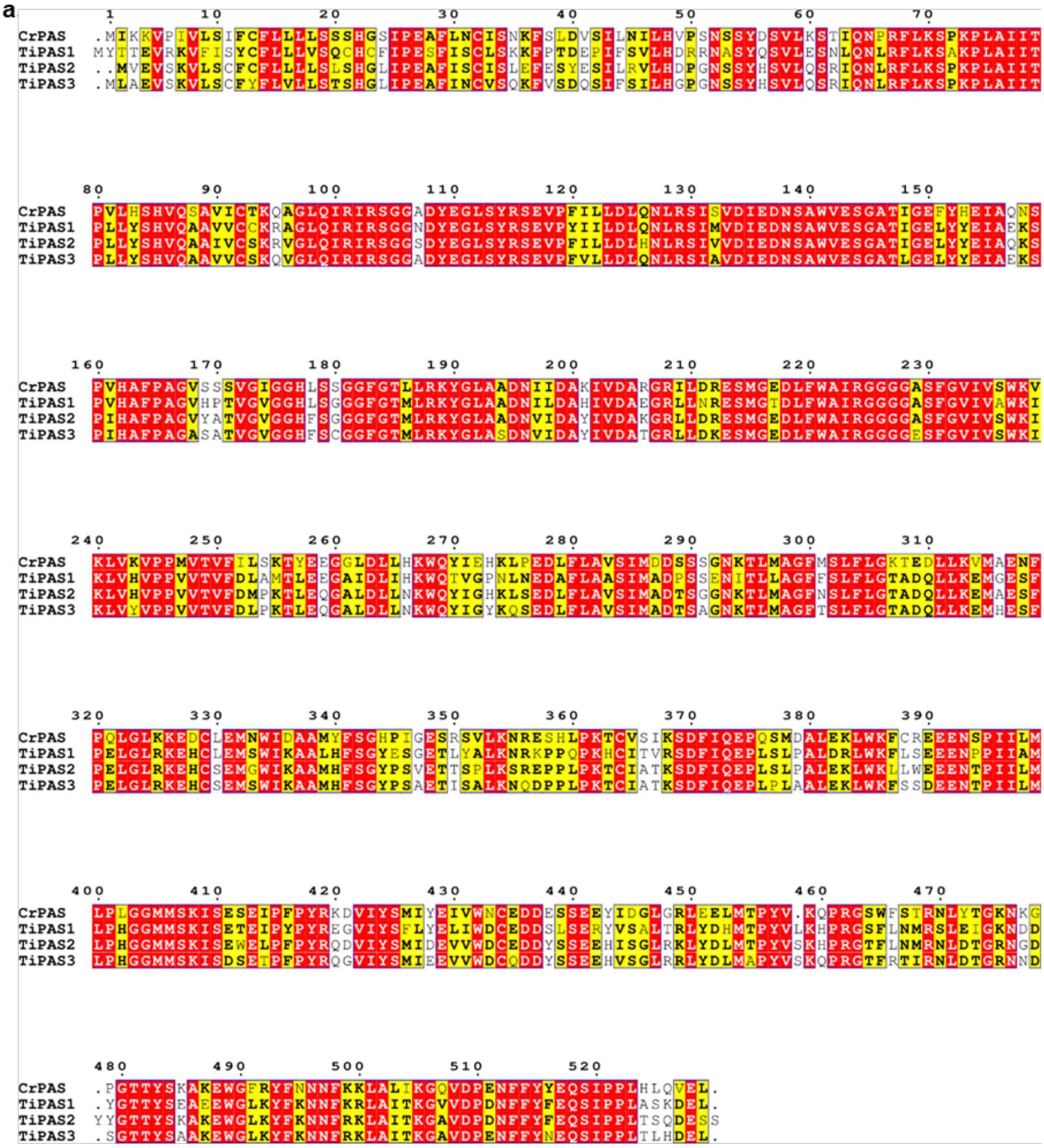

b.

| Enzyme | CrPAS | TiPAS1 | TiPAS2 | TiPAS3 |
|---|---|---|---|---|
| CrPAS | | 67.2 | 73.8 | 72.9 |
| TiPAS1 | 67.2 | | 75.9 | 73.3 |
| TiPAS2 | 73.8 | 75.9 | | 86.1 |
| TiPAS3 | 72.9 | 73.3 | 86.1 | |

**Figure S4. Amino-acid alignments of oxidases. a)** Protein sequence alignment of Flavin-dependent oxidase (PAS). Protein sequence alignment was generated using Geneious Prime (version 2022.0.1) Muscle v5 algorithm[10]. Aligned sequences were visualized using ESPript V3[11]. Genbank ID of proteins; *Cr*PAS (AWJ76616.1)[1], *Ti*PAS1 (QED20589.1)[3], *Ti*PAS2 (QED20590.1)[3], *Ti*PAS3 (QED20591.1)[3]. **b)** Amino-acid similarity matrix of flavin-dependent oxidases. Abbreviations; *Cr*PAS, *Catharanthus roseus* precondylocarpine acetate synthase; *Ti*PAS1, *Tabernanthe iboga* precondylocarpine acetate synthase 1; *Ti*PAS2, *Tabernanthe iboga* precondylocarpine acetate synthase 2; *Ti*PAS3, *Tabernanthe iboga* precondylocarpine acetate synthase 3.

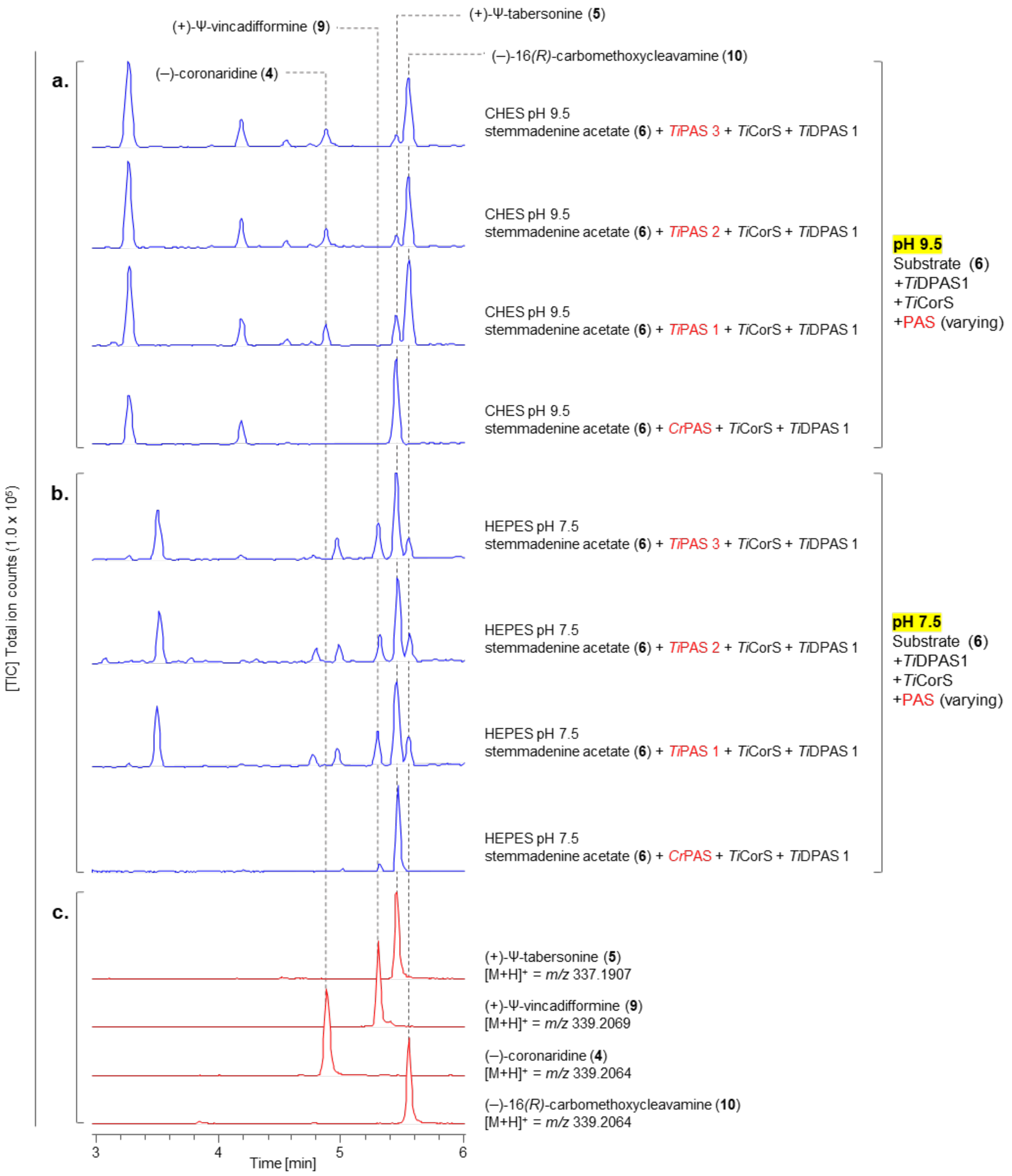


**Figure S5.** Systematic optimization of conditions that lead to production of (+)-Ψ-tabersonine (**5**), (+)-Ψ-vincadifformine (**9**) and (–)-coronaridine (**4**) from stemmadenine acetate (**6**), *Ti*CorS and *Ti*DPAS1. pH conditions and the homologue of PAS that is used in varied. **a)** Conditions at pH 9.5 with *Cr*PAS, *Ti*PAS1, *Ti*PAS2, *Ti*PAS3. **b)** Conditions at pH 7.5 with *Cr*PAS, *Ti*PAS1, *Ti*PAS2, *Ti*PAS3. **c)** Authentic standards of products (TIC).

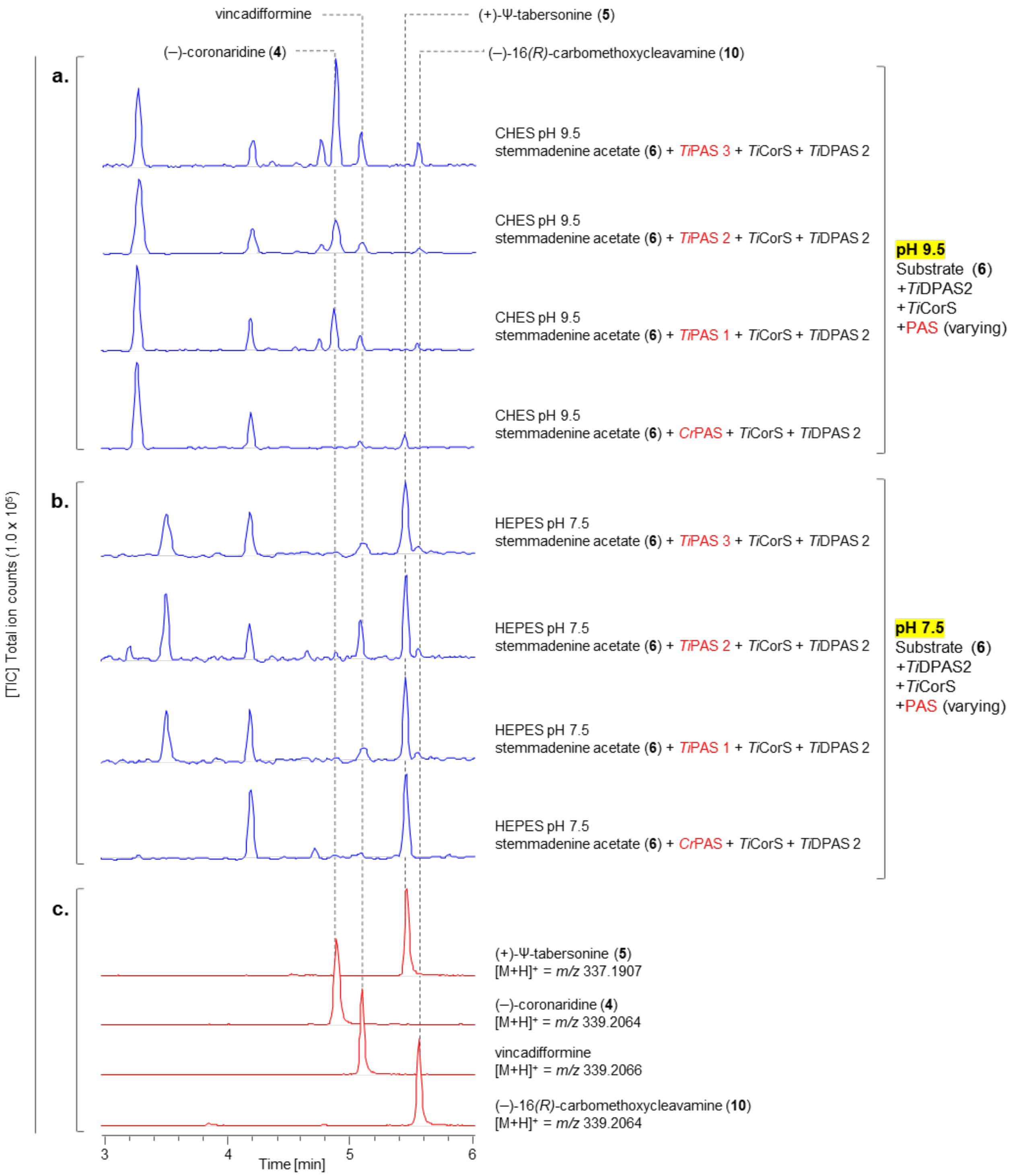


**Figure S6.** Systematic optimization of conditions that lead to production of (+)-Ψ-tabersonine (**5**) and (–)-coronaridine (**4**) from stemmadenine acetate (**6**), *Ti*CorS and *Ti*DPAS2. pH conditions and the homologue of PAS that is used is varied. **a)** Conditions at pH 9.5 with *Cr*PAS, *Ti*PAS1, *Ti*PAS2, *Ti*PAS3. **b)** Conditions at pH 7.5 with *Cr*PAS, *Ti*PAS1, *Ti*PAS2, *Ti*PAS3. **c)** Authentic standards of products (TIC).

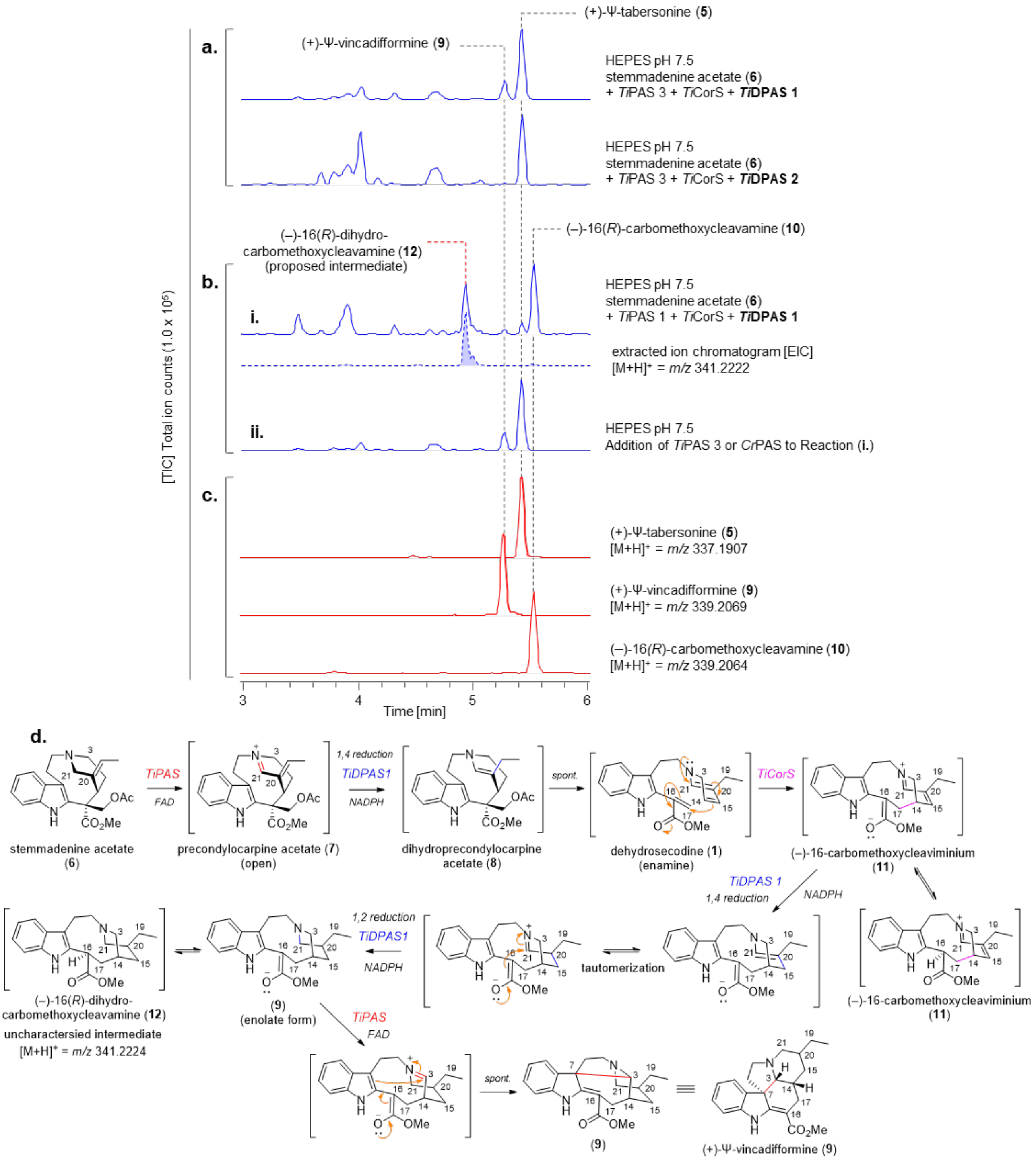


**Figure S7.** ***In vitro* production of the over-reduced product (+)-Ψ-vincadifformine (9). a)** TICs showing the optimized selective production of (+)-Ψ-vincadifformine (**9**) from stemmadenine acetate (**6**) with *Ti*DPAS1 at pH 7.5. **b)** TICs of *in vitro* enzymatic assays showing the uncharacterized over-reduced intermediate, (–)-16(*R*)-dihydrocarbomethoxycleavamine (**12**) (*m/z* = 341.2222) observed (i.) in production of (+)-Ψ-vincadifformine (**9**). Stepwise addition of oxidase (PAS) to reaction (i.) resulted in the formation of desired product (+)-Ψ-vincadifformine (**9**) (ii.) via the proposed intermediate, (–)-16(*R*)-dihydro-carbomethoxycleavamine (**12**) (*m/z* = 341.2224). Production of (+)-Ψ-tabersonine (**5**) was observed in (ii.) because of the oxidation of the intermediate (–)-16(*R*)-carbomethoxycleavamine (**10**) by the same oxidase (PAS). **c)** Authentic standards (TICs) of products. **d)** Proposed mechanism for the production of (+)-Ψ-vincadifformine (**9**) from stemmadenine acetate (**6**) from PAS, *Ti*DPAS1 and *Ti*CorS.

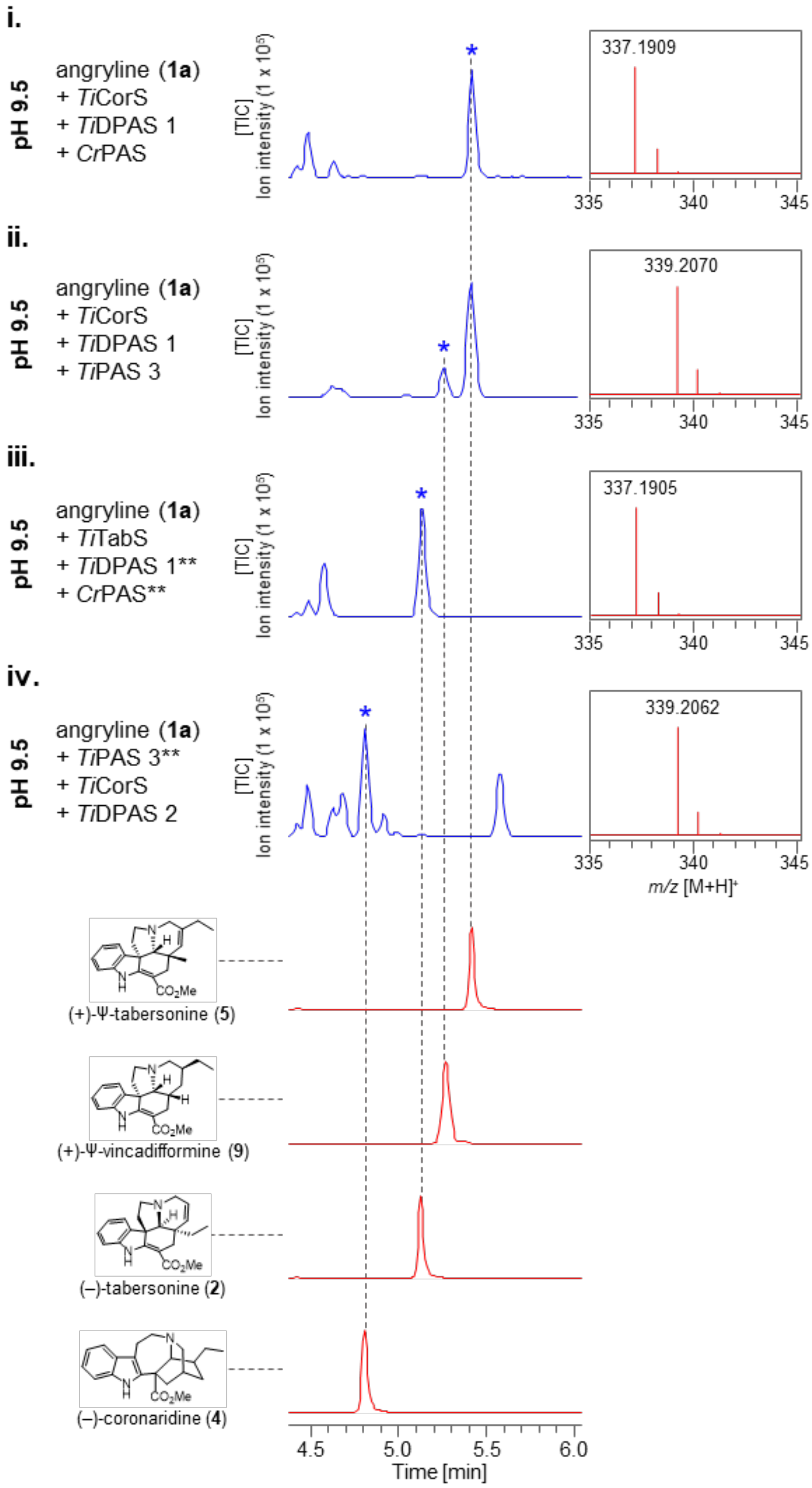


**Figure S8.** ***In vitro*** **coupled enzymatic assays showing optimized production of (+)-Ψ-tabersonine (5), ii. (+)-Ψ-vincadifformine (9), iii. (–)-tabersonine (2) and iv. (–)-coronaridine (4) from angryline (1a).** The enzyme combination and pH conditions have been optimized for each product. *MS, $[M+H]^+$ spectra of observed product. **Enzymes not required for the biosynthesis of desired product (*) but included to show consistency across all enzyme reactions.

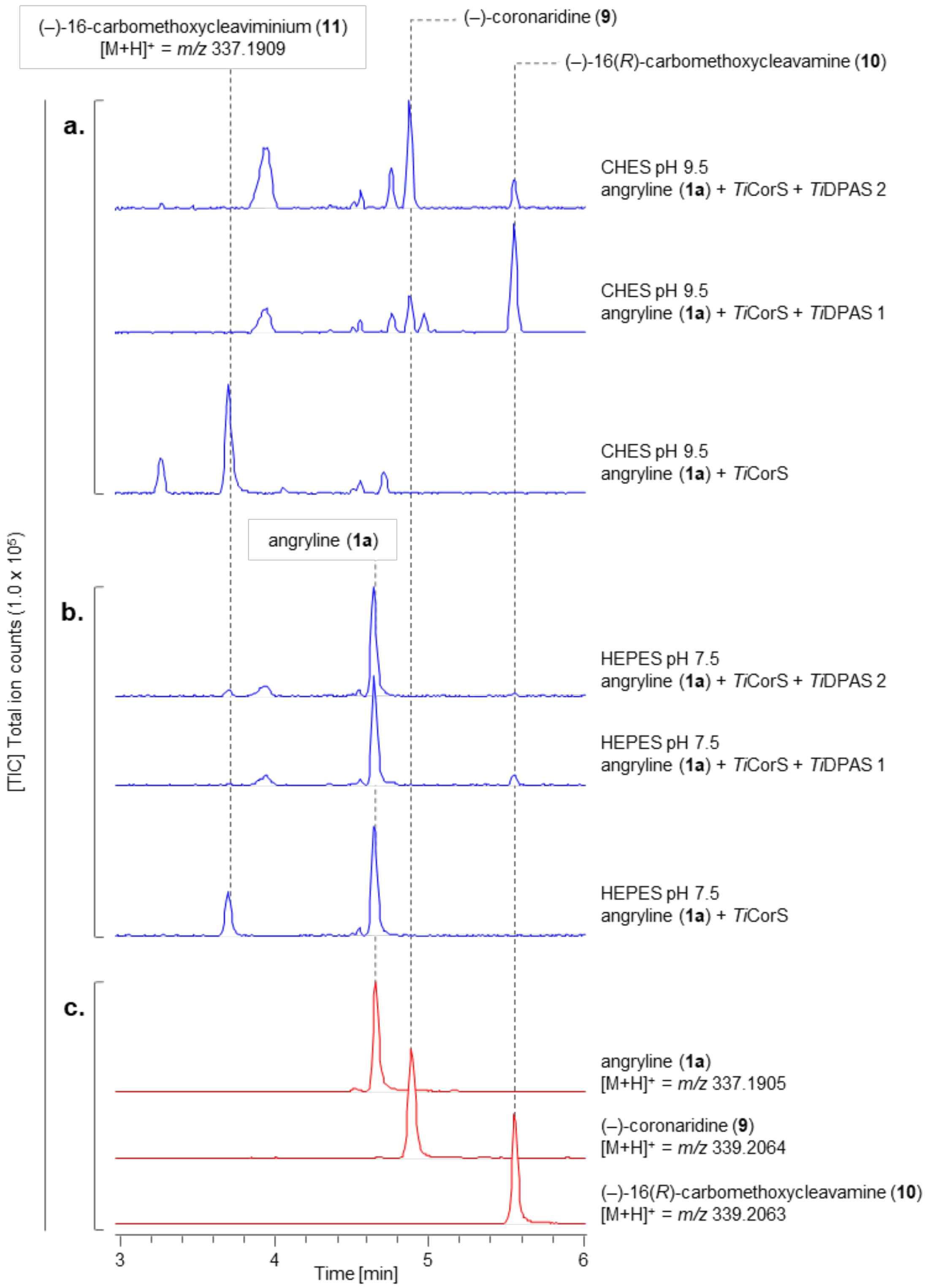


**Figure S9. Systematic optimization of conditions that lead to production of (–)-coronaridine (4) from angryline (1a), with *Ti*CorS and *Ti*DPAS homologues. a)** TICs of *in vitro* reactions for the production of (–)-coronaridine (**4**) from angryline (**1a**) at pH 9.5 with stepwise addition of *Ti*CorS, and *Ti*DPAS1, *Ti*DPAS2 homologues. **b)** TICs of *in vitro* reactions using angryline (**1a**) as substrate with stepwise addition of *Ti*CorS, and *Ti*DPAS1, *Ti*DPAS2 homologues at pH 7.5. **c)** Authentic standards (TICs) of products and substrate.

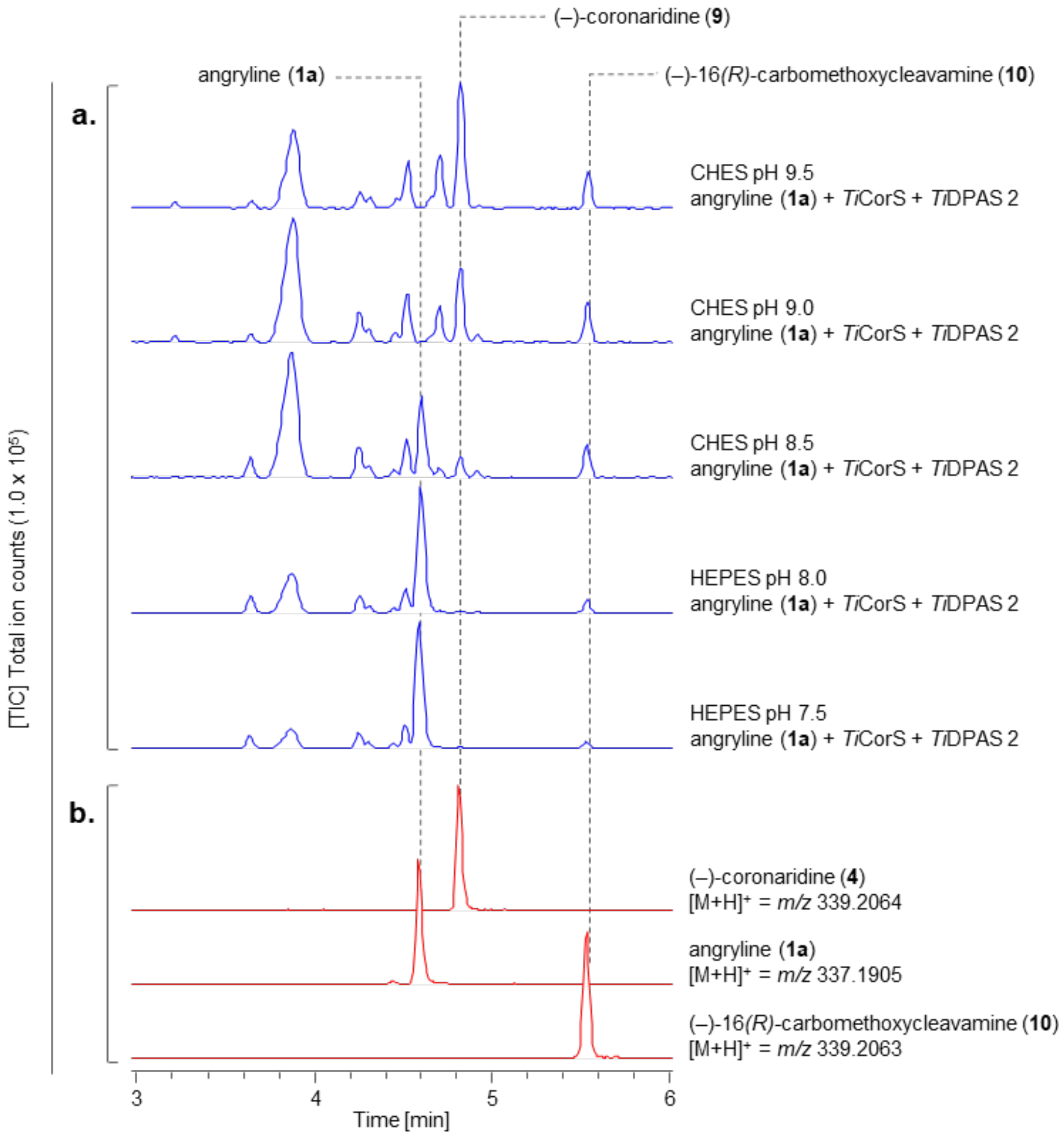


**Figure S10. Systematic optimization of pH conditions that lead to production of (–)-coronaridine (4) from angryline, *Ti*CorS and *Ti*DPAS2. a)** TICs of *in vitro* coupled enzymatic assays for the production of (–)-coronaridine (**4**) from angryline (**1a**) using *Ti*CorS and *Ti*DPAS2 enzymes at pH conditions 9.5, 9.0, 8.5, 8.0, and 7.5. **b)** Authentic standards (TICs) of products and substrate.

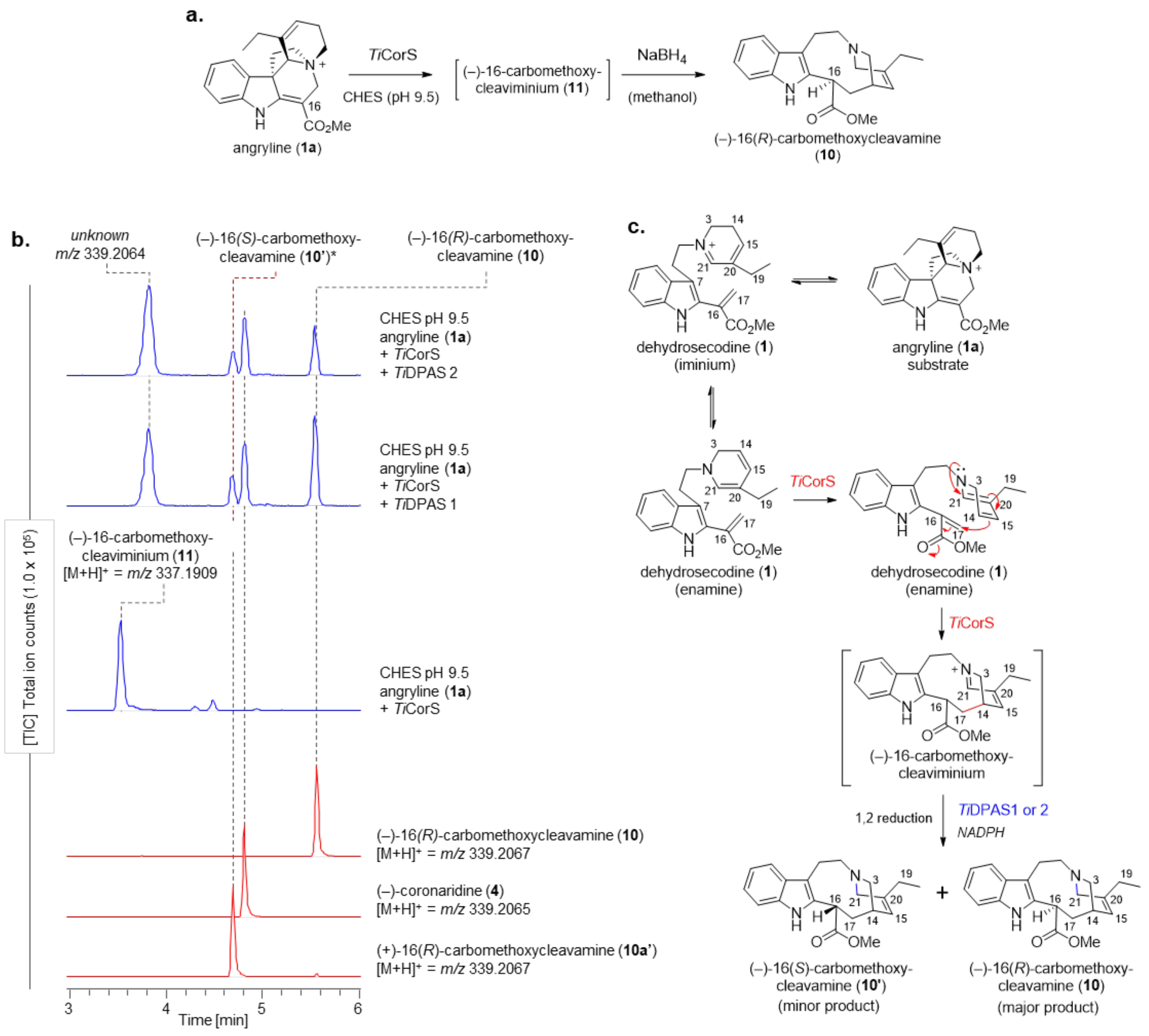


**Figure S11. Generation of (–)-16(*R*)-carbomethoxycleavamine (10) from angryline (1a), by *Ti*CorS and *Ti*DPAS1. a)** Reaction scheme showing enzymatic product of *Ti*CorS, (–)-16-carbomethoxycleaviminium (**11**) chemically reduced to (–)-16(*R*)-carbomethoxycleavamine (**10**) with $NaBH_4$ in methanol. **b)** TICs showing the reaction products using angryline, when *Ti*CorS and either *Ti*DPAS1 or *Ti*DPAS2 is added stepwise at pH 9.5. The peak assigned as (–)-16(*S*)-carbomethoxycleavamine (**10'**) is putative and is based on the retention time and fragmentation spectra of (+)-16(*R*)-carbomethoxycleavamine (**10a**), the enantiomer generated from (+)-catharanthine (**3**) (red-dotted line). Authentic standards (TICs) of the products are shown in red. **c)** Proposed mechanism for formation of (–)-16(*R*)-carbomethoxycleavamine (**10**) and (–)-16(*S*)-carbomethoxycleavamine (**10'**) from angryline (**1a**).

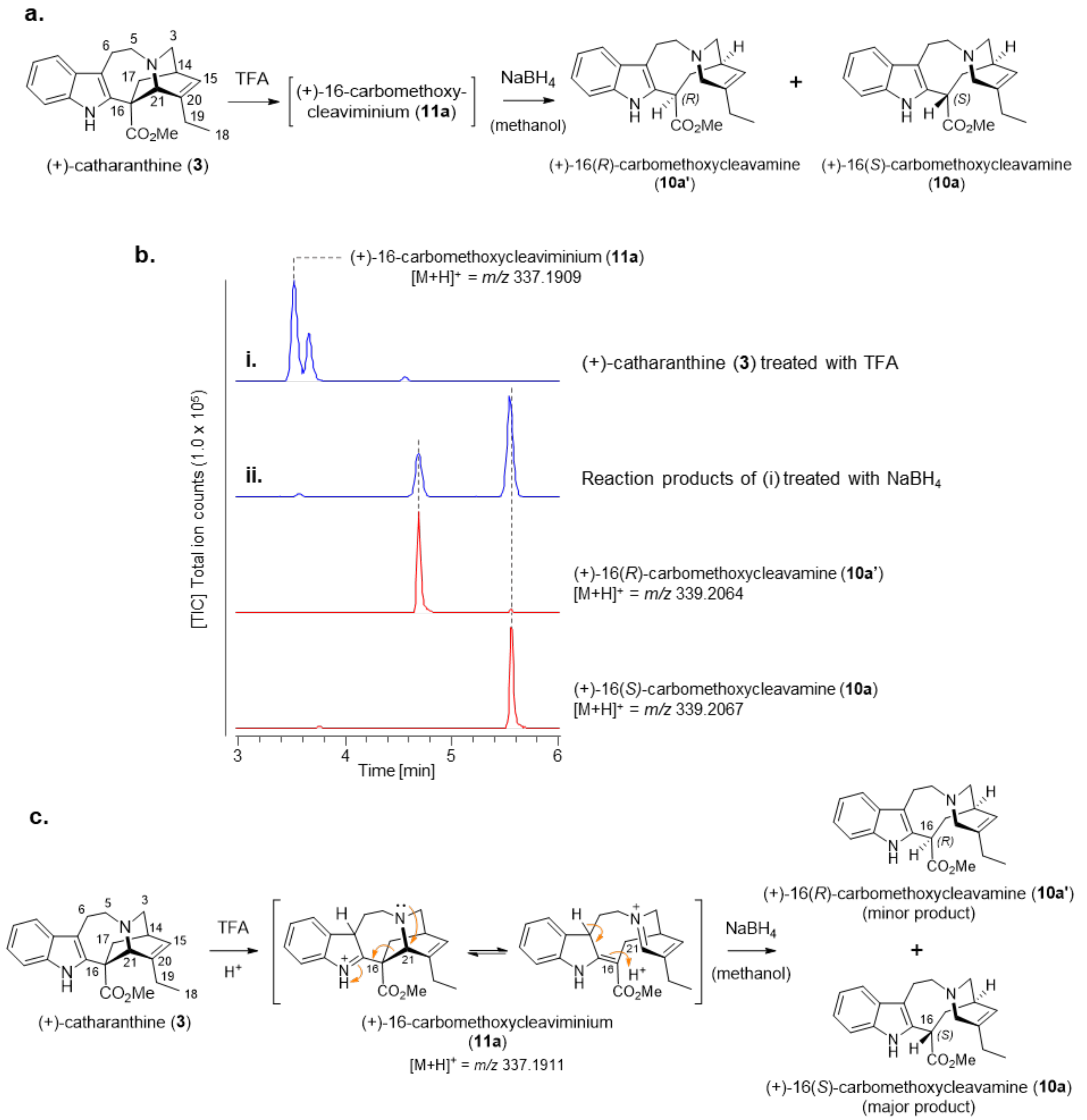


**Figure S12. Generation of 16-carbomethoxycleavamine (10a) from acid-catalyzed opening and reduction of (+)-catharanthine (3). a)** Reaction scheme showing conversion of (+)-catharanthine (**3**) to (+)-16(*R*)-carbomethoxycleavamine (**10a'**) and (+)-16(*S*)-carbomethoxycleavamine (**10a**). **b)** TICs showing the reaction products of acid catalyzed opening and reduction of (+)-catharanthine (**3**). Authentic standards (TICs) of products are shown in red. **c)** Proposed reaction mechanism for the production of (+)-16(*R*)-carbomethoxycleavamine (**10a**) and (+)-16(*S*)-carbomethoxycleavamine (**10a'**) from (+)-catharanthine (**3**).

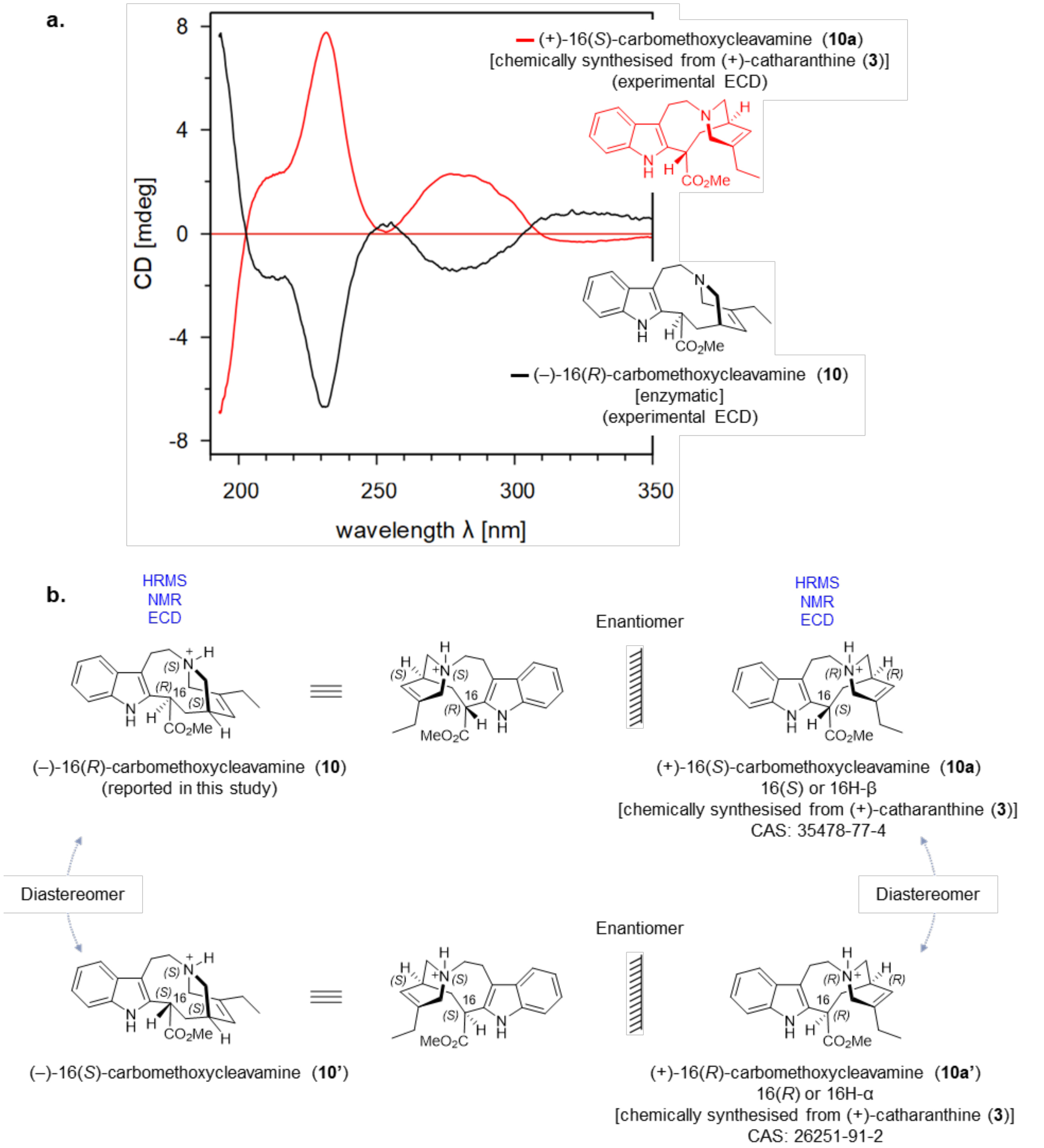


**Figure S13. Stereochemistry of (–)-16(*R*)-carbomethoxycleavamine (. a)** CD spectra of enzymatically prepared and isolated (–)-16(*R*)-carbomethoxycleavamine (**10**) and chemically prepared and isolated (+)-16(*S*)-carbomethoxycleavamine (**10a**).[14] **b)** Stereocenters of 16-carbomethoxycleavamine isomers described in this study.

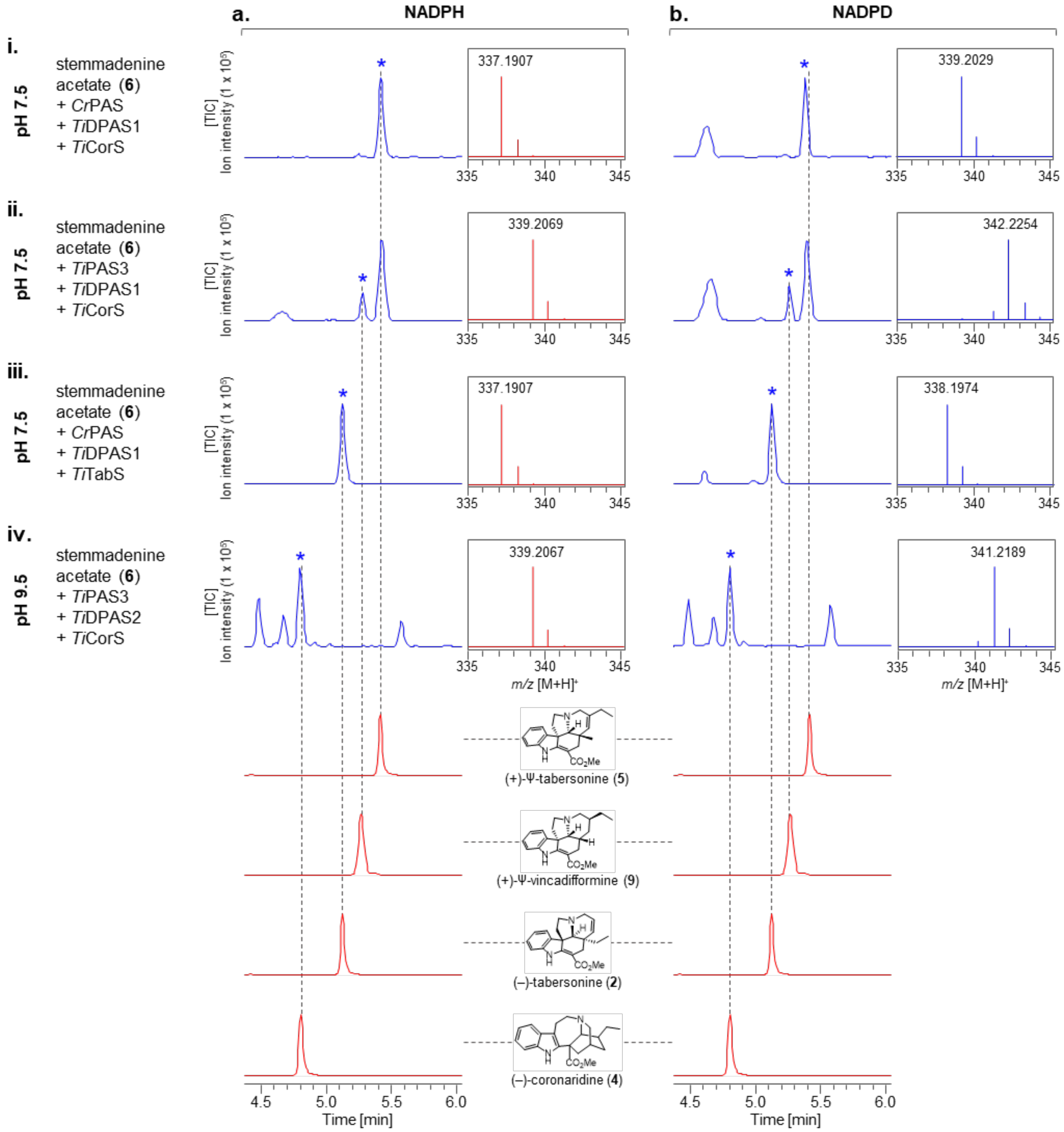


**Figure S14. *In vitro* deuterium labelled enzyme assays using stemmadenine acetate (6) as starting material**. TICs of *in vitro* enzymatic assays showing optimized production of **i.** (+)-Ψ-tabersonine (**5**), **ii.** (+)-Ψ-vincadifformine (**9**), **iii.** (–)-tabersonine (**2**), and **iv.** (–)-coronaridine (**4**) from stemmadenine acetate (**6**). The enzyme combination PAS, DPAS, cyclase (CorS or TabS) and pH conditions have been optimized for each of the respective products. TICs of the authentic standards of the final products are shown in red. **a)** Use of NADPH cofactor. **b)** Use of isotopically labeled NADPD cofactor. *MS, $[M+H]^+$ spectra of observed product.

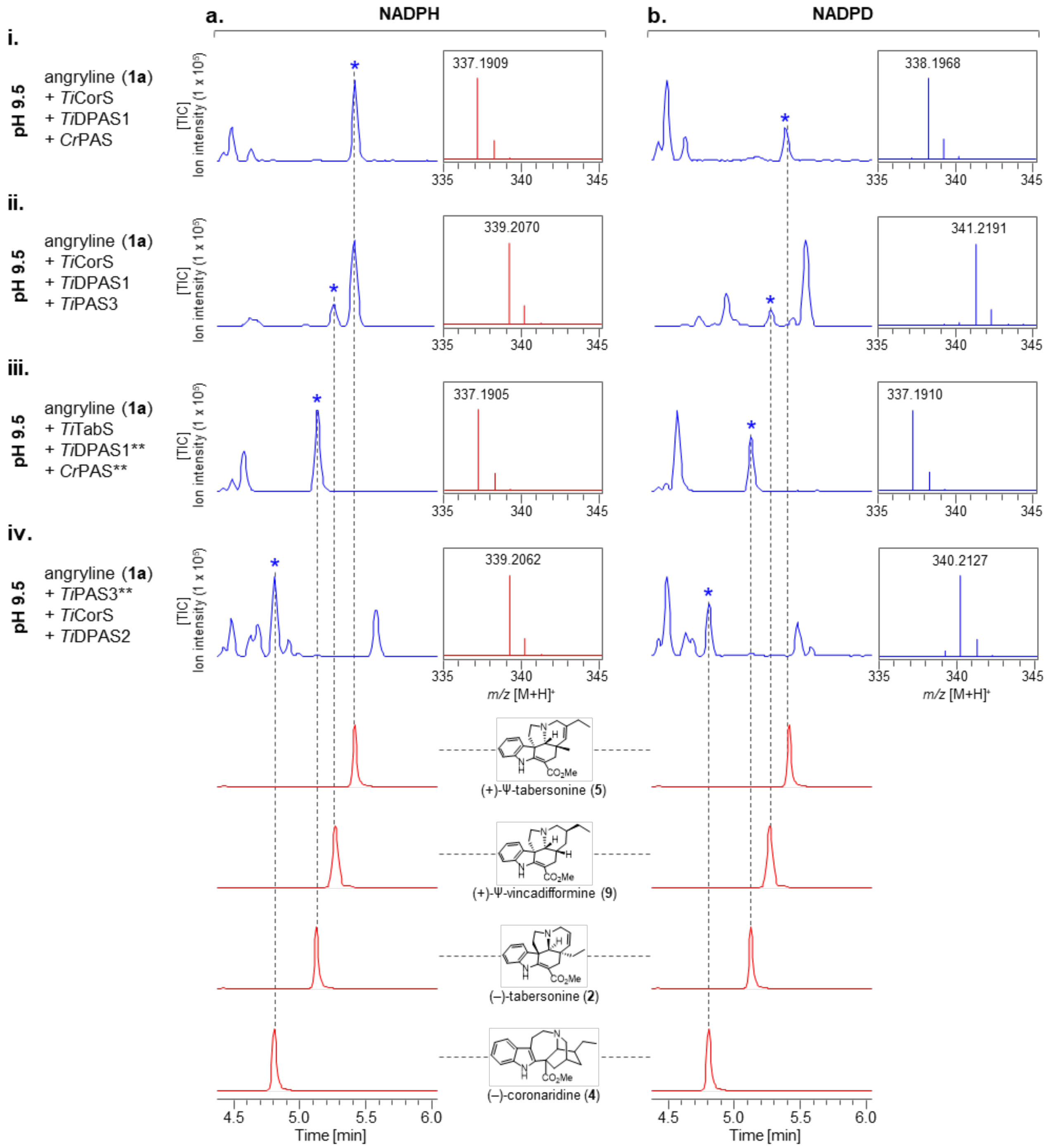


**Figure S15.** ***In vitro* deuterium labelled enzyme assays from angryline (1b)**. TICs of *in vitro* enzymatic assays showing optimized production of **i.** (+)-Ψ-tabersonine (**5**), **ii.** (+)-Ψ-vincadifformine (**9**), **iii.** (–)-tabersonine (**2**), and **iv.** (–)-coronaridine (**4**) from angryline (**1a**). The enzyme combination PAS, DPAS, cyclase (CorS or TabS) and pH conditions have been optimized for each of the respective products. Authentic standards (TICs) of the final products are shown in red. **a)** Use of NADPH cofactor. **b)** Use of isotopically labeled NADPD cofactor. *MS $[M+H]^+$ spectra of the observed product. **Enzymes not required for the biosynthesis of desired product (*) but included to show consistency across all enzyme reactions.

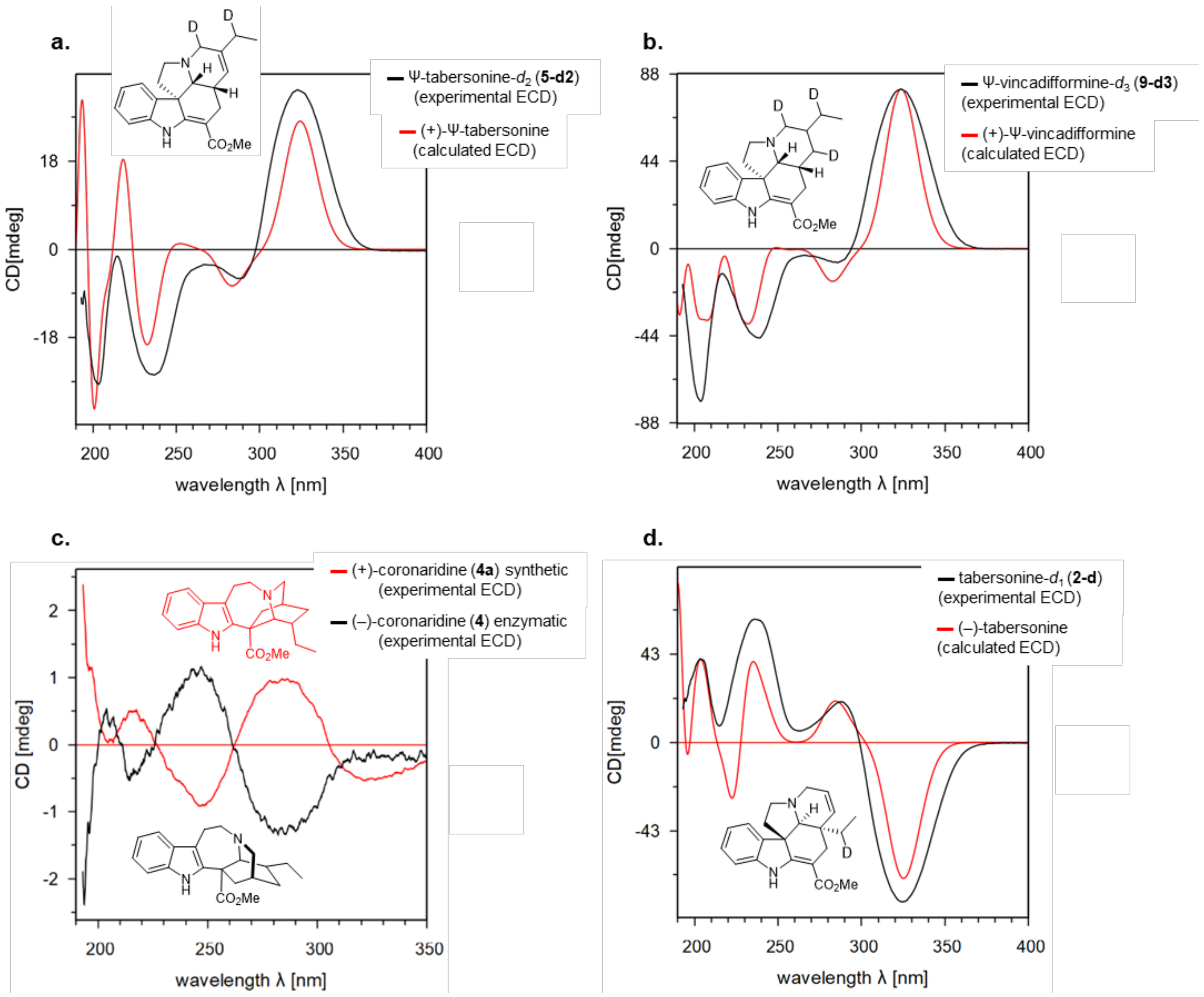


**Figure S16. CD spectra of biosynthetic products. a)** CD spectra of enzymatically prepared and isolated (+)-Ψ-tabersonine-$d_2$ (**5-d2**) (black) alongside computationally calculated spectra (red). **b)** CD spectra of enzymatically prepared and isolated (+)-Ψ-vincadifformine-$d_3$ (**9-d3**) (black) alongside computationally calculated spectra (red). **c)** CD spectra of enzymatically prepared and isolated (–)-coronaridine (**4**) (black) alongside chemically synthesized (+)-coronaridine (**4a**) (red). **d)** CD spectra of enzymatically prepared and isolated (–)-tabersonine-$d_1$ (**2-d**) (black) alongside computationally calculated spectra (red).

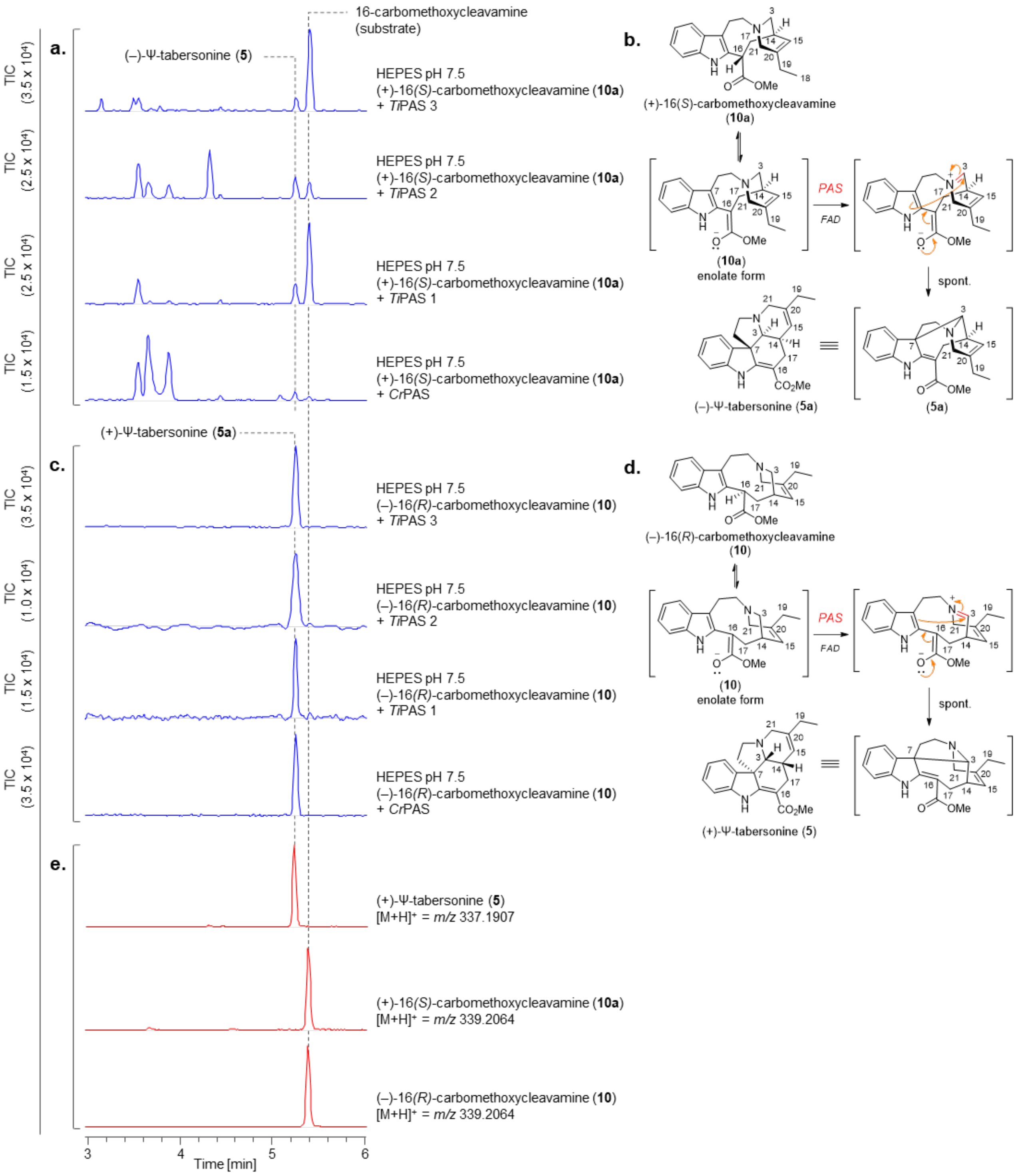


**Figure S17. Production of (–)-Ψ-tabersonine (5a) and (+)-Ψ-tabersonine (5) from 16-carbomethoxycleavine enantiomers. a)** TICs showing the production of (–)-Ψ-tabersonine (**5a**) from (+)-16(*S*)-carbomethoxycleavamine (**10a**) using PAS homologues (oxidases) described in the study. (–)-Ψ-tabersonine (**5a**) could not be structurally characterized and is a putative assignment. **b)** Proposed mechanism for the production of (–)-Ψ-tabersonine (**5a**). **c)** TICs showing the production of (+)-Ψ-tabersonine (**5**) from (–)-16(*R*)-carbomethoxycleavamine (**10**) using PAS homologues (oxidases) described in the study. **d)** Proposed mechanism for the production of (+)-Ψ-tabersonine (**5**). **e)** TICs of authentic standards of standards.

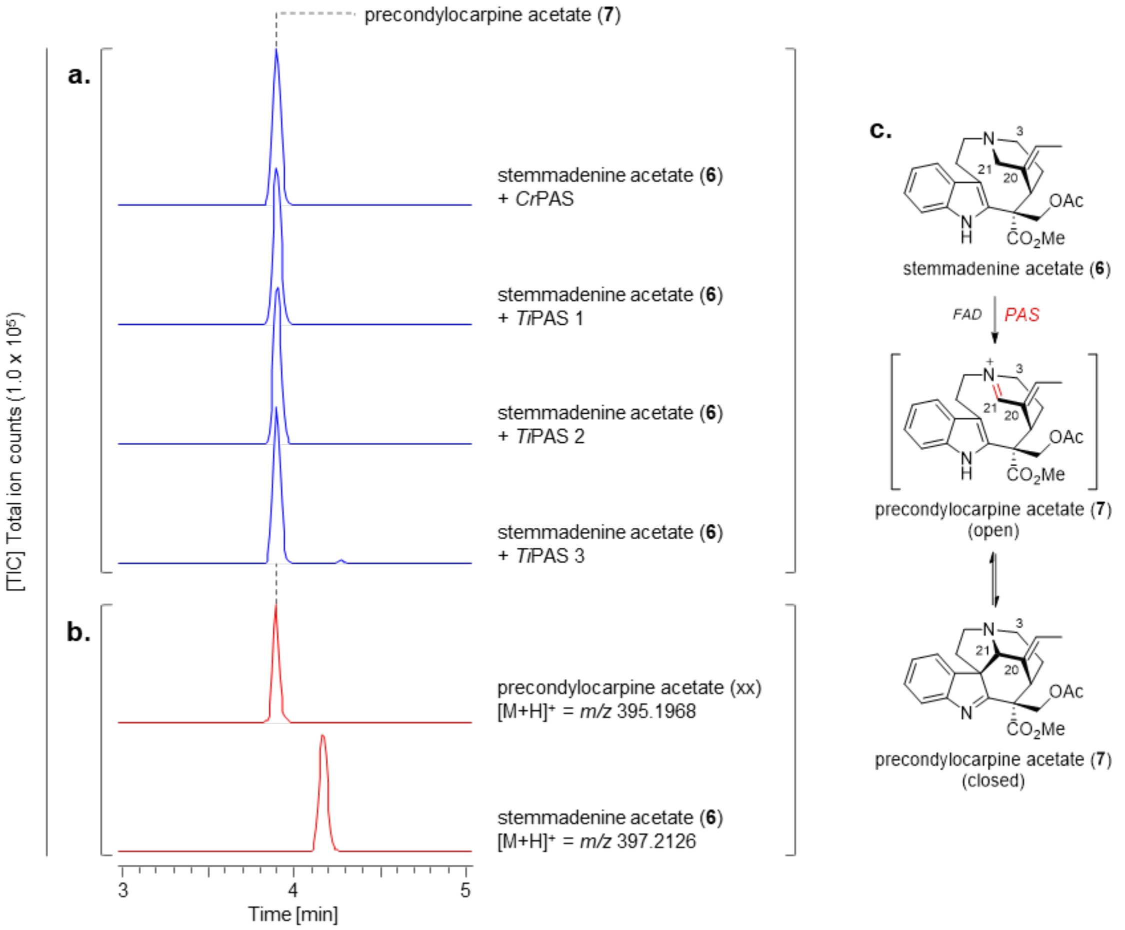


**Figure S18.** Production of precondylocarpine acetate (**7**) from stemmadenine acetate (**6**). **a)** TICs of *in vitro* enzyme assays showing the production of precondylocarpine acetate (**7**) from stemmadenine acetate (**6**) by PAS (oxidase) homologues described in this study. **b)** Authentic standards of product (precondylocarpine acetate (**7**)) and substrate stemmadenine acetate (**6**). **c)** Proposed reaction for the formation of precondylocarpine acetate (**7**) by PAS (oxidase).

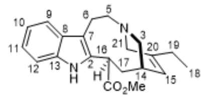

$C_{21}H_{26}N_2O_2$
$[M+H]^+$ = 339.2067

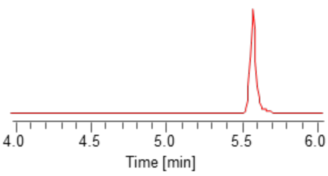


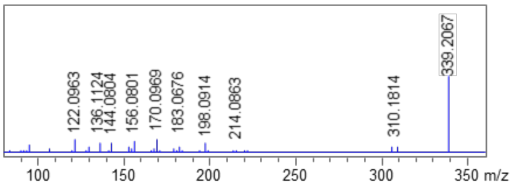


(–)-16(*R*)-carbomethoxycleavamine (**10**) or *ent*-**10a**

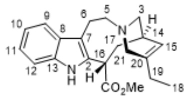

$C_{21}H_{26}N_2O_2$
$[M+H]^+$ = 339.2067

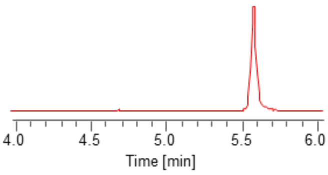


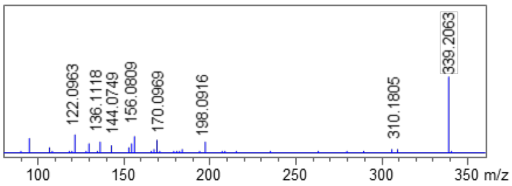


(+)-16(*S*)-carbomethoxycleavamine (**10a**)

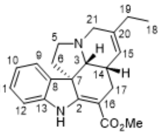

$C_{21}H_{24}N_2O_2$
$[M+H]^+$ = 337.1911

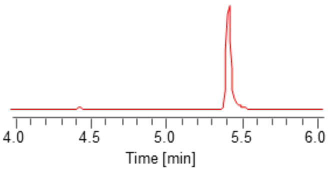


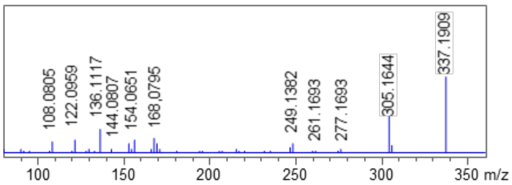


(+)-Ψ-tabersonine (**5**)

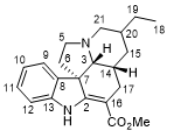

$C_{21}H_{26}N_2O_2$
$[M+H]^+$ = 339.2067

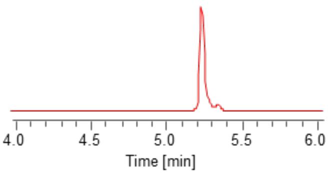


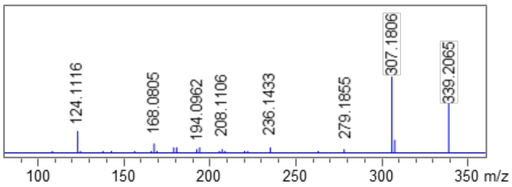


(+)-Ψ-vincadifformine (**9**)

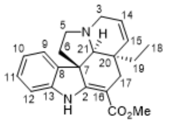

$C_{21}H_{24}N_2O_2$
$[M+H]^+$ = 337.1911

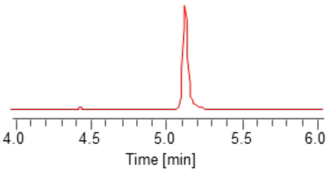


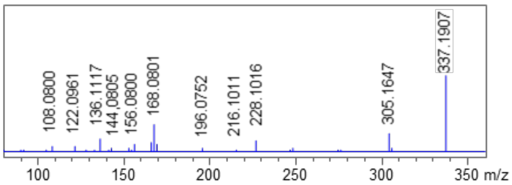


(–)-tabersonine (**2**)

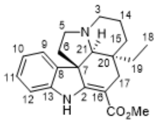

$C_{21}H_{26}N_2O_2$
$[M+H]^+$ = 339.2067

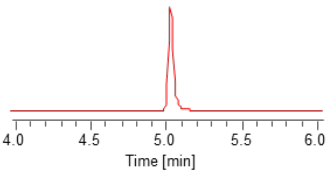


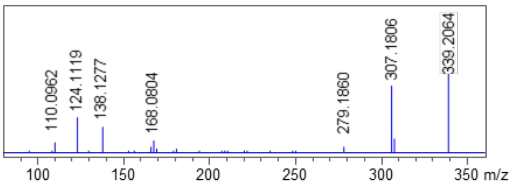


(–)-vincadifformine

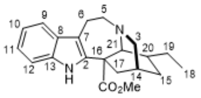

$C_{21}H_{26}N_2O_2$
$[M+H]^+$ = 339.2067

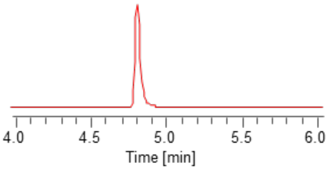


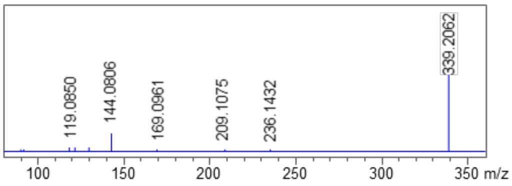


(–)-coronaridine (**4**)

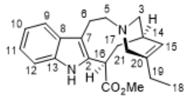

$C_{21}H_{26}N_2O_2$
$[M+H]^+$ = 339.2067

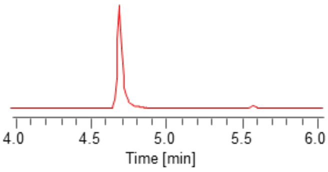


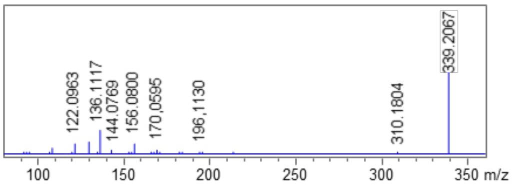


(+)-16(*R*)-carbomethoxycleavamine (**10a'**)

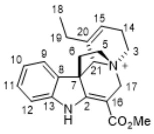

$C_{21}H_{25}N_2O_2^+$
$[M+H]^+$ = 337.1911

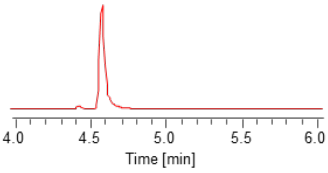


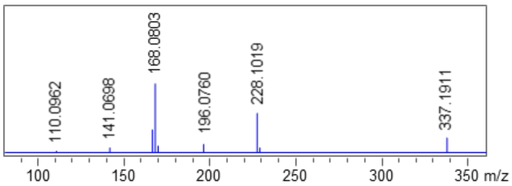


angryline (**1a**)

*Continued to next page.*

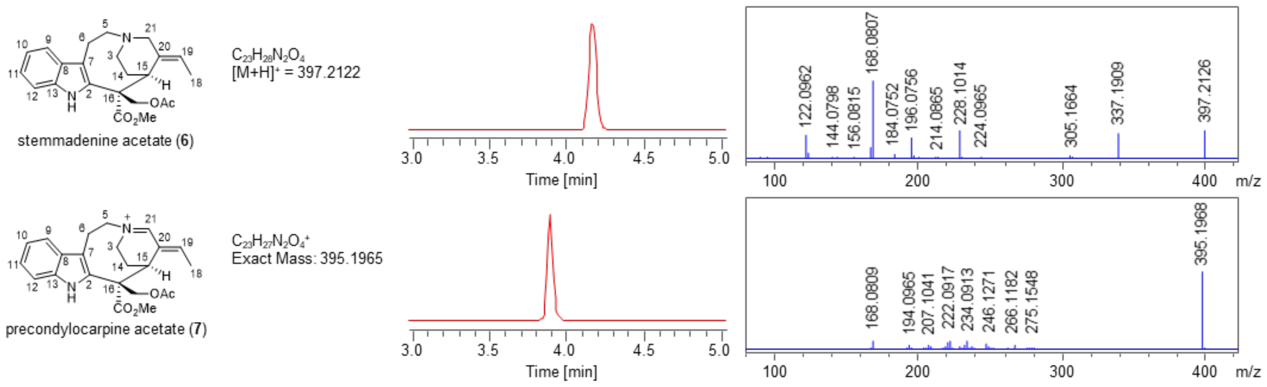


**Figure S19. Authentic standards of chemicals used in this study.** Atom numbering of molecules, total ion chromatograms (TICs) and $MS^2$ fragmentation spectra of authentic standards are shown.

### Synthesis of compounds

**Synthesis of racemic pseudo-tabersonine [Ψ-tabersonine] (5) and pseudo-vincadifformine [Ψ-vincadifformine] (9).**

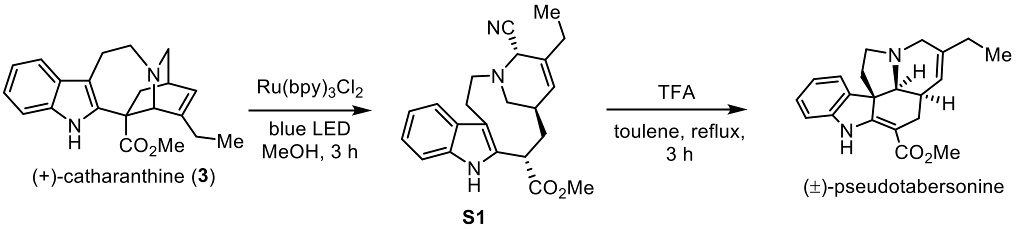


(±)-Ψ-tabersonine (**5a**) was prepared according to the reported method[15]. The reaction mixture containing (+)-catharanthine (85 mg, 0.253 mmol), $Ru(bpy)_3Cl_2$ (5.7 mg, 0.0076 mmol) and methanol (2.5 mL) was degassed *via* the freeze pump-thaw method (3 cycles) and the flask back-filled with argon. TMSCN (63 μL, 0.506 mmol) was added to the solution. The reaction mixture was irradiated with blue LED and stirred for 3 hours. The reaction was quenched with sat. $NaHCO_3$ (10 mL), extracted with EtOAc (5 mL x 3). The combined organic layers were washed with brine (5 mL) and dried with anhydrous $Na_2SO_4$. The solvents were evaporated in *vacuo* to afford product **S1** (66 mg, 72%).

The solution of **S1** (36 mg, 0.1 mmol) and toluene (2 mL) was degassed *via* the freeze pump-thaw method (3 cycles) and the flask back-filled with argon. Trifluoroacetic acid (7.4 μL, 0.1 mmol) was added to the solution. The reaction mixture was heated to 120 °C and stirred for 3 hours. The reaction was quenched with sat. $NaHCO_3$ (5 mL), extracted with EtOAC (5 mL x 3). The combined organic layers were washed with brine (5 mL) and dried with anhydrous $Na_2SO_4$, filtered and concentrated in *vacuo.* The residue was purified on silica gel chromatography (petroleum ether/EtOAc = 20/1) to afford (±)-Ψ-tabersonine (12 mg, 36%). The spectroscopic data of (±)-Ψ-tabersonine (**5**) are in accordance with the literature values reported[15]. Full NMR characterization is provided below.

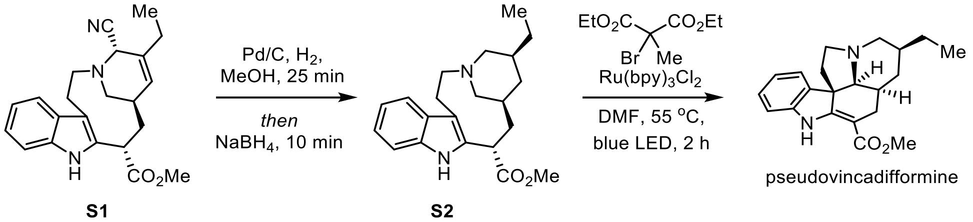


(±)-Ψ-vincadifformine (**9a**) was prepared according to the reported method[15]. The solution of **S1** (9 mg, 0.0248 mmol), 10% Pd/C (5.4 mg) and MeOH (1 mL) was degassed *via* the freeze pump-thaw method (3 cycles) and the flask back-filled with argon. Hydrogen was bubbled through the reaction from a balloon for 25 minutes. Then the reaction mixture was cooled to 0 °C and $NaBH_4$ (3.8 mg, 0.0992 mmol) was added and stirred at the same temperature for 10 minutes. The reaction was filtered through a column of Celite, washed through with ethyl acetate, and concentrated in *vacuo.* The resulting residue **S2** was used directly in the next step. The above residue was dissolved in anhydrous DMF (1 mL) and $Ru(bpy)_3Cl_2$ (0.2 mg, 0.000267 mmol) was added with stirring. The solution was degassed *via* the freeze pump-thaw method (3 cycles) and the flask back-filled with argon. Diethyl 2-bromo-2-methylmalonate (14 μL, 0.0744 mmol) was added to the solution. The reaction mixture was heated to 55 °C and irradiated with blue LED. After 2 hours, the reaction was diluted with EtOAc (2 mL) and partitioned with water (2 mL). Triethylamine (0.5 mL) was added to the separatory funnel, and the mixture was extracted with ethyl acetate (2 mL, x3). The combined organic layers were washed with brine (5 mL) dried with anhydrous $Na_2SO_4$, filtered and concentrated in *vacuo.* The residue was purified by silica gel chromatography (petroleum ether/EtOAc = 20/1, argon-sparged) under argon to afford (±)-Ψ-vincadifformine (**9a**) (2 mg, 24%). The spectroscopic data of Ψ-vincadifformine are in accordance with the literature values reported[15]. NMR spectra are shown below.

**Supplementary NMR spectra**

**NMR data for synthesized (–)-Ψ-tabersonine (5a) (pseudo-tabersonine)**

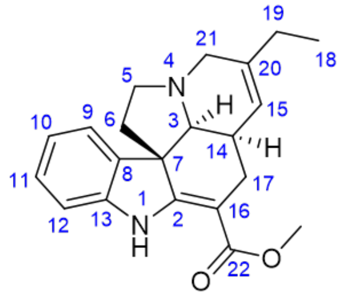


(–)-Ψ-tabersonine (**5a**)
$^{1}$H-NMR, MeOH-$d_3$
500 MHz, 25 °C

| pos. | $\delta_H$ | mult. | $J_{HH}$ | $\delta_C$ |
|---|---|---|---|---|
| 1 | 9.29 | *bs* | - | - |
| 2 | - | - | - | 166.2 |
| 3α | 3.16 | *m* | - | 66.4 |
| 4 | - | - | - | - |
| 5α | 2.92 | *m* | - | 51.6 |
| 5β | 3.05 | *m* | - | 51.6 |
| 6α | 1.88 | *ddd* | 11.9/5.2/2.0 | 45.2 |
| 6β | 2.03 | *m* | - | 45.2 |
| 7 | - | - | - | 56.8 |
| 8 | - | - | - | 138.1 |
| 9 | 7.34 | *bd* | 7.4 | 122.3 |
| 10 | 6.87 | *ddd* | 7.4/7.3/1.0 | 121.3 |
| 11 | 7.13 | *ddd* | 7.7/7.3/1.0 | 128.5 |
| 12 | 6.92 | *bd* | 7.7 | 110.3 |
| 13 | - | - | - | 145.0 |
| 14α | 1.79 | *m* | - | 37.3 |
| 15 | 5.57 | *m* | - | 122.3 |
| 16 | - | - | - | 95.3 |
| 17α | 2.72 | *dd* | 15.1/3.3 | 27.1 |
| 17β | 2.13 | *dd* | 15.1/11.6 | 27.1 |
| 18 | 1.07 | *t* | 7.5 | 12.4 |
| 19a | 2.07 | *m* | - | 28.4 |
| 19b | 2.07 | *m* | - | 28.4 |
| 20 | - | - | - | 139.6 |
| 21a | 3.40 | *m* | - | 53.5 |
| 21b | 3.40 | *m* | - | 53.5 |
| 22 | - | - | - | 169.3 |
| OMe | 3.78 | *s* | - | 51.0 |

**(–)-Ψ-tabersonine (5a): Phase sensitive HSQC**, full range in MeOH-$d_3$

Red: $CH_2$, black: CH, $CH_3$

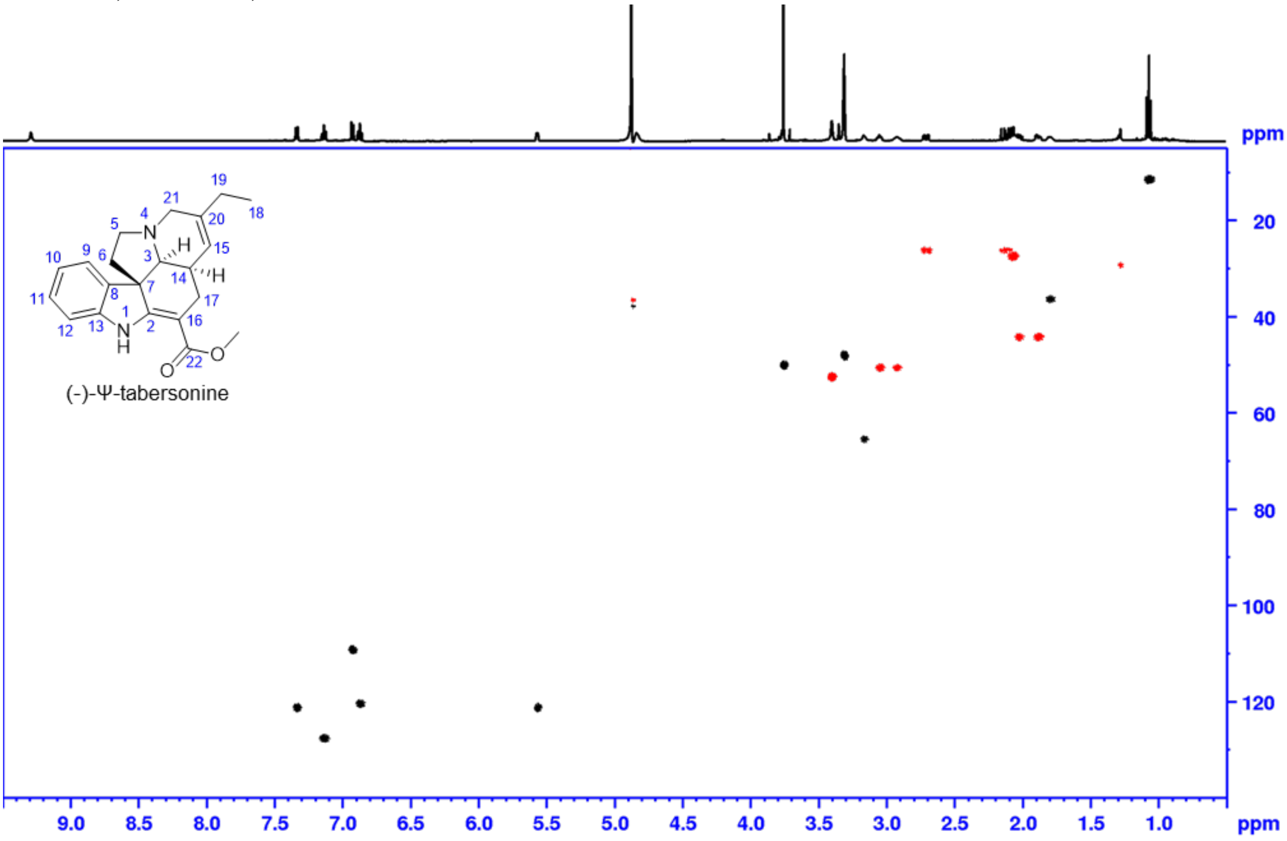


**(–)-Ψ-tabersonine (5a): Phase sensitive HSQC**, aliphatic range in MeOH-$d_3$

Shaded areas mark impurities and solvent, red: $CH_2$, black: CH, $CH_3$

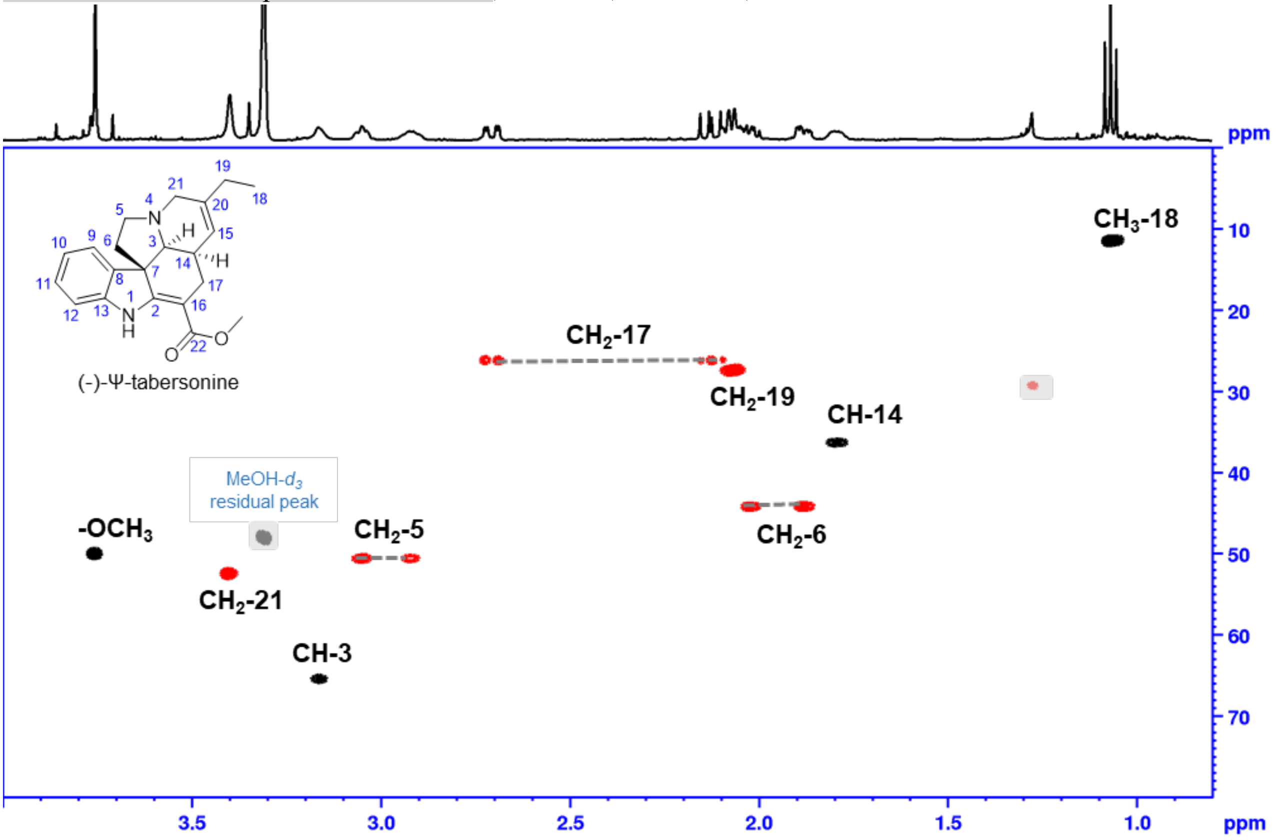

**Structure of (–)-Ψ-tabersonine (5a).** Structure optimized using Gaussian 16W (PM6, solvent MeOH). Important ROESY correlations in green.

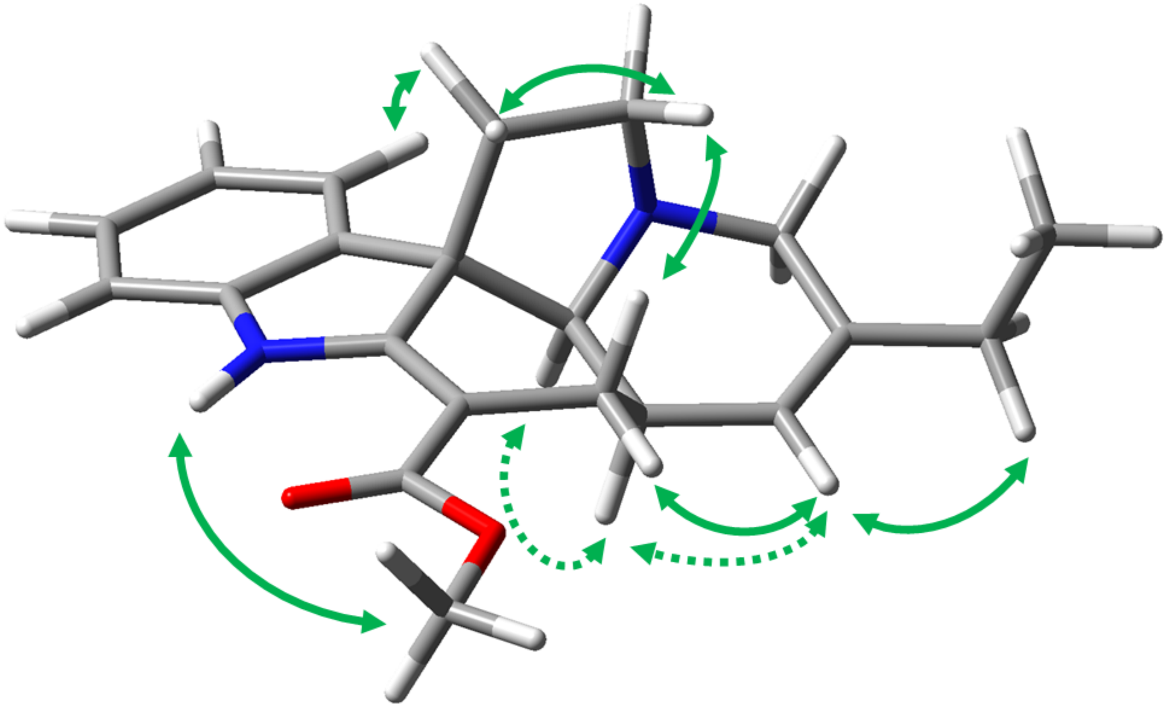

**NMR data for (+)-Ψ-tabersonine-$d_2$ (5-d2) (Enzymatic product, formic acid salt).**

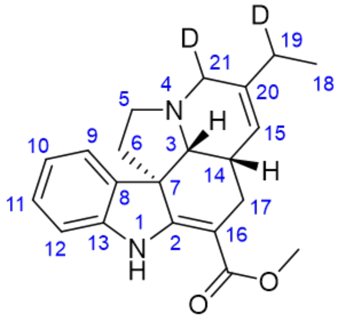


(+)-Ψ-tabersonine-$d_2$ (**5-d2**)
$^1$H-NMR, MeOH-$d_3$
700 MHz, 25 °C

| pos. | δ$_H$ | mult. | J$_{HH}$ | δ$_C$ |
|---|---|---|---|---|
| 1 | 9.27 | *bs* | - | - |
| 2 | - | - | - | 166.8 |
| 3β | 3.05 | *m* | - | 66.4 |
| 4 | - | - | - | - |
| 5α | 2.99 | *ddd* | 8.5/6.8/2.0 | 51.5 |
| 5β | 2.84 | *ddd* | 9.8/8.5/5.3 | 51.5 |
| 6α | 2.00 | *ddd* | 11.9/9.8/6.8 | 45.4 |
| 6β | 1.85 | *ddd* | 11.9/5.3/2.0 | 45.4 |
| 7 | - | - | - | 56.5 |
| 8 | - | - | - | 138.6 |
| 9 | 7.32 | *bd* | 7.6 | 122.2 |
| 10 | 6.86 | *bdd* | 7.6/7.6 | 121.2 |
| 11 | 7.12 | *bdd* | 7.8/7.6 | 128.4 |
| 12 | 6.92 | *bd* | 7.8 | 110.1 |
| 13 | - | - | - | 144.8 |
| 14α | 1.73 | *m* | - | 37.5 |
| 15 | 5.54 | *bd* | 4.9 | 122.2 |
| 16 | - | - | - | 95.3 |
| 17α | 2.13 | *dd* | 15.0/11.6 | 27.3 |
| 17β | 2.68 | *ddd* | 15.0/3.6/0.9 | 27.3 |
| 18 | 1.05 | *d* | 7.5 | 12.4 |
| 19a* | 2.05 | *m* | - | 28.2 |
| 19b | - | - | - | 28.2 |
| 20 | - | - | - | 140.0 |
| 21a* | 3.32 | *m*** | - | 53.1 |
| 21b | - | - | - | 53.1 |
| 22 | - | - | - | 169.5 |
| OMe | 3.75 | *s* | - | 51.0 |
| *as CH signal | | | | |
| ** overlapped signals J unresolved | | | | |

**(+)-Ψ-tabersonine-*d*₂ (5-d2): Phase sensitive HSQC**, full range in MeOH-*d*₃

red: $CH_2$, black: CH, $CH_3$

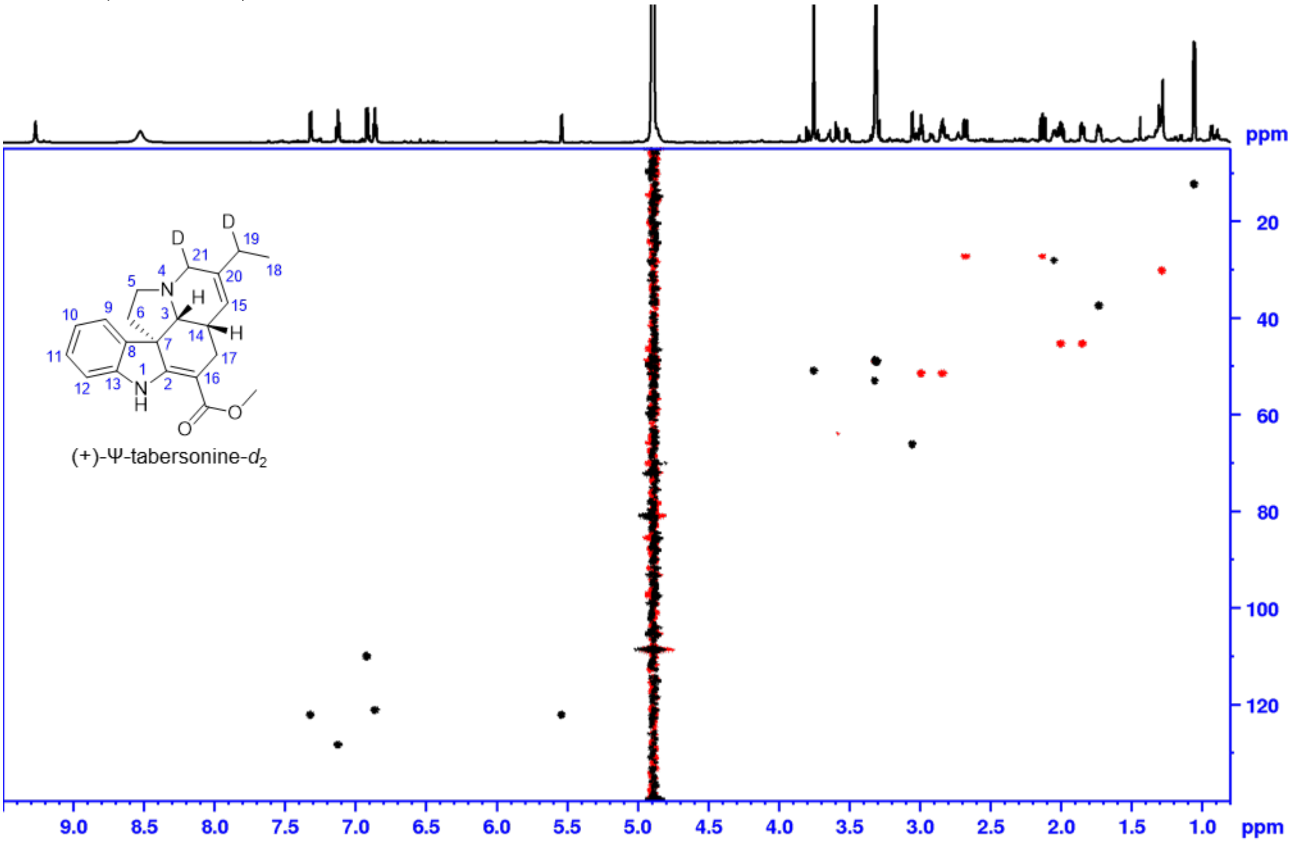


**(+)-Ψ-tabersonine-*d*₂ (5-d2): Phase sensitive HSQC**, aliphatic range in MeOH-*d*₃

Shaded areas mark impurities and solvent, red: $CH_2$, black: CH, $CH_3$

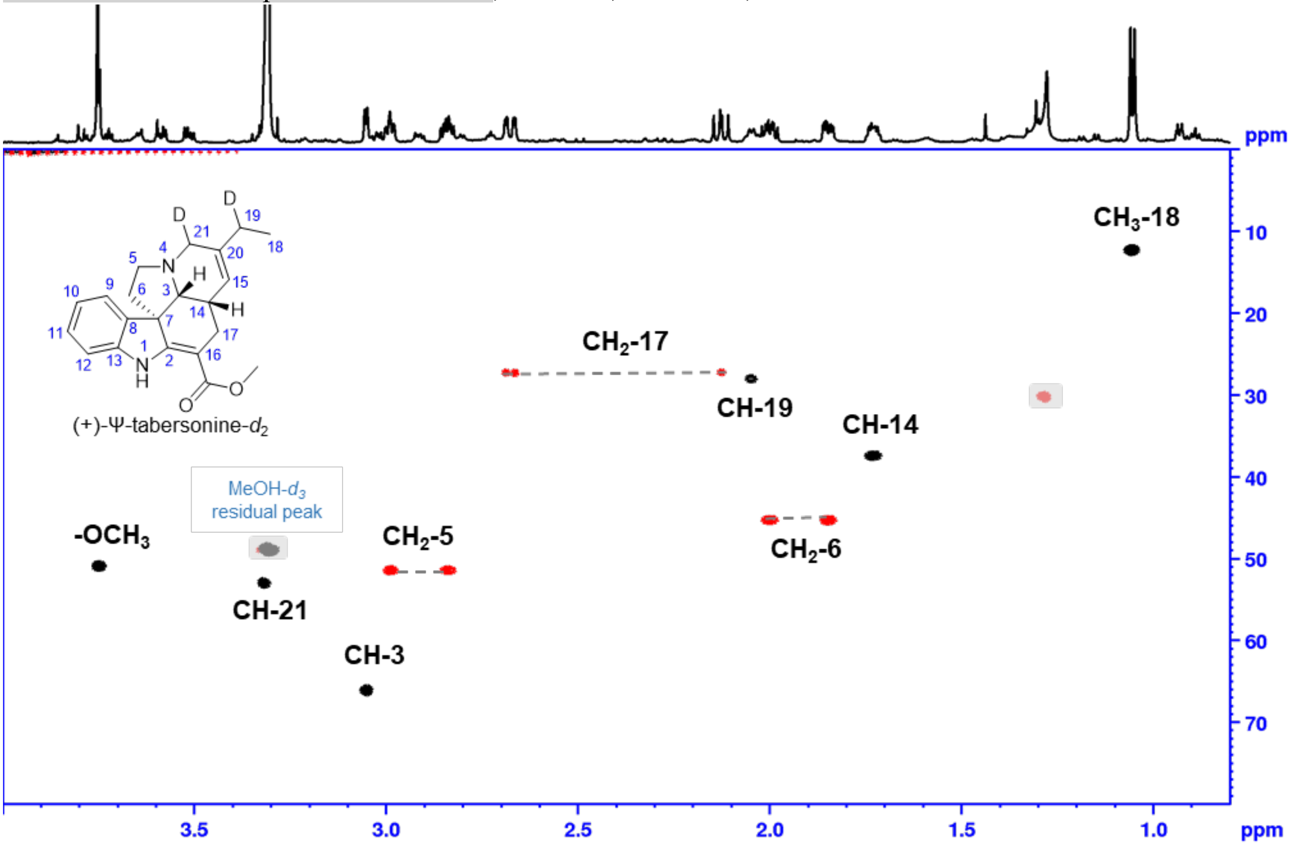

**NMR data for synthesized (–)-Ψ-vincadifformine (9a) (pseudo-vincadifformine)**

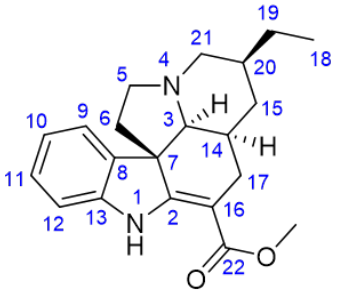


(–)-Ψ-vincadifformine (**9a**)
$^{1}$H-NMR, MeOH-$d_3$
700 MHz, 25 °C

| pos. | $\delta_H$ | mult. | $J_{HH}$ | $\delta_C$ |
|---|---|---|---|---|
| 1 | 9.32 | *bs* | - | - |
| 2 | - | - | - | 164.6 |
| 3α | 3.55 | *m* | - | 66.4 |
| 4 | - | - | - | - |
| 5a | 3.13 | *m* | - | 53.0 |
| 5b | 3.13 | *m* | - | 53.0 |
| 6α | 1.86 | *m* | - | 44.0 |
| 6β | 2.16 | *m* | - | 44.0 |
| 7 | - | - | - | 56.5 |
| 8 | - | - | - | 136.7 |
| 9 | 7.32 | *bd* | 7.3 | 122.1 |
| 10 | 6.89 | *bdd* | 7.7/7.3 | 121.4 |
| 11 | 7.16 | *bdd* | 7.8/7.7 | 129.0 |
| 12 | 6.94 | *bd* | 7.8 | 110.4 |
| 13 | - | - | - | 144.7 |
| 14α | 1.68 | *m* | - | 35.6 |
| 15α | 2.02 | *m* | - | 31.8 |
| 15β | 1.37 | *ddd* | 13.8/8.9/4.6 | 31.8 |
| 16 | - | - | - | 96.2 |
| 17α | 2.65 | *dd* | 15.4/3.4 | 26.9 |
| 17β | 2.27 | *dd* | 15.4/12.2 | 26.9 |
| 18 | 0.96 | *t* | 7.4 | 11.8 |
| 19a | 1.48 | *m* | - | 29.0 |
| 19b | 1.42 | *m* | - | 29.0 |
| 20 | 1.83 | *m* | - | 35.0 |
| 21α | 3.20 | *m* | - | 55.1 |
| 21β | 2.93 | *dd* | 9.9/9.4 | 55.1 |
| 22 | - | - | - | 169.9 |
| OMe | 3.78 | *s* | - | 51.1 |

**(–)-Ψ-vincadifformine (9a): Phase sensitive HSQC**, full range in MeOH-$d_3$

Shaded areas mark impurities and solvent, red: $CH_2$, black: CH, $CH_3$

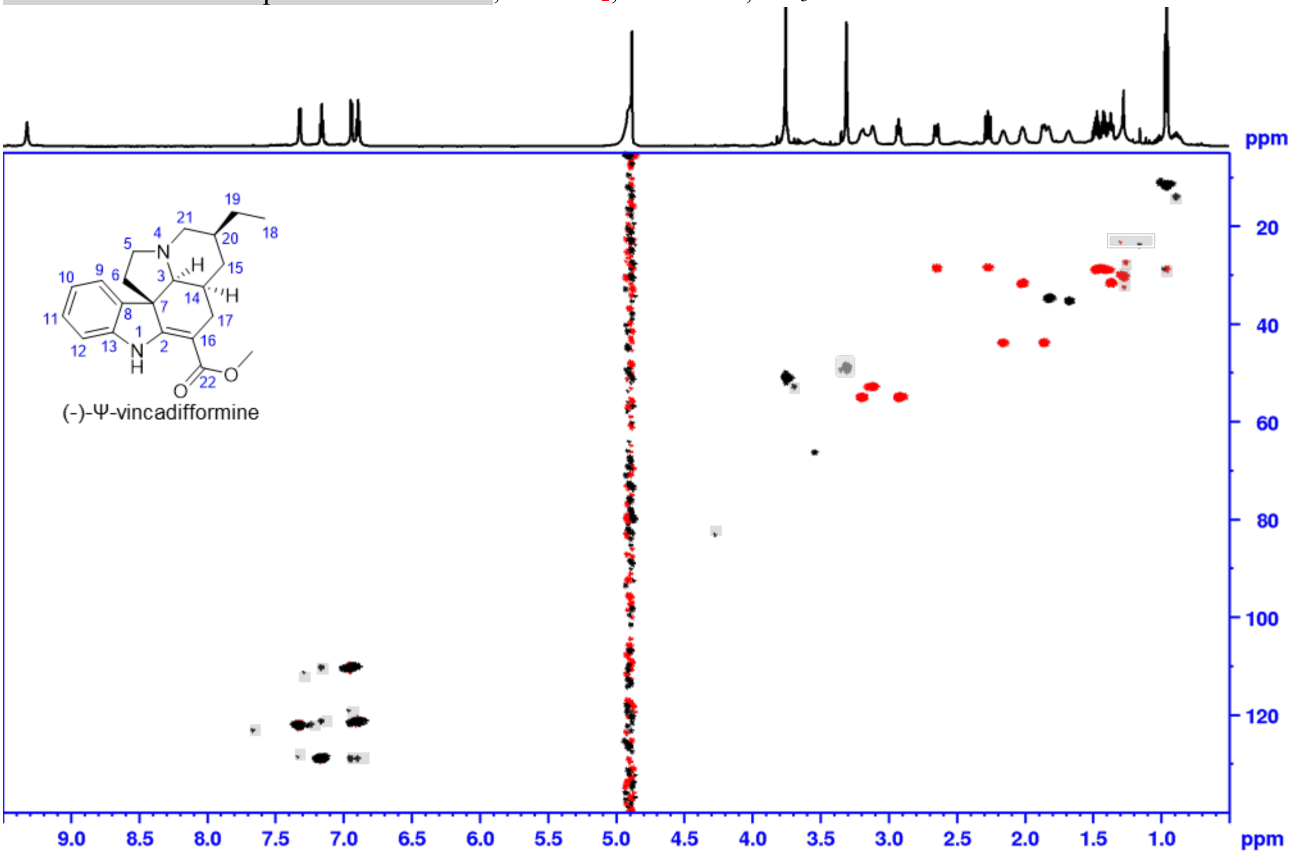


**(–)-Ψ-vincadifformine (9a): Phase sensitive HSQC**, aliphatic range in MeOH-$d_3$

Shaded areas mark impurities and solvent, red: $CH_2$, black: CH, $CH_3$

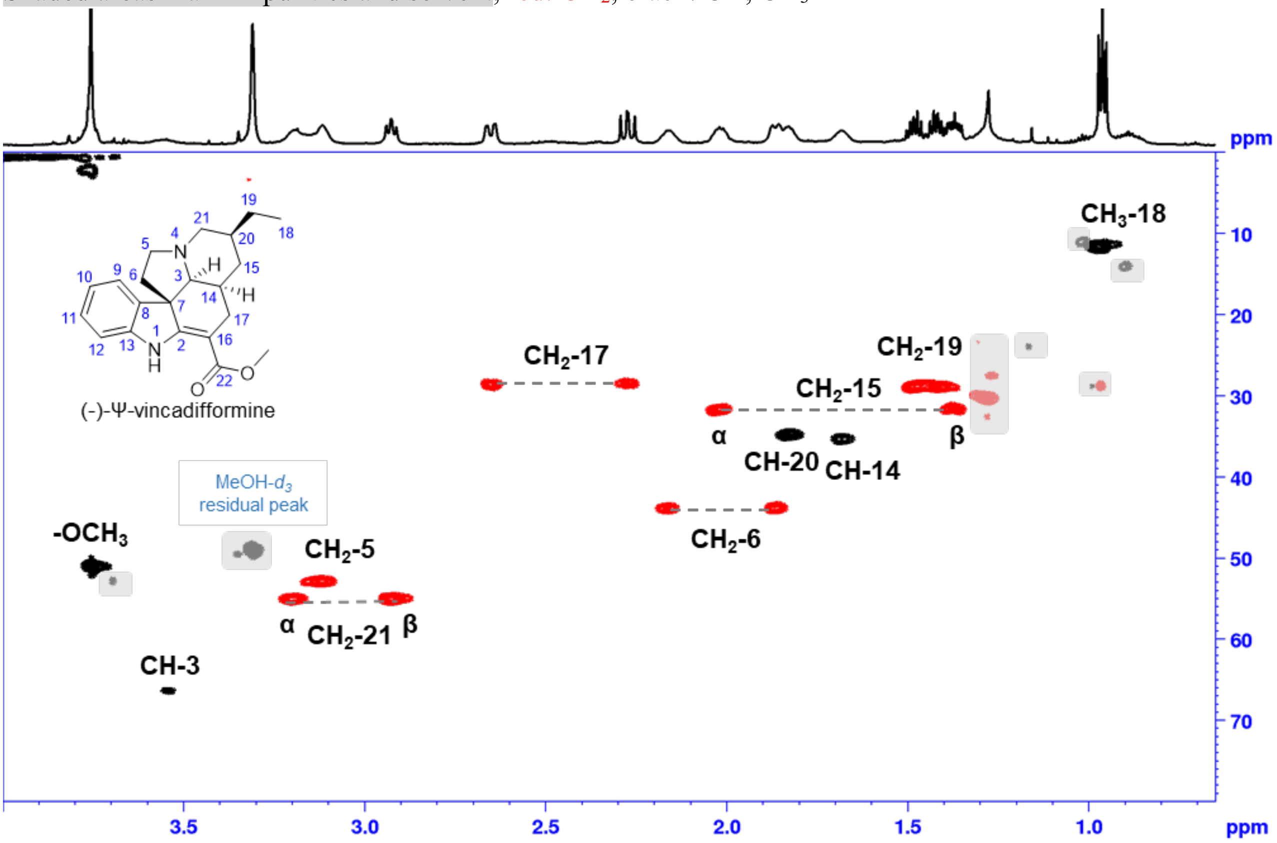

**Structure of (–)-Ψ-vincadifformine (9a).** Structure optimized using Gaussian 16W (PM6, solvent MeOH). Important ROESY correlations in green.

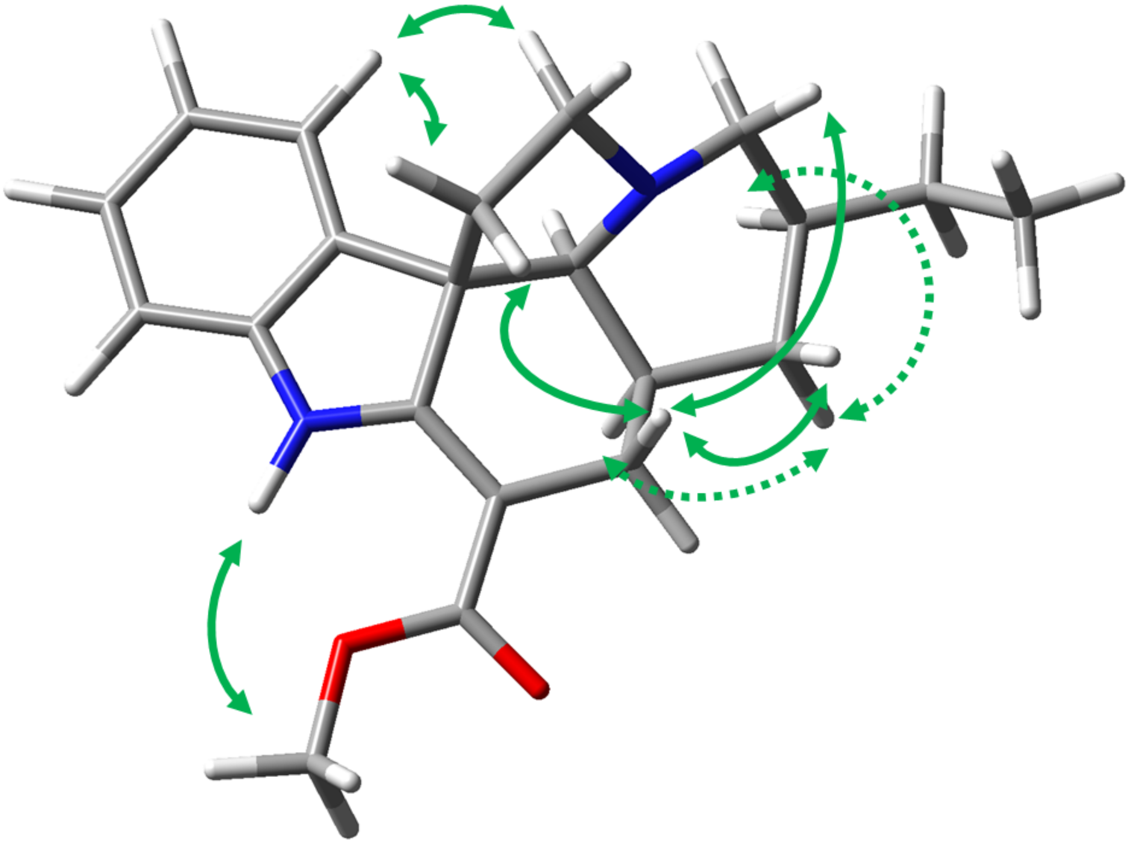

**NMR data for (+)-Ψ-vincadifformine-$d_3$ (9-d3) (pseudo-vincadifformine-$d_3$) (Enzymatic product, formic acid salt).**

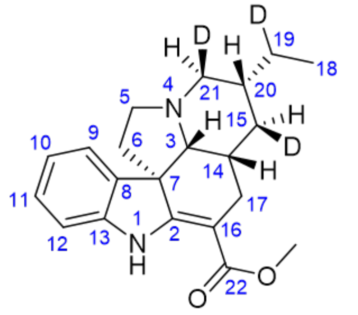


(+)-Ψ-vincadifformine-$d_3$ (**9-d3**)
$^1$H-NMR, MeOH-$d_3$
700 MHz, 25 °C

| pos. | δ H | mult. | J HH | δ C |
|---|---|---|---|---|
| 1 | 9.20 | *bs* | - | - |
| 2 | - | - | - | 166.7 |
| 3β | 3.12 | *bd* | 3.8 | 66.9 |
| 4 | - | - | - | - |
| 5a | 2.92 | *m* | - | 52.6 |
| 5b | 2.81 | *m*** | - | 52.6 |
| 6α | 2.00 | *ddd* | 11.2/11.2/6.3 | 45.4 |
| 6β | 1.72 | *bdd* | 11.2/4.6 | 45.4 |
| 7 | - | - | - | 56.9 |
| 8 | - | - | - | 138.6 |
| 9 | 7.25 | *bd* | 7.4 | 122.5 |
| 10 | 6.86 | *bdd* | 7.5/7.4 | 121.4 |
| 11 | 7.12 | *bdd* | 7.8/7.5 | 128.8 |
| 12 | 6.91 | *bd* | 7.8 | 110.4 |
| 13 | - | - | - | 145.1 |
| 14β | 1.46 | *m* | - | 37.1 |
| 15α* | 1.35 | *m* | - | 32.8 |
| 15β | - | - | - | 32.8 |
| 16 | - | - | - | 96.3 |
| 17α | 2.28 | *dd* | 14.7/11.8 | 28.2 |
| 17β | 2.55 | *ddd* | 14.7/3.5/0.8 | 28.2 |
| 18 | 0.93 | *d* | 7.4 | 12.3 |
| 19a* | 1.47 | *m* | - | 29.2 |
| 19b | - | - | - | 29.2 |
| 20 | 1.67 | *ddd* | 7.3/7.3/7.3 | 36.2 |
| 21α* | 2.80 | *m*** | - | 55.3 |
| 21β | - | - | - | 55.3 |
| 22 | - | - | - | 169.8 |
| OMe | 3.75 | *s* | - | 51.3 |
| *as CH signal | | | | |
| ** overlapped signals J unresolved | | | | |

**(+)-Ψ-vincadifformine-$d_3$ (9-d3): Phase sensitive HSQC**, full range in MeOH-$d_3$

Shaded areas mark impurities and solvent, red: $CH_2$, black: CH, $CH_3$

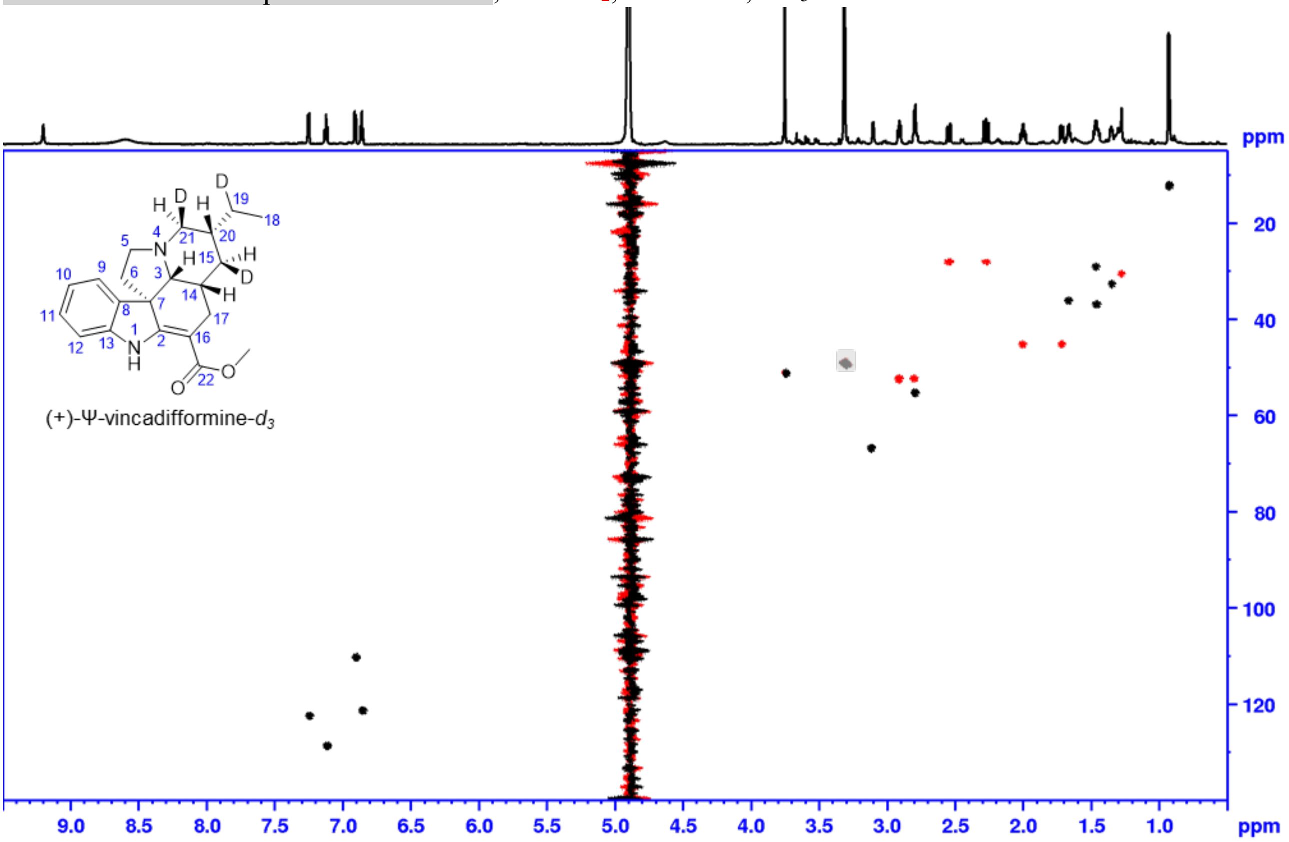


**(+)-Ψ-vincadifformine-$d_3$ (9-d3): Phase sensitive HSQC**, aliphatic range in MeOH-$d_3$

Shaded areas mark impurities and solvent, red: $CH_2$, black: CH, $CH_3$

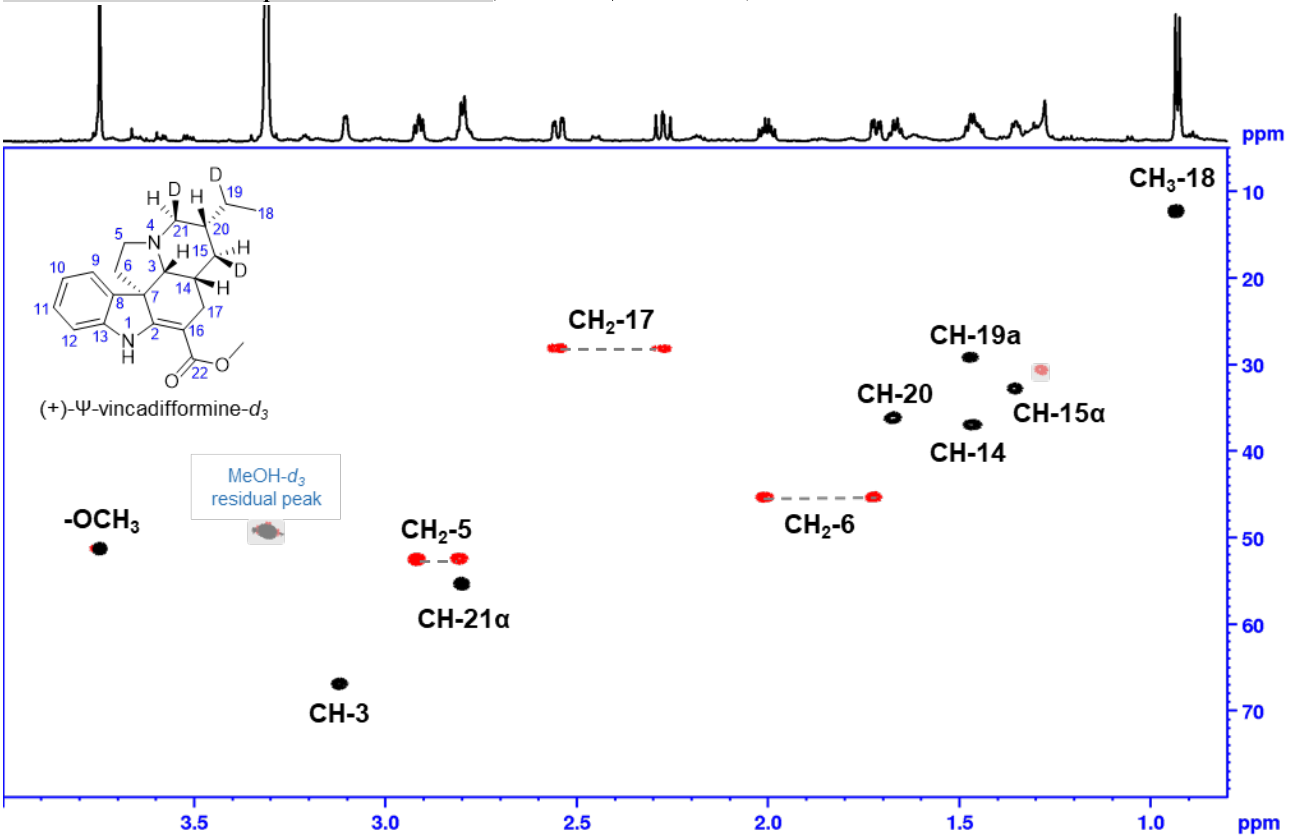

**NMR data for (–)-tabersonine-*d* (2-d)**
**(Enzymatic product, formic acid salt).**

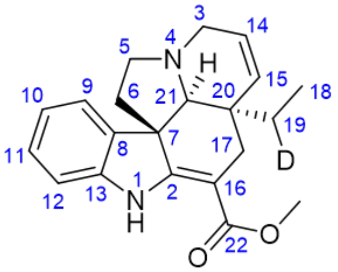


(–)-tabersonine-*d* (**2-d**)
$^1$H-NMR, MeOH-$d_3$
700 MHz, 25 °C

| pos. | $\delta_H$ | mult. | $J_{HH}$ | $\delta_C$ |
|---|---|---|---|---|
| 1 | 9.33 | *bs* | - | - |
| 2 | - | - | - | 167.1 |
| 3α | 3.39 | *bd* | 15.6 | 51.2 |
| 3β | 3.55 | *bdd* | 15.6/3.8 | 51.2 |
| 4 | - | - | - | - |
| 5α | 2.90 | *m* | - | 51.7 |
| 5β | 3.14 | *m* | - | 51.7 |
| 6α | 1.81 | *dd* | 11.6/4.5 | 45.5 |
| 6β | 2.05 | *ddd* | 11.6/11.6/6.3 | 45.5 |
| 7 | - | - | - | 56.2 |
| 8 | - | - | - | 138.2 |
| 9 | 7.31 | *bd* | 7.4 | 122.1 |
| 10 | 6.87 | *bdd* | 7.6/7.4 | 121.7 |
| 11 | 7.15 | *bdd* | 7.7/7.6 | 129.1 |
| 12 | 6.95 | *bd* | 7.7 | 110.8 |
| 13 | - | - | - | 144.8 |
| 14 | 5.83 | *bdd* | 9.8/3.8 | 125.1 |
| 15 | 5.76 | *bd* | 9.8 | 134.1 |
| 16 | - | - | - | 92.5 |
| 17α | 2.61 | *bd* | 15.4 | 29.6 |
| 17β | 2.43 | *dd* | 15.4 | 29.6 |
| 18 | 0.63 | *d* | 7.4 | 7.6 |
| 19a* | 0.90 | *q* | 7.4 | 27.5 |
| 19b | - | - | - | 27.5 |
| 20 | - | - | - | 42.2 |
| 21α | 2.96 | *bs* | - | 71.3 |
| 22 | - | - | - | 170.0 |
| OMe | 3.76 | *s* | - | 51.4 |
| *as CH signal | | | | |

**(–)-tabersonine-*d* (2-d): Phase sensitive HSQC**, full range in MeOH-$d_3$

Shaded areas mark impurities and solvent, red: $CH_2$, black: CH, $CH_3$

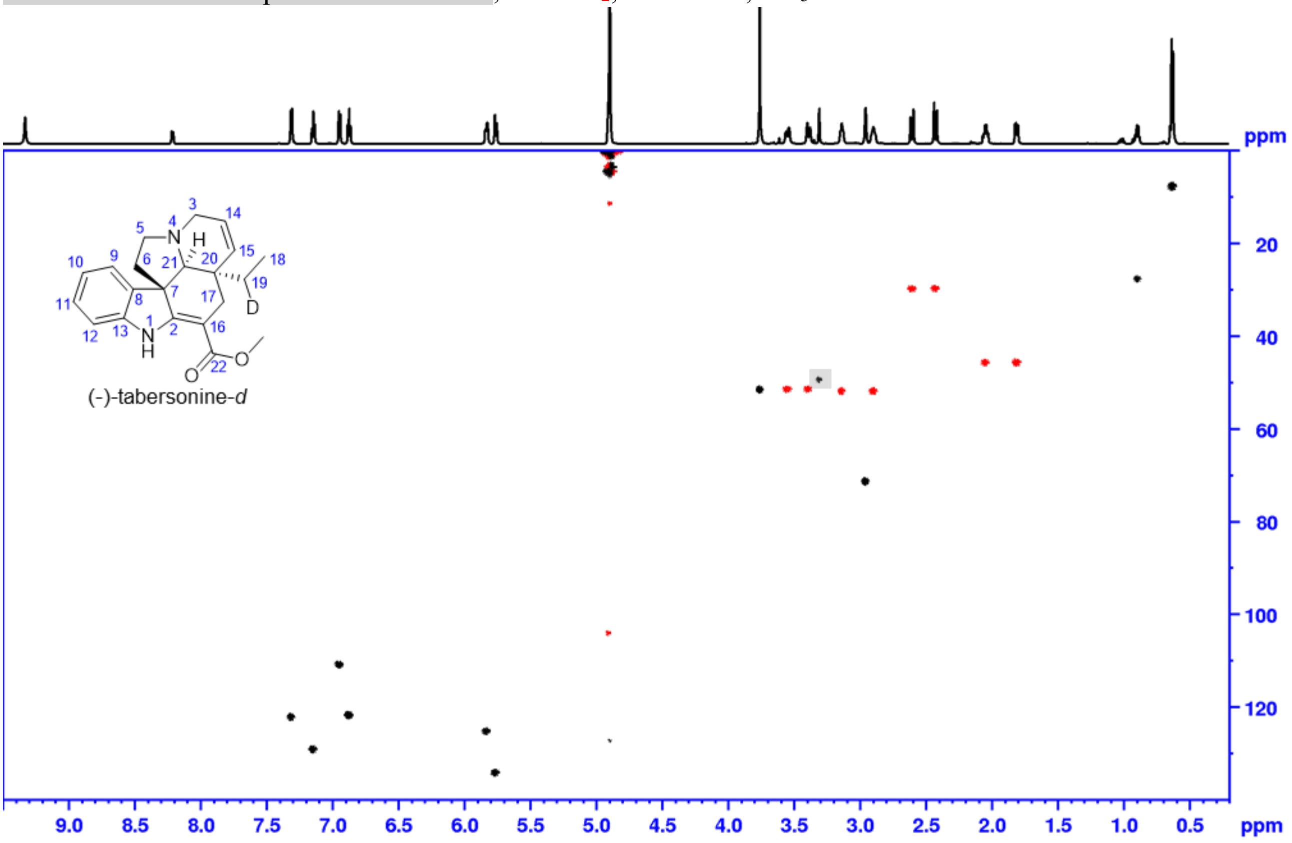


**(–)-tabersonine-*d* (2-d): Phase sensitive HSQC**, aliphatic range in MeOH-$d_3$

Shaded areas mark impurities and solvent, red: $CH_2$, black: CH, $CH_3$

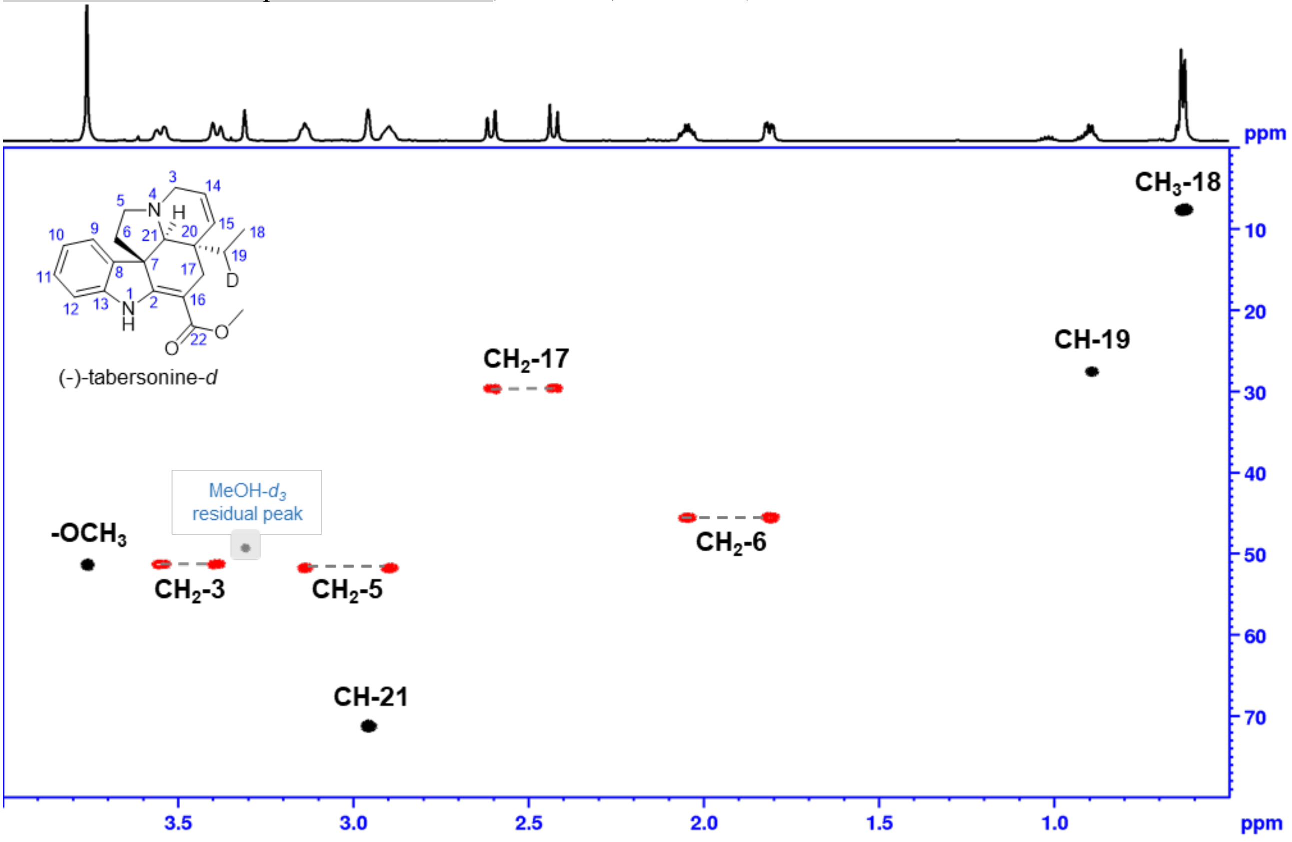

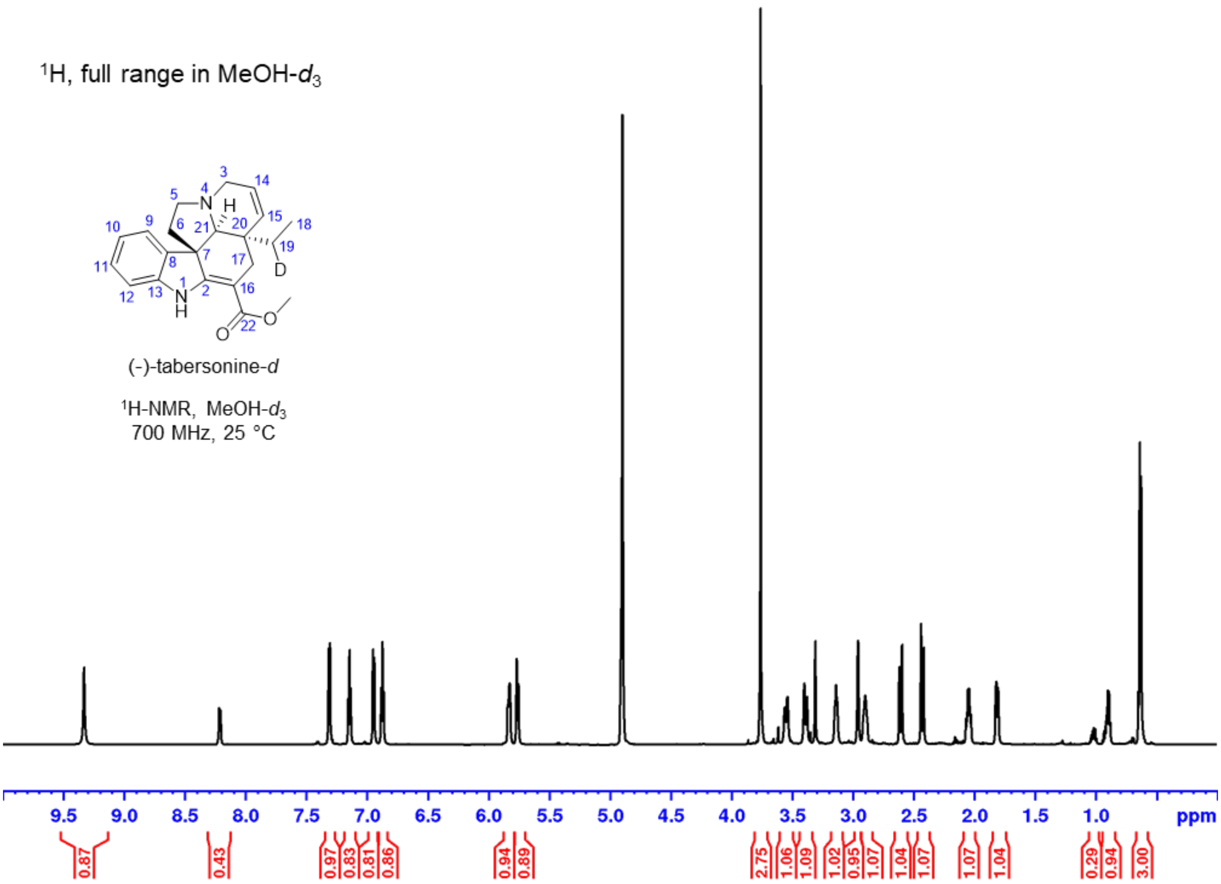

$^1$H, full range in MeOH-$d_3$
(-)-tabersonine-*d*
$^1$H-NMR, MeOH-$d_3$
700 MHz, 25 °C
ppm


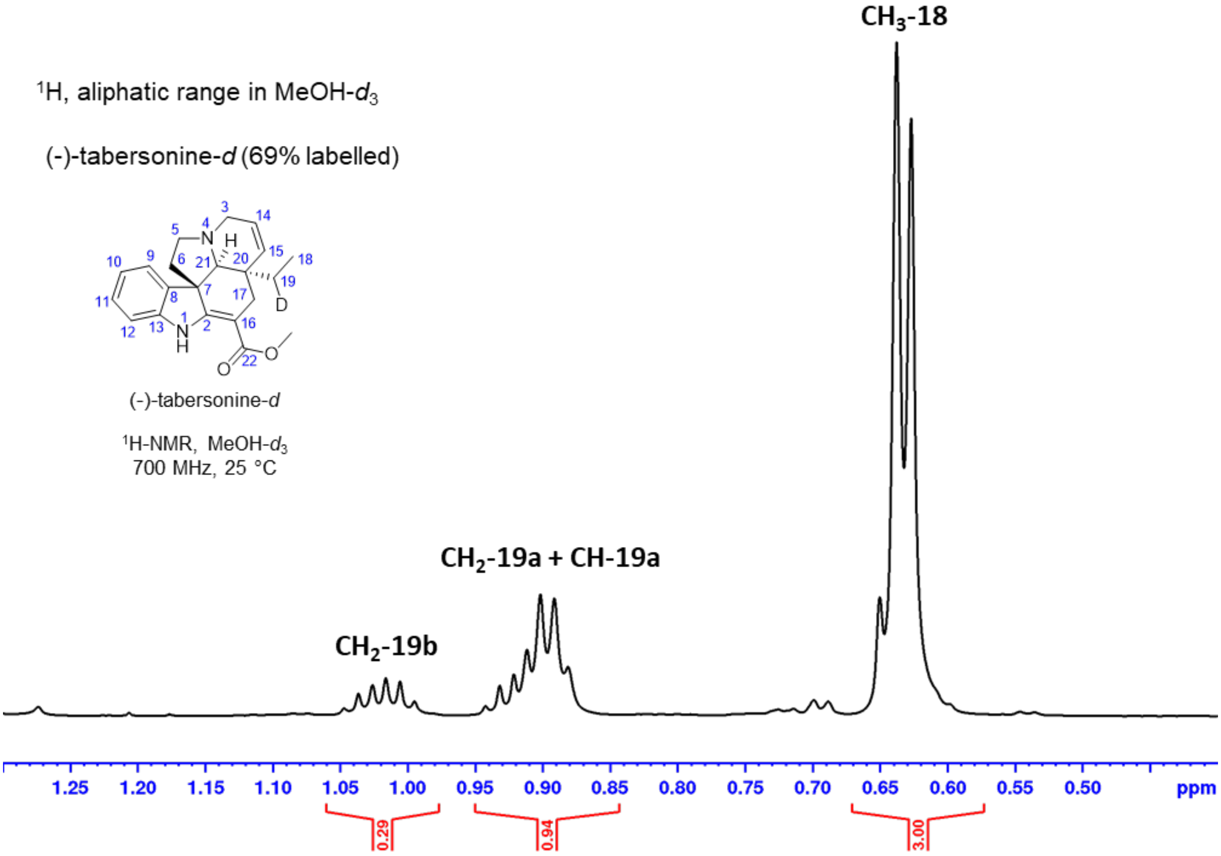

$^1$H, aliphatic range in MeOH-$d_3$
(-)-tabersonine-*d* (69% labelled)
(-)-tabersonine-*d*
$^1$H-NMR, MeOH-$d_3$
700 MHz, 25 °C
$CH_3$-18
$CH_2$-19a + CH-19a
$CH_2$-19b
ppm

**NMR data for (–)-16(*R*)-carbomethoxycleavamine (10).**

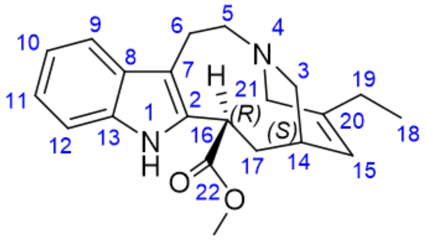


(–)-16(*R*)-carbomethoxycleavamine (**10**)
$^{1}$H-NMR, MeOH-$d_3$
700 MHz, 25 °C

| pos. | $\delta_H$ | mult. | $J_{HH}$ | $\delta_C$ |
|---|---|---|---|---|
| 1 | 10.1 | *bs* | - | - |
| 2 | - | - | - | 135.1 |
| 3a | 2.33 | *m*** | - | 53.4 |
| 3b | 2.22 | *m*** | - | 53.4 |
| 4 | - | - | - | - |
| 5a | 2.72 | *ddd* | 13.3/2.9/2.9 | 53.1 |
| 5b | 2.30 | *m*** | - | 53.1 |
| 6a | 2.85 | *m*** | - | 26.0 |
| 6b | 2.82 | *m*** | - | 26.0 |
| 7 | - | - | - | 110.5 |
| 8 | - | - | - | 127.6 |
| 9 | 7.4 | *bd* | 7.8 | 117.3 |
| 10 | 6.95 | *bdd* | 7.8/7.6 | 118.0 |
| 11 | 7.02 | *bdd* | 7.8/7.6 | 120.6 |
| 12 | 7.29 | *bd* | 7.8 | 110.3 |
| 13 | - | - | - | 136.4 |
| 14 | 2.14 | *bs* | - | 34.6 |
| 15 | 5.31 | *d* | 4.1 | 121.9 |
| 16 | 5.27 | *d* | 10.1 | 38.5 |
| 17a | 2.30 | *m*** | - | 36.5 |
| 17b | 2.23 | *m*** | - | 36.5 |
| 18 | 1.06 | *t* | 7.4 | 11.7 |
| 19a | 2.05 | *q* | 7.4 | 27.3 |
| 19b | 2.05 | *q* | 7.4 | 27.3 |
| 20 | - | - | - | 141.3 |
| 21a | 3.16 | *d* | 15.4 | 54.8 |
| 21b | 3.03 | *d* | 15.4 | 54.8 |
| 22 | - | - | - | 175.1 |
| OMe | 3.61 | *s* | - | 50.8 |
| ** overlapped signals J unresolved | | | | |

**(–)-16(*R*)-carbomethoxycleavamine (10):**
**$^{1}$H NMR** with water suppression (zgpr) full range in MeOH-*d*$_3$

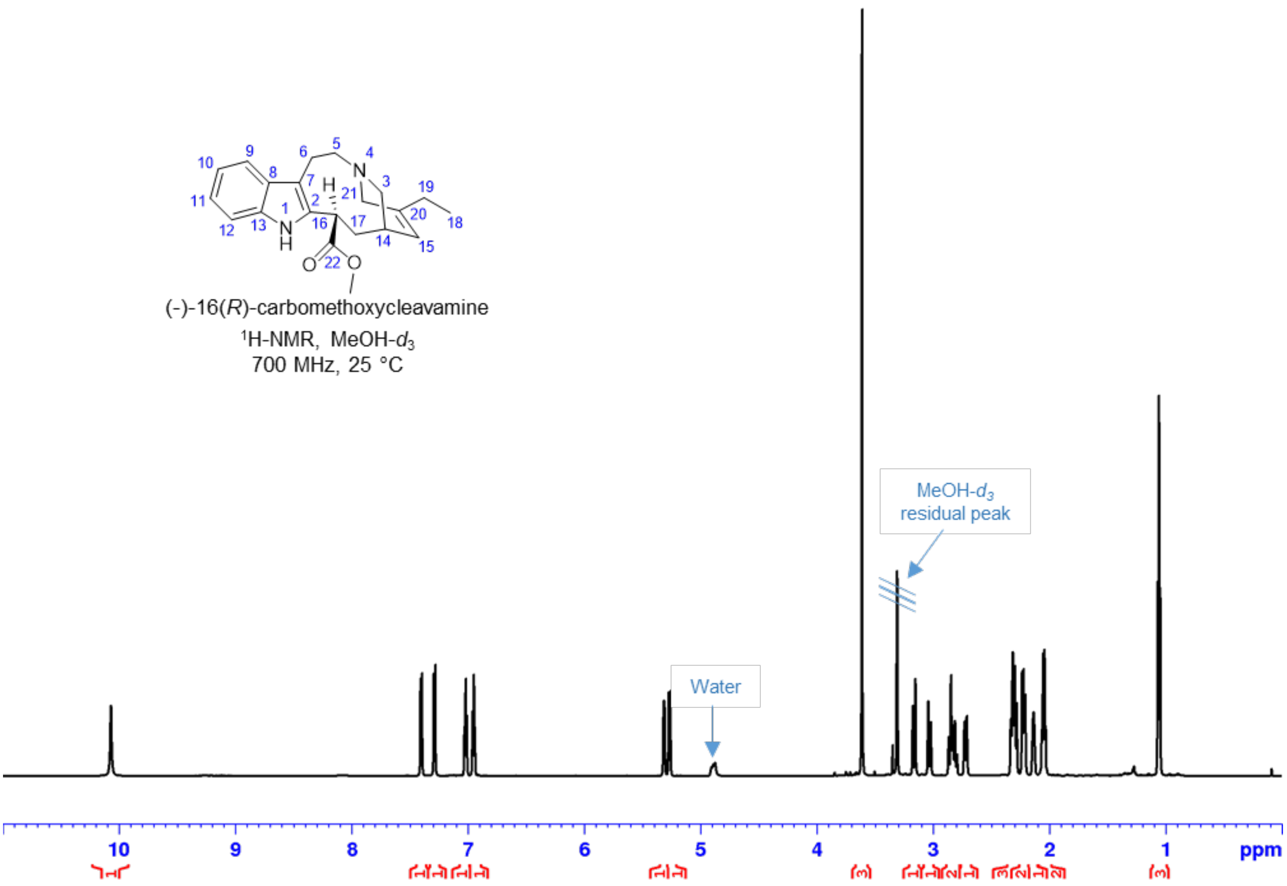


**(–)-16(*R*)-carbomethoxycleavamine (10):**
**$^{1}$H NMR** with water suppression (zgpr) aromatic range in MeOH-*d*$_3$

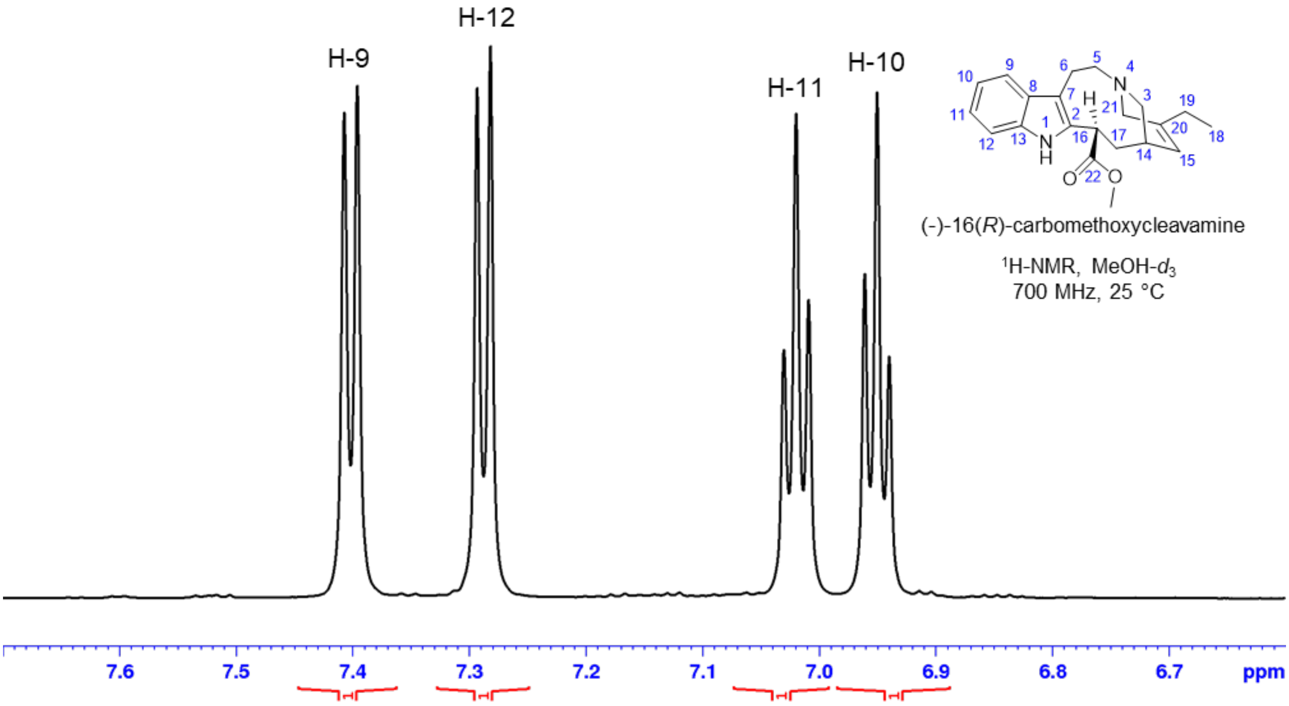

**(–)-16(*R*)-carbomethoxycleavamine (10):**
**$^1$H NMR** with water suppression (zgpr) 5.0-5.5 ppm in MeOH-$d_3$

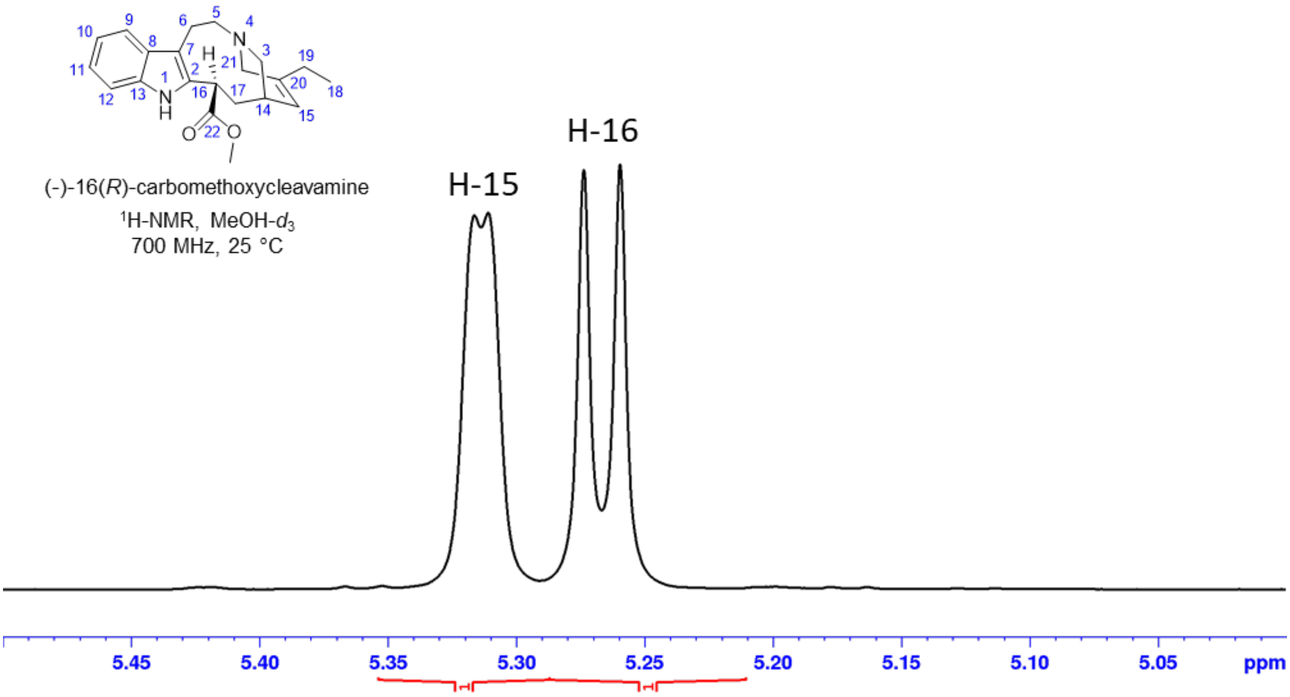


**(–)-16(*R*)-carbomethoxycleavamine (10):**
**$^1$H NMR** with water suppression (zgpr) aliphatic range in MeOH-$d_3$

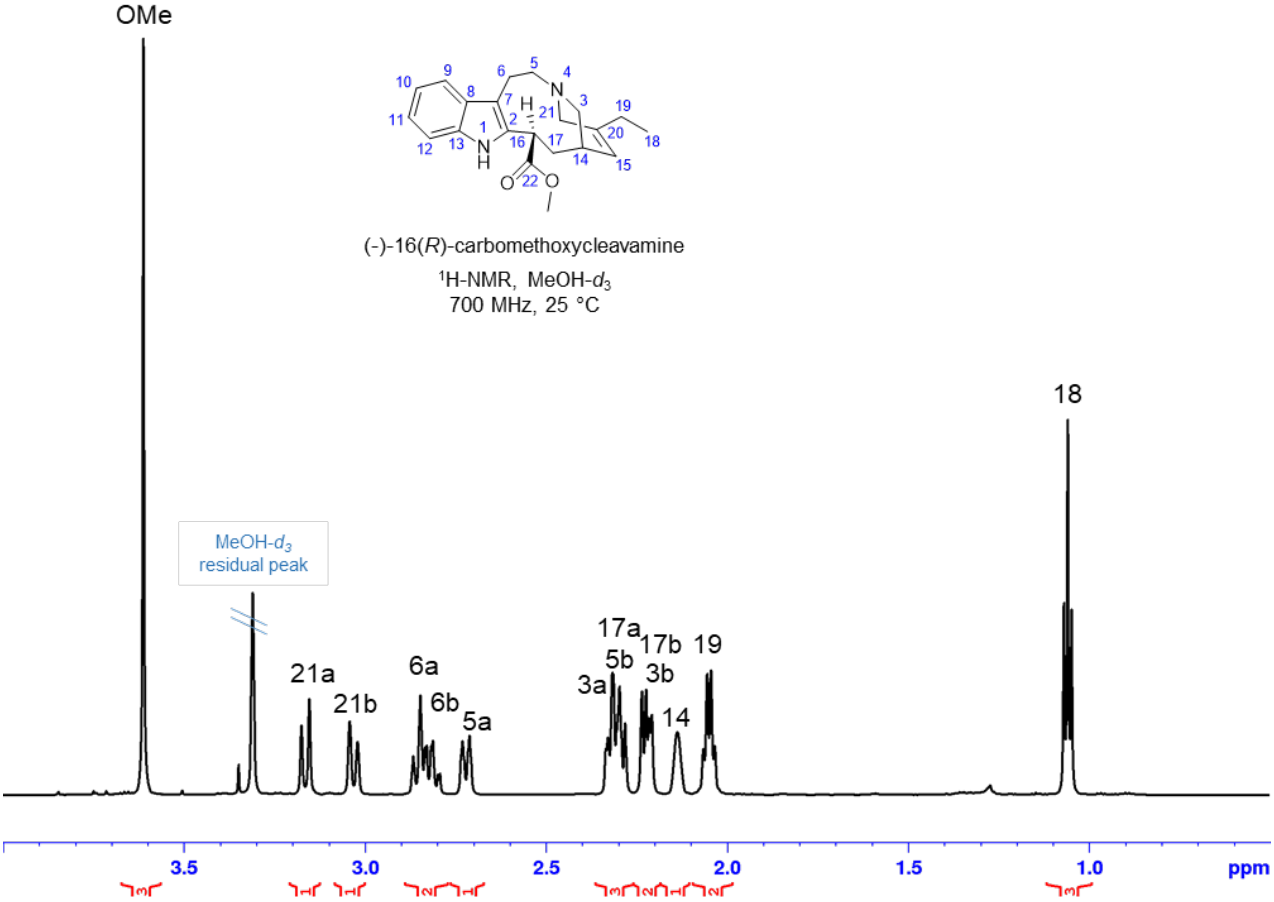

**(–)-16(*R*)-carbomethoxycleavamine (10):**
**Phase sensitive HSQC**, full range in MeOH-$d_3$
Shaded areas mark impurities and solvent, red: $CH_2$, black: CH, $CH_3$

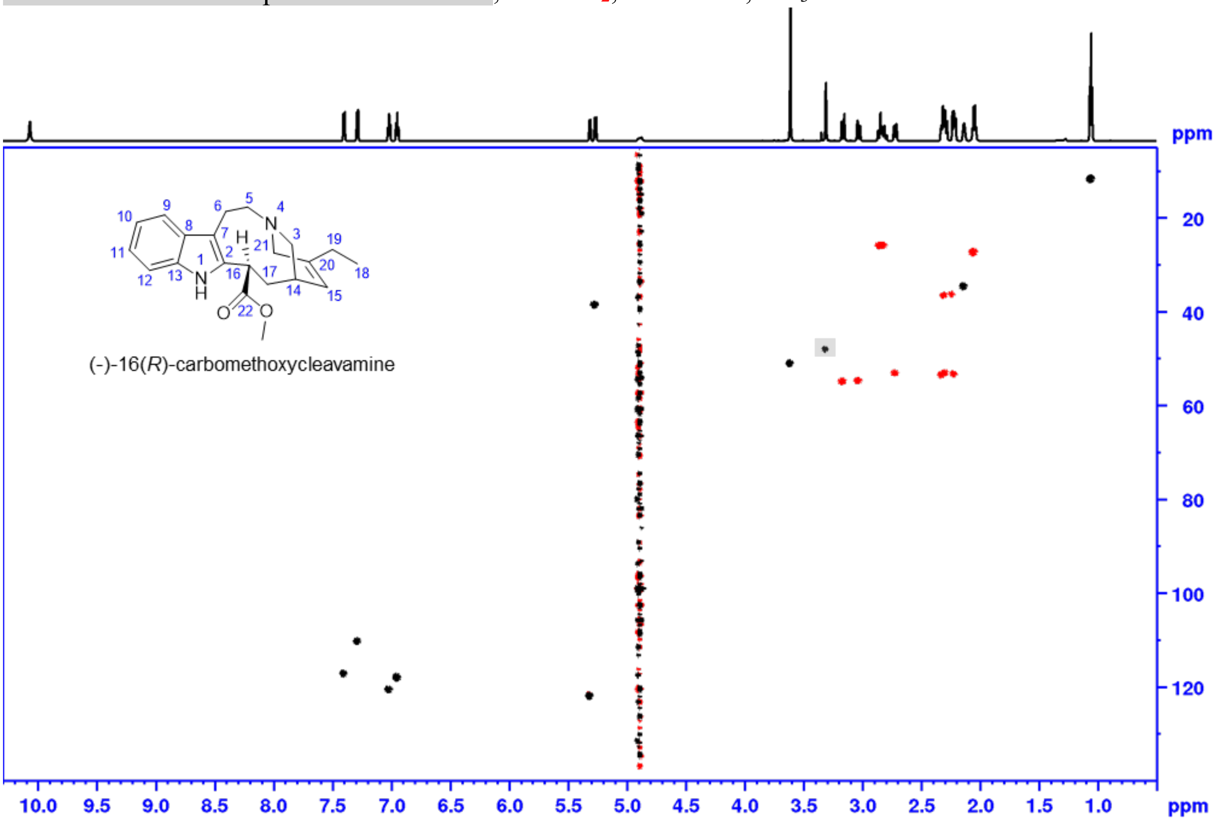


**(–)-16(*R*)-carbomethoxycleavamine (10):**
**Phase sensitive HSQC**, aliphatic range in MeOH-$d_3$
black: CH, $CH_3$

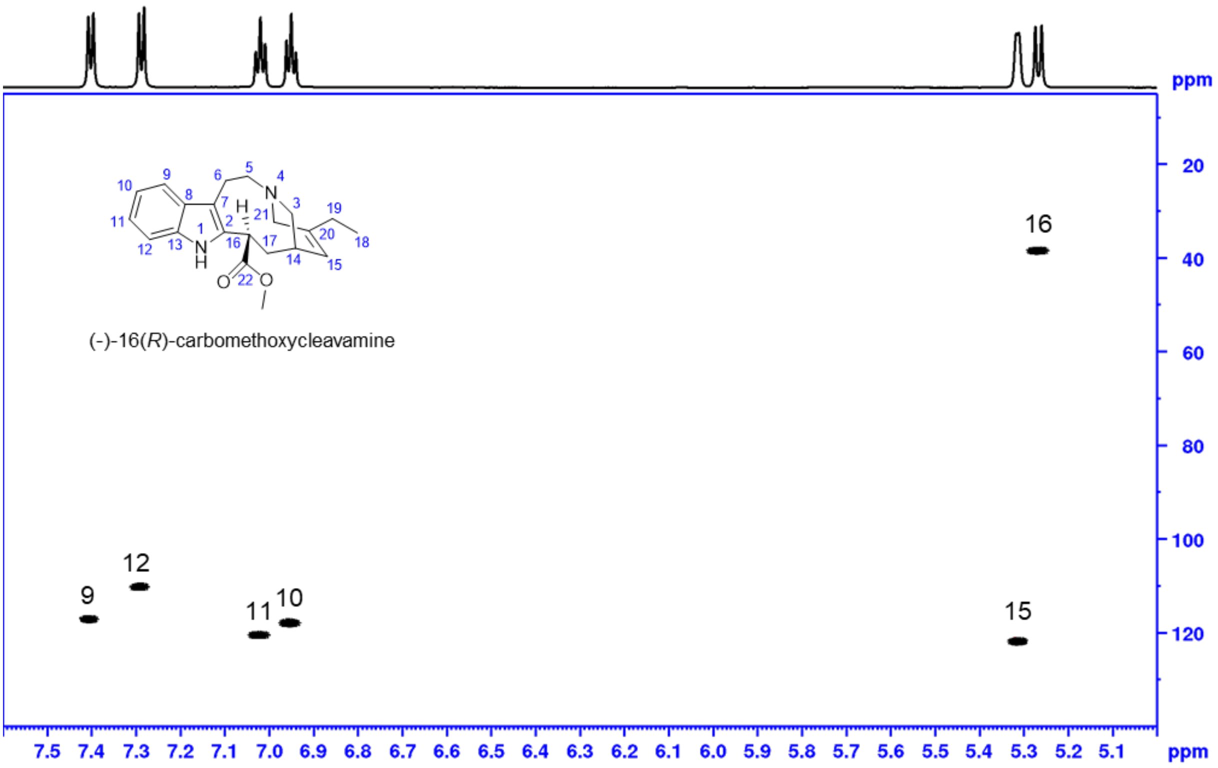

**(–)-16(*R*)-carbomethoxycleavamine (10): Phase sensitive HSQC**, aliphatic range in MeOH-$d_3$

Shaded areas mark impurities and solvent, red: $CH_2$, black: CH, $CH_3$

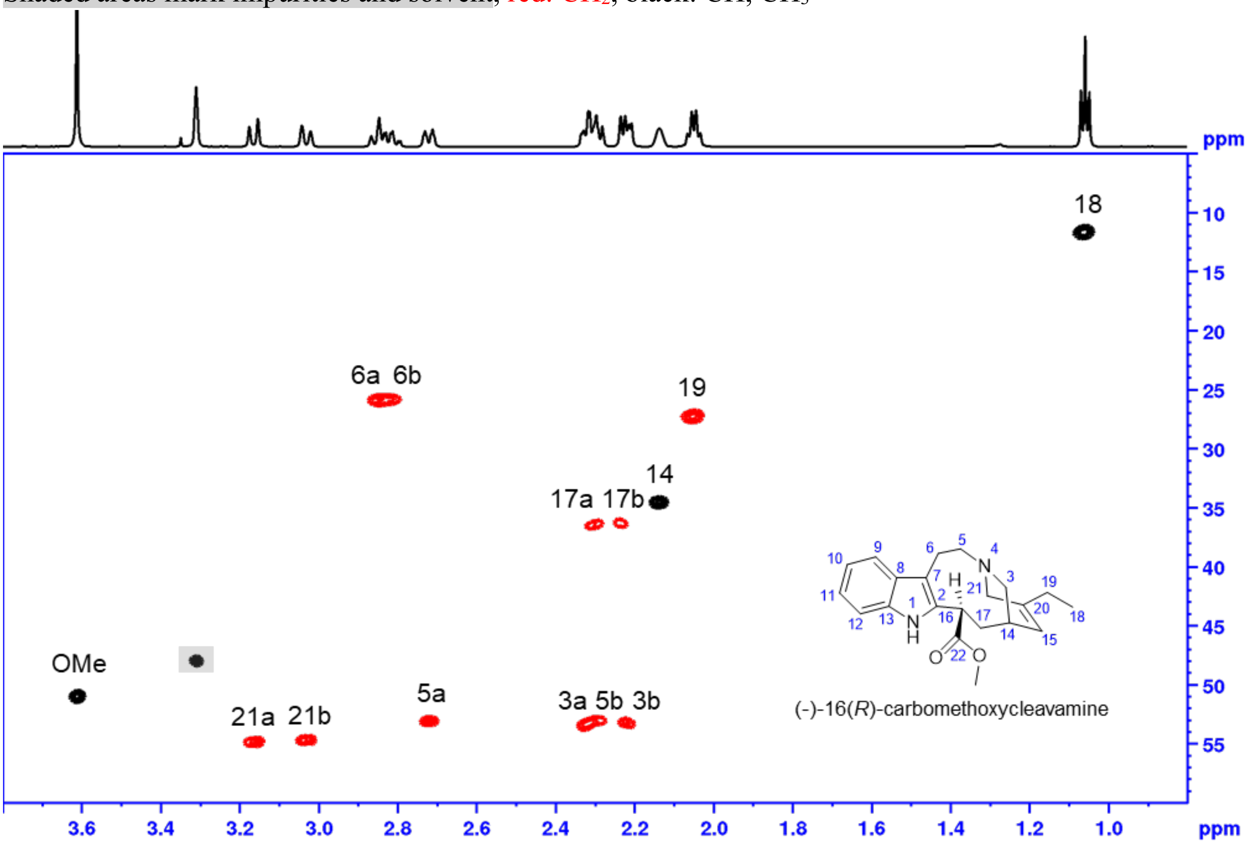


**(–)-16(*R*)-carbomethoxycleavamine (10): HMBC**, full range in MeOH-$d_3$

Shaded areas mark impurities and solvent

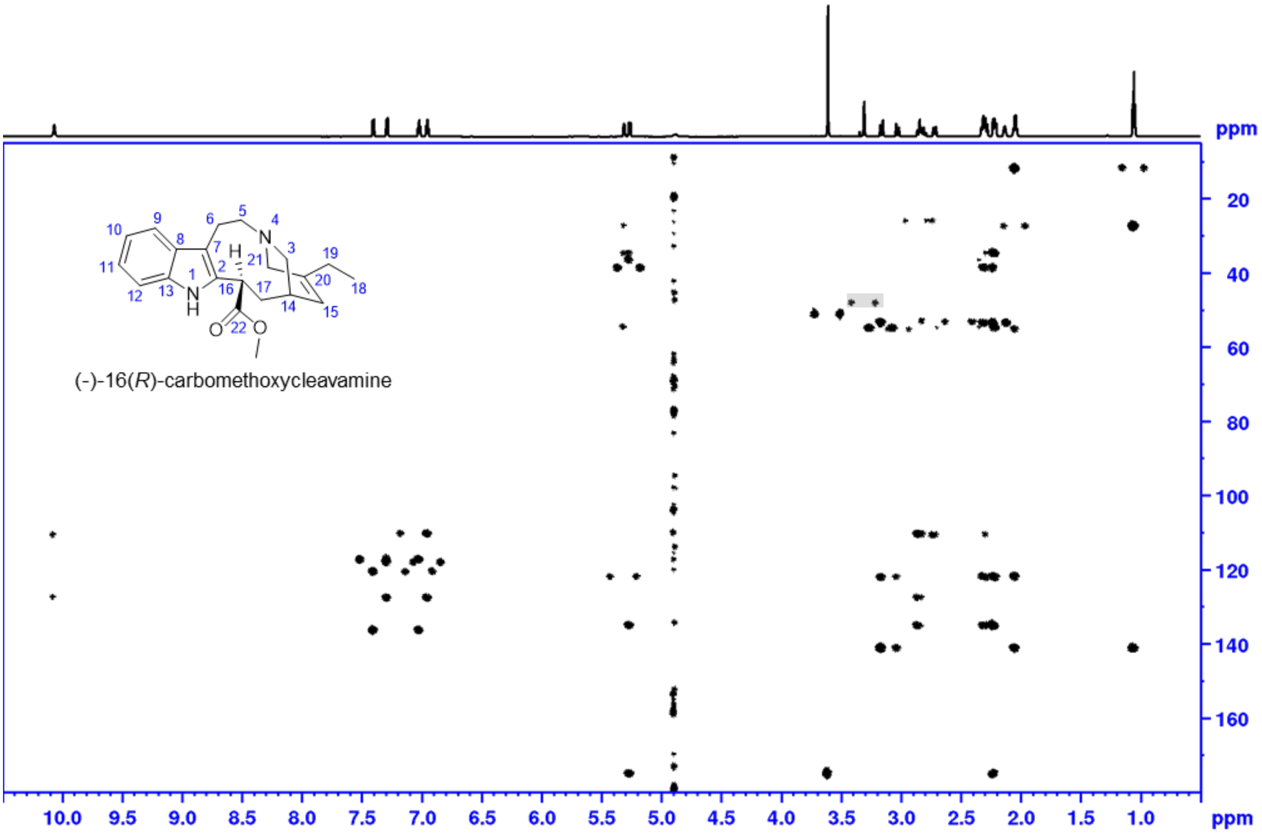

**(–)-16(*R*)-carbomethoxycleavamine (10): HMBC**, aromatic range in MeOH-$d_3$

Shaded areas mark impurities and solvent

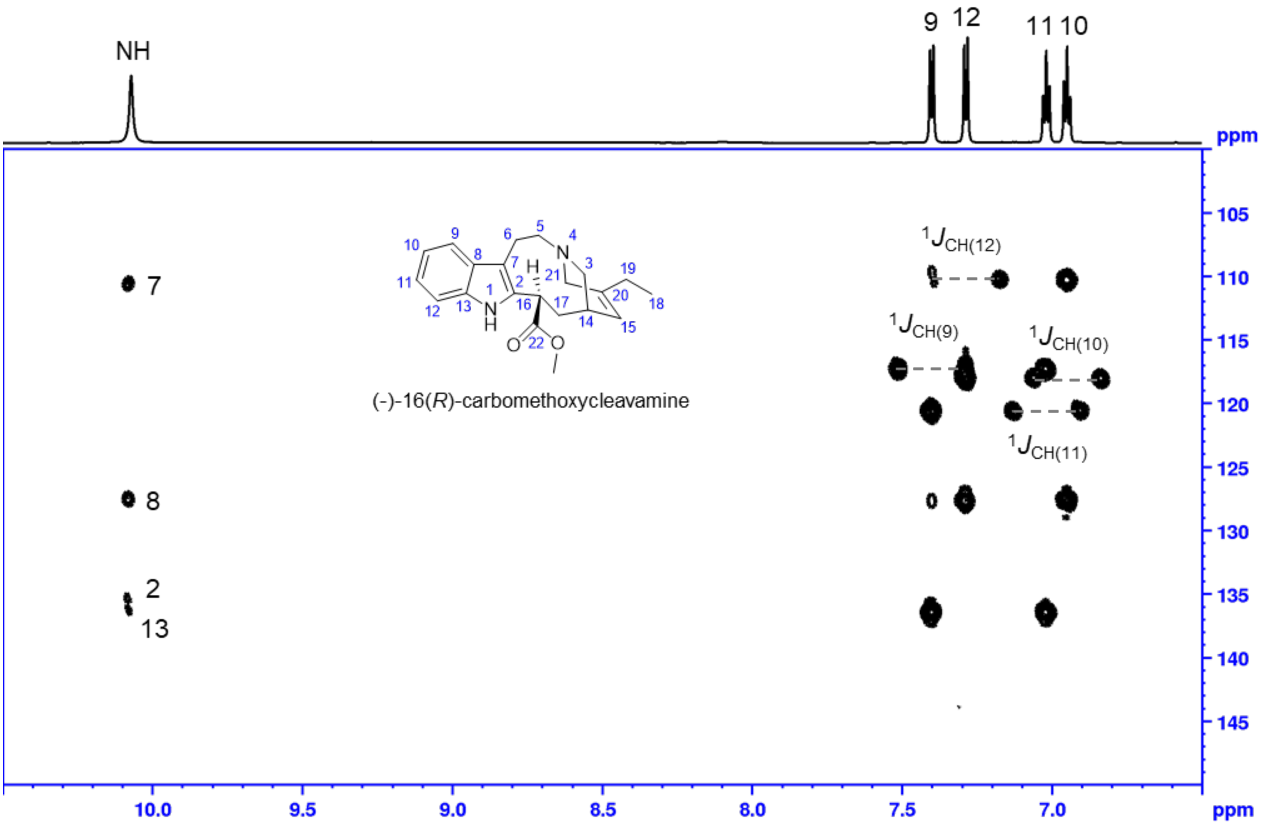


**(–)-16(*R*)-carbomethoxycleavamine (10): HMBC**, in MeOH-$d_3$

5.0-5.7 ppm in MeOH-$d_3$

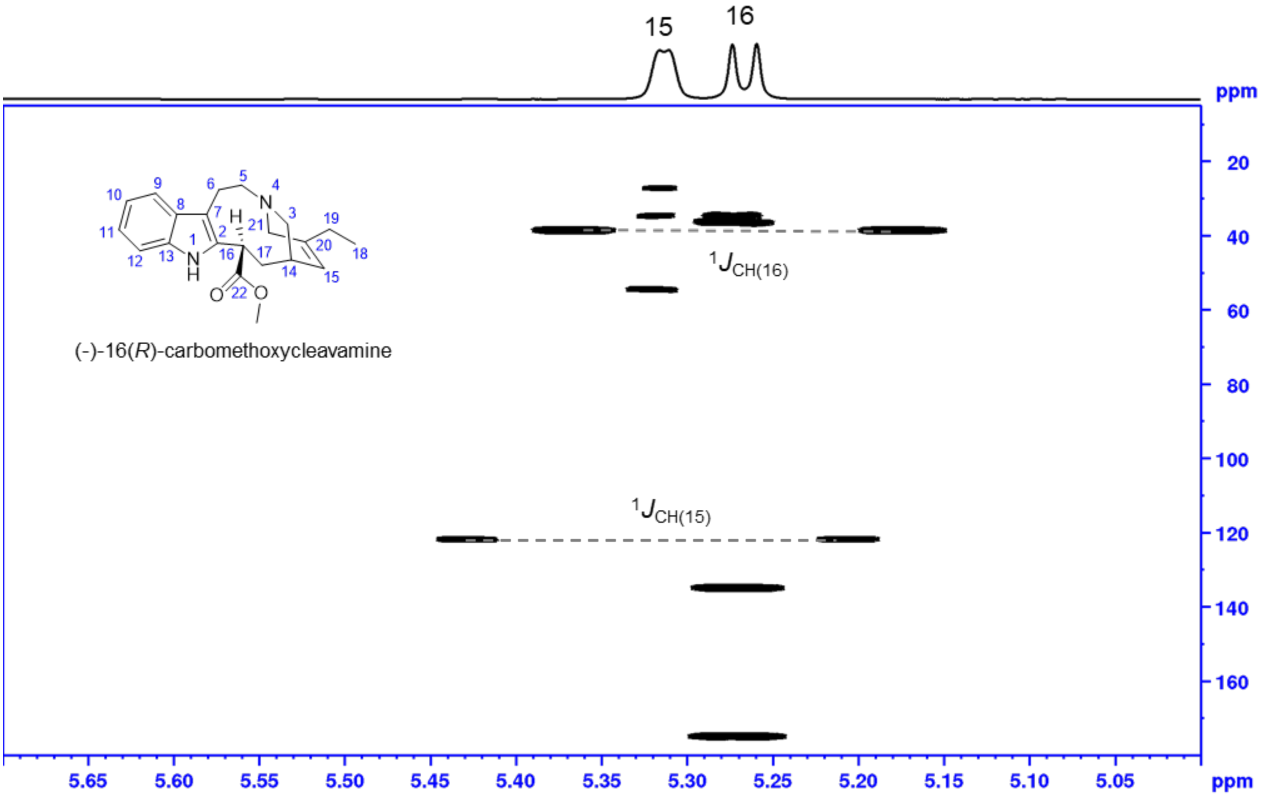

**(–)-16(*R*)-carbomethoxycleavamine (10): HMBC**, aliphatic range in MeOH-$d_3$

Shaded areas mark impurities and solvent

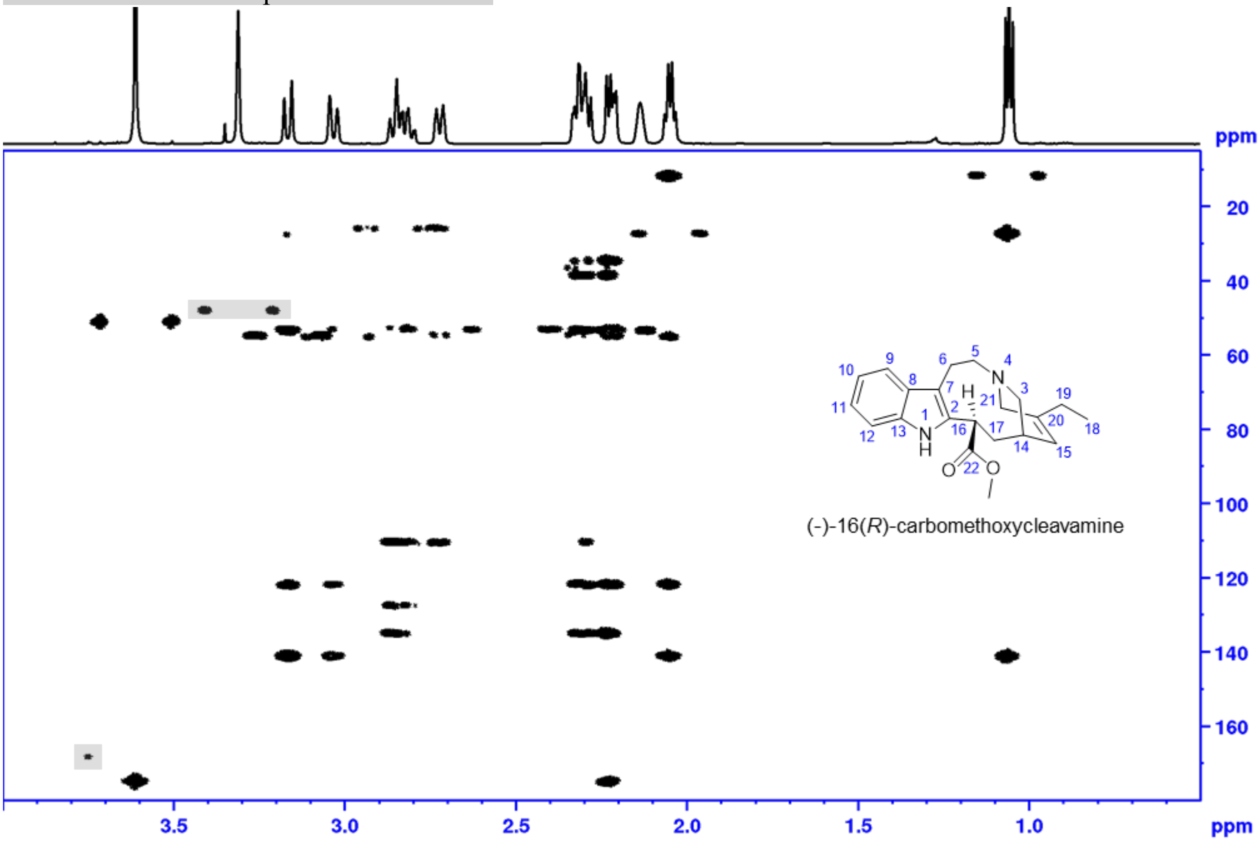


**(–)-16(*R*)-carbomethoxycleavamine (10)**

**$^1$H-$^1$H DQF COSY** with water suppression, full range in MeOH-$d_3$

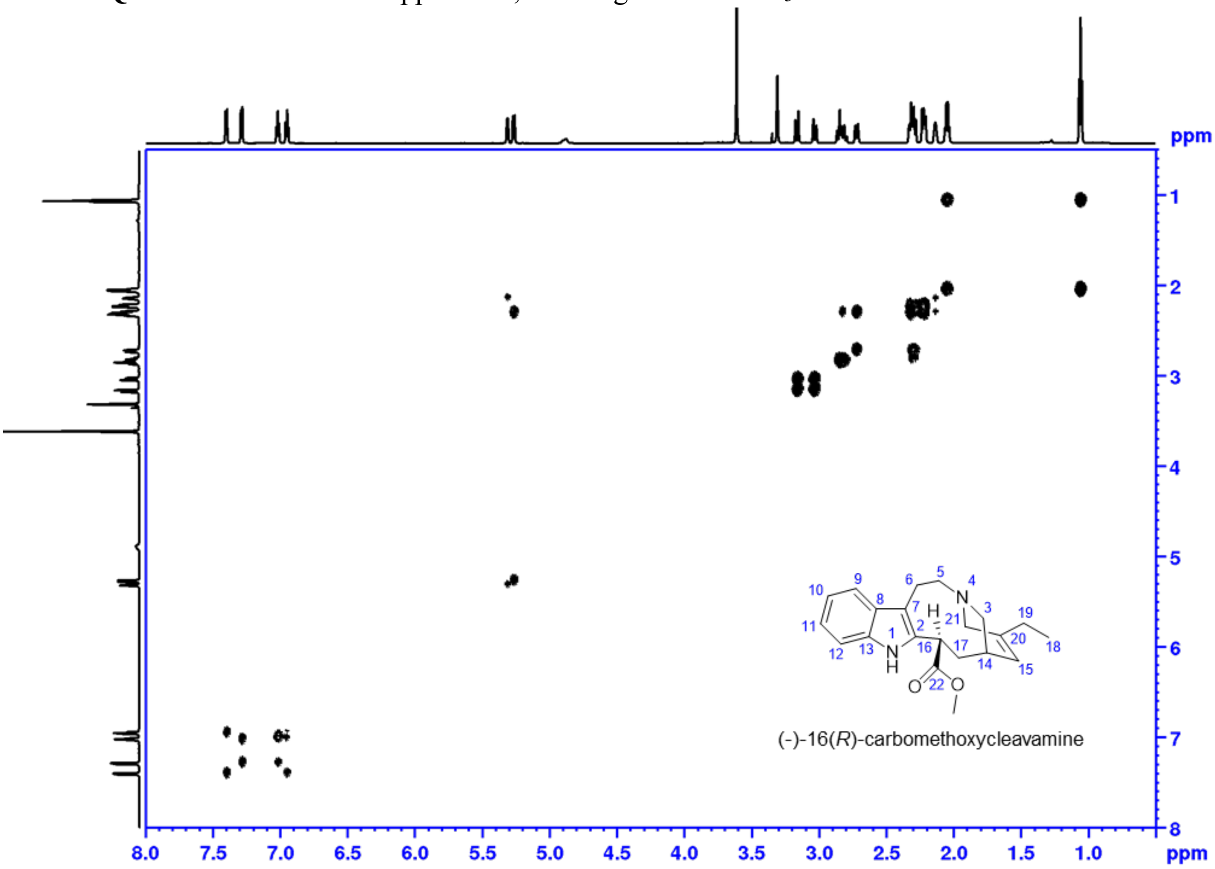

**(–)-16(*R*)-carbomethoxycleavamine (10)**

**$^{1}$H-$^{1}$H DQF COSY** with water suppression, aromatic range in MeOH-$d_3$

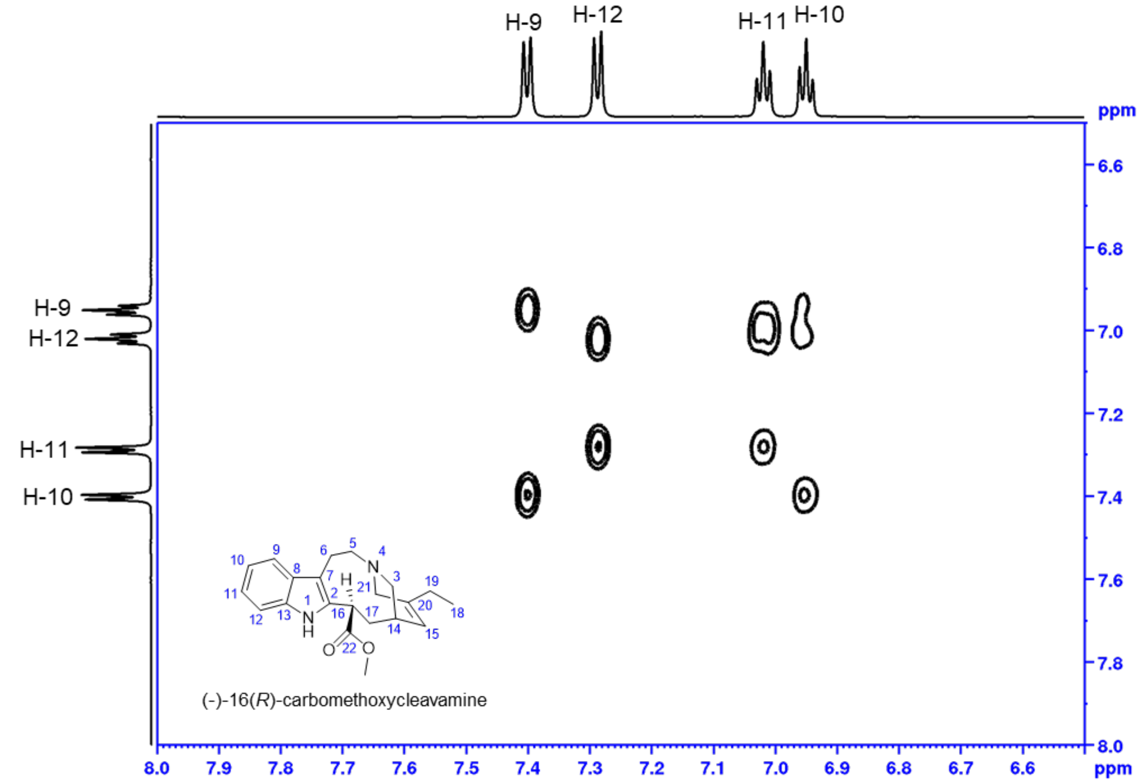


**$^{1}$H-$^{1}$H DQF COSY** with water suppression, aliphatic range in MeOH-$d_3$

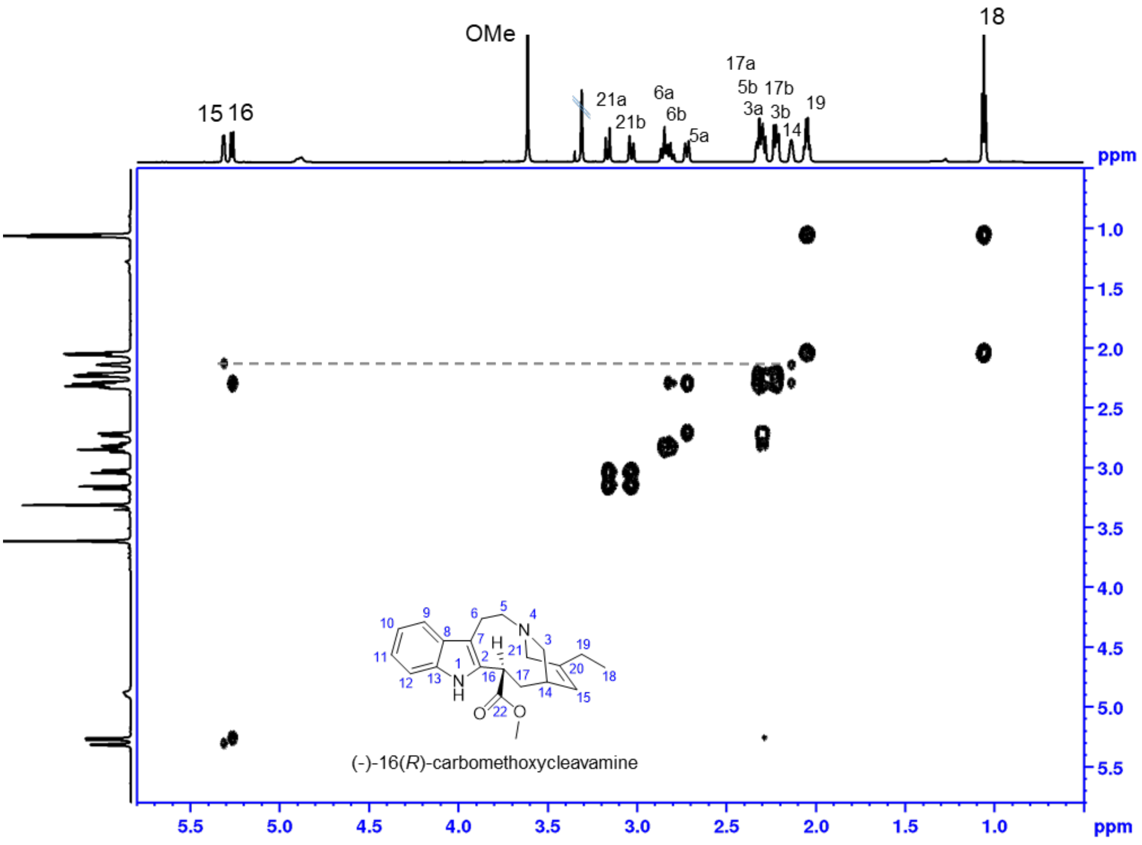

**(–)-16(*R*)-carbomethoxycleavamine (10):**
**$^{1}$H-$^{1}$H ROESY** with water suppression, full range in MeOH-$d_3$

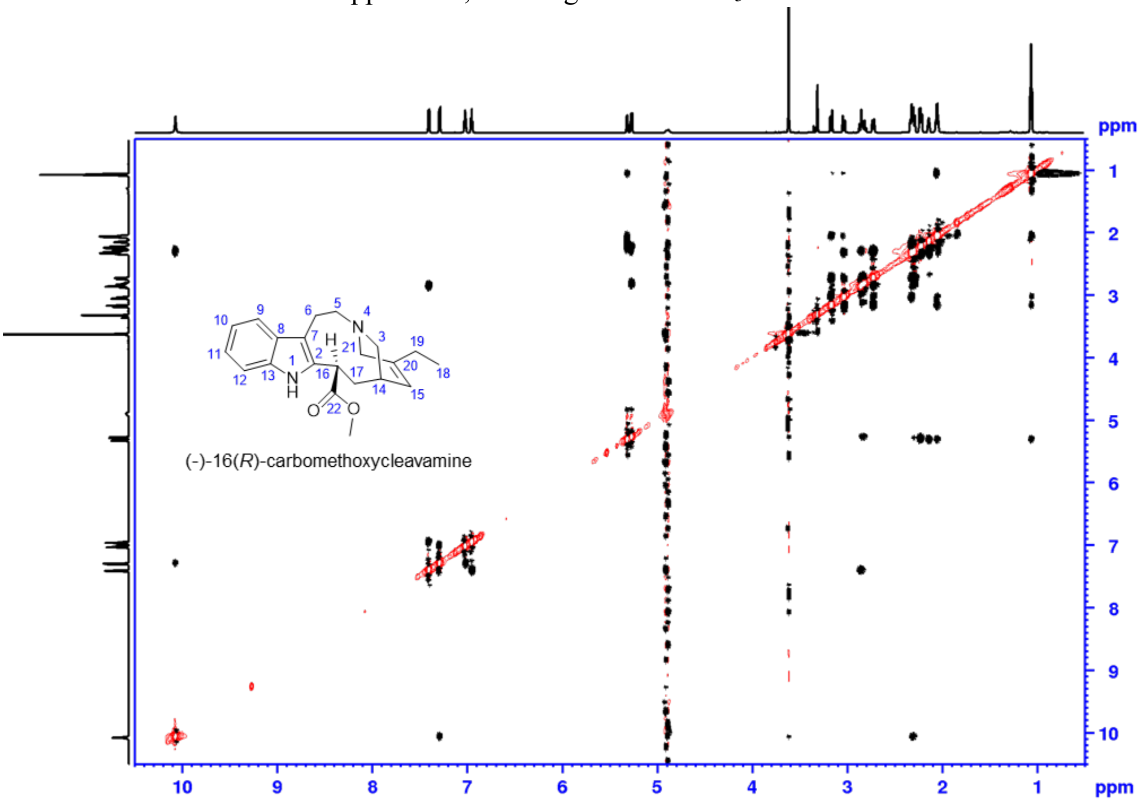


**(–)-16(*R*)-carbomethoxycleavamine (10)**
**$^{1}$H-$^{1}$H ROESY** with water suppression, aromatic range in MeOH-$d_3$

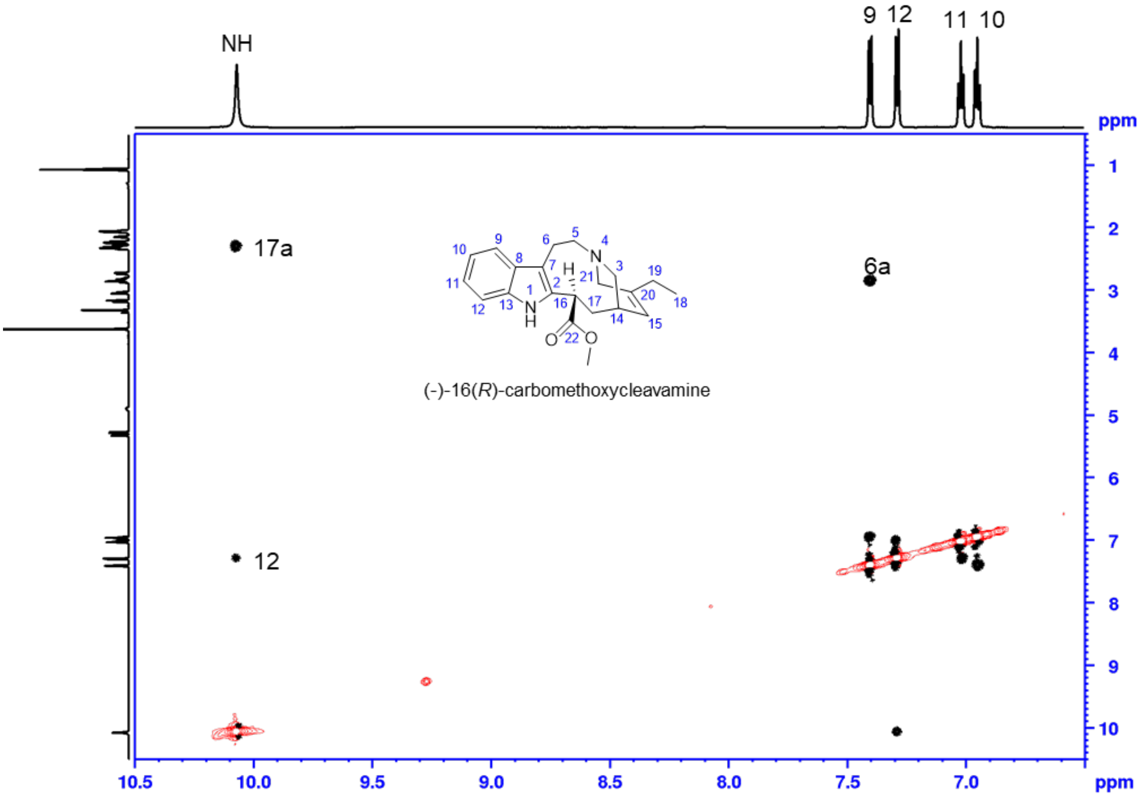

**(–)-16(*R*)-carbomethoxycleavamine (10)**
**$^{1}$H-$^{1}$H ROESY** with water suppression, 5.0-5.8 ppm, in MeOH-$d_3$

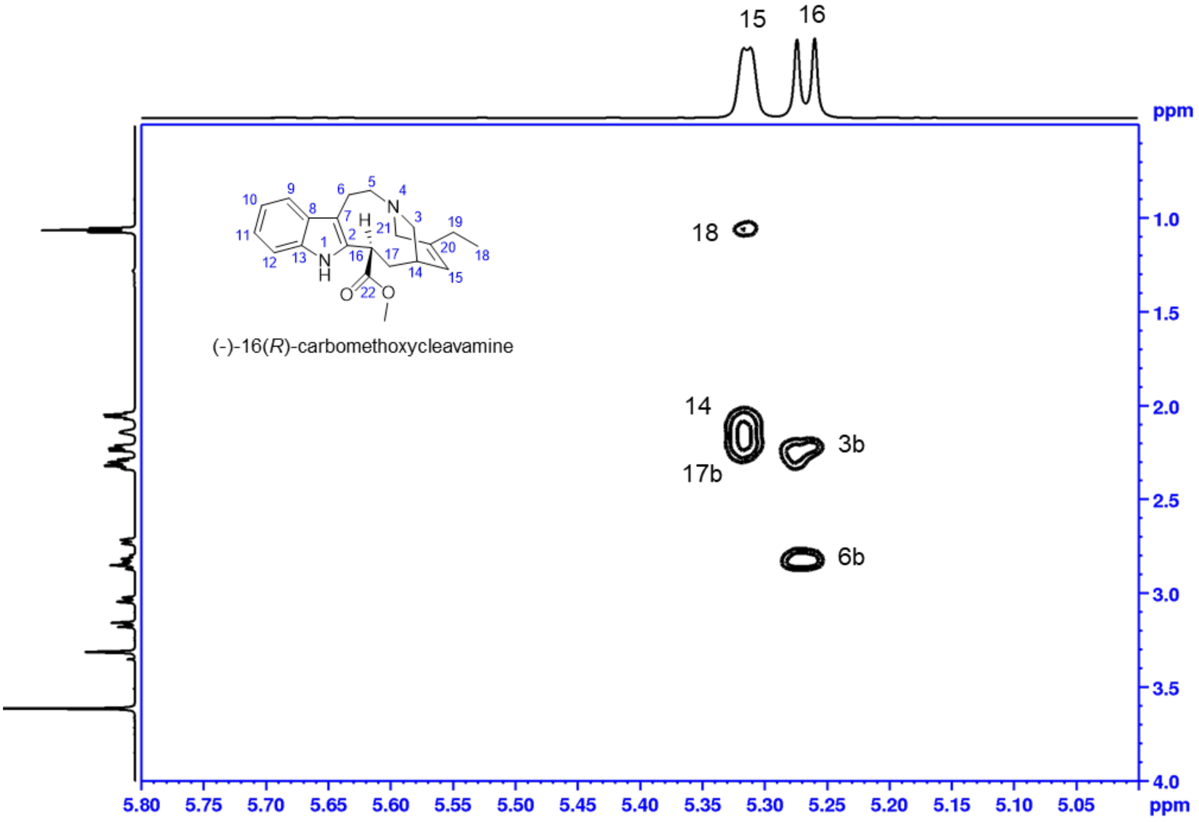


**(–)-16(*R*)-carbomethoxycleavamine (10)**
**$^{1}$H-$^{1}$H ROESY** with water suppression aliphatic range in MeOH-$d_3$

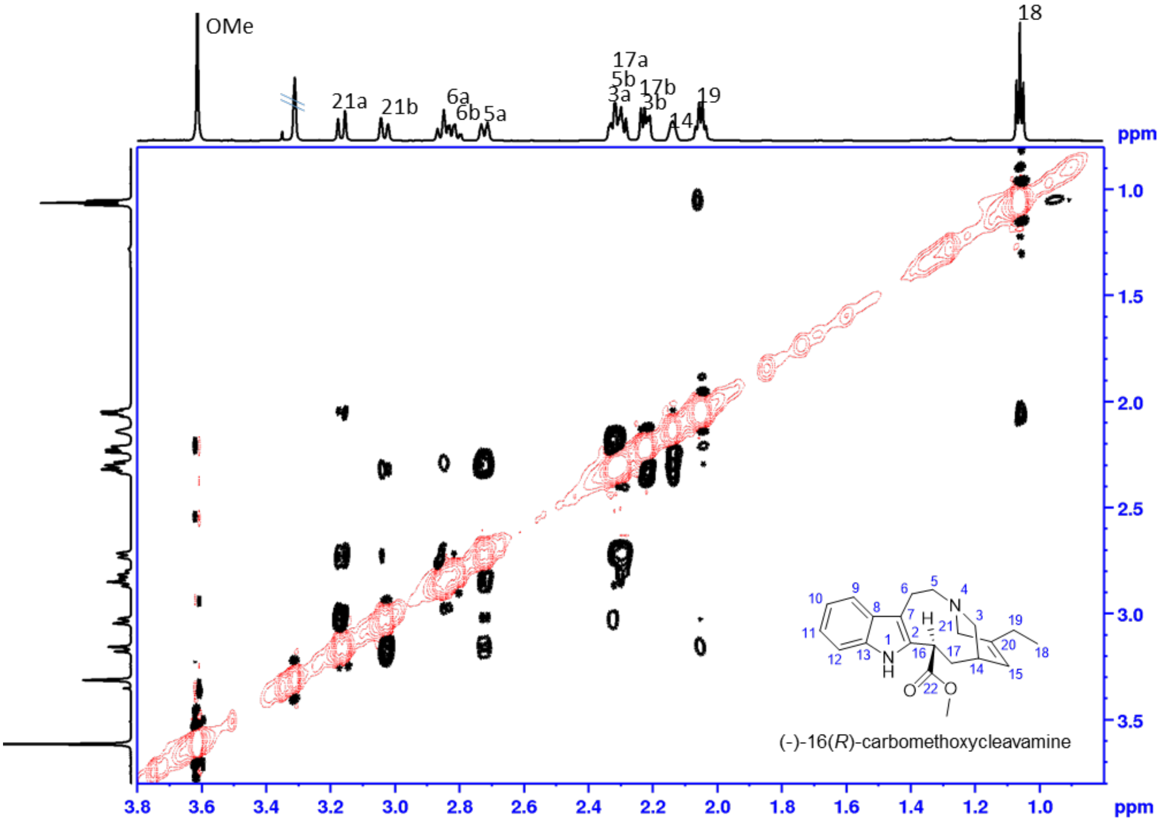

**(–)-16(*R*)-carbomethoxycleavamine (10)**

**$^1$H-$^1$H ROESY** with water suppression aliphatic range in MeOH-$d_3$

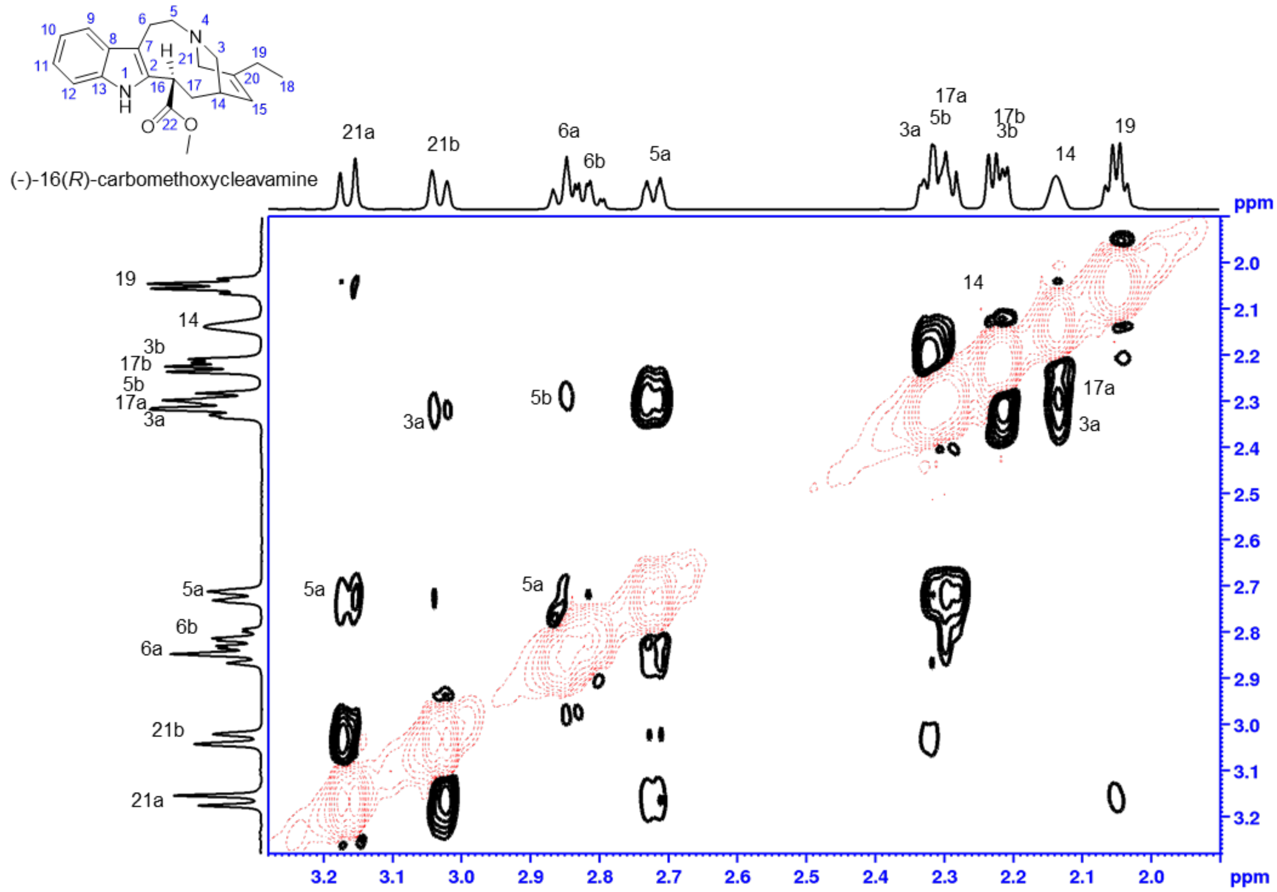


**Structure of (–)-16(*R*)-carbomethoxycleavamine (10)**. Structure optimized Gaussian 16W (DFT, B3LYP-6-311G+(d,p), solvent MeOH). Important ROESY correlations are depicted in green.

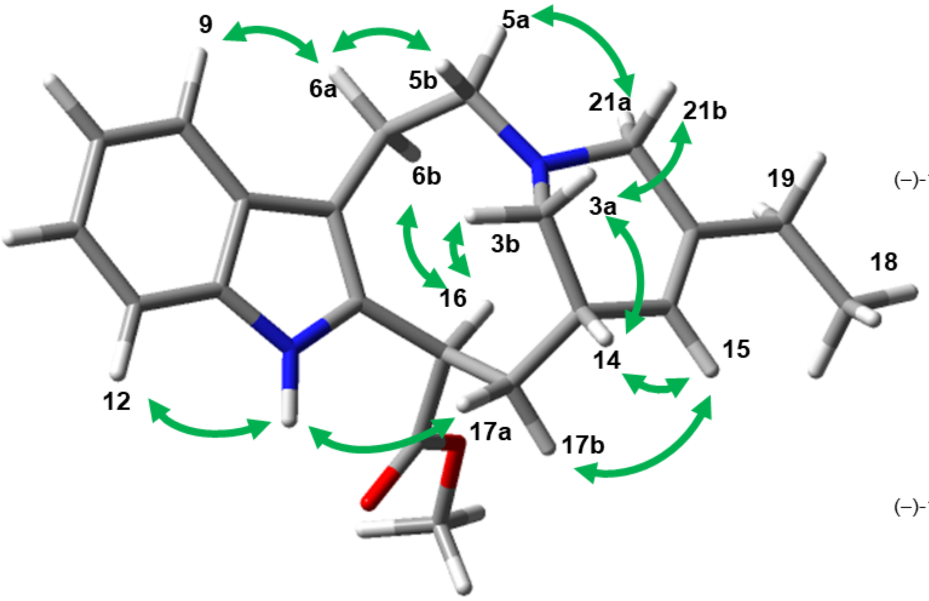


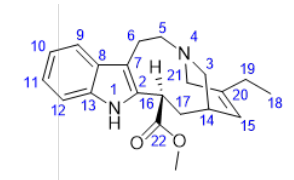

(–)-16(*R*)-carbomethoxycleavamine (**10**)

|||

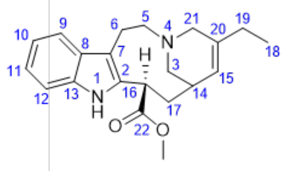

(–)-16(*R*)-carbomethoxycleavamine (**10**)

**NMR data for (–)-16(*R*)-carbomethoxycleavamine-*d* (10-d).**

$^{1}$**H NMR** with water suppression (zgpr) full range in MeOH-$d_3$

Shaded areas (grey bars) mark impurities and solvent

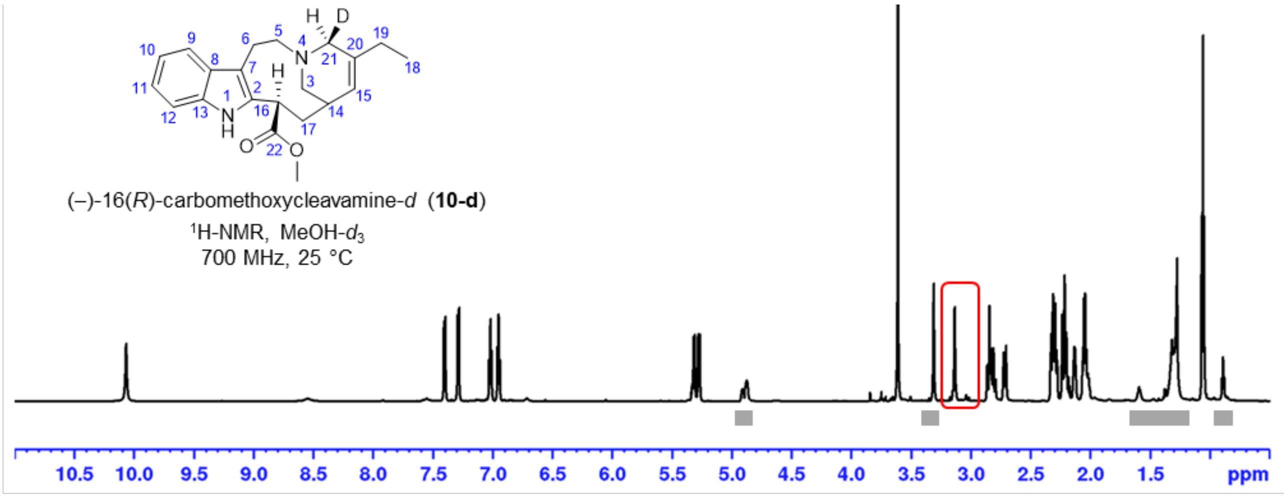


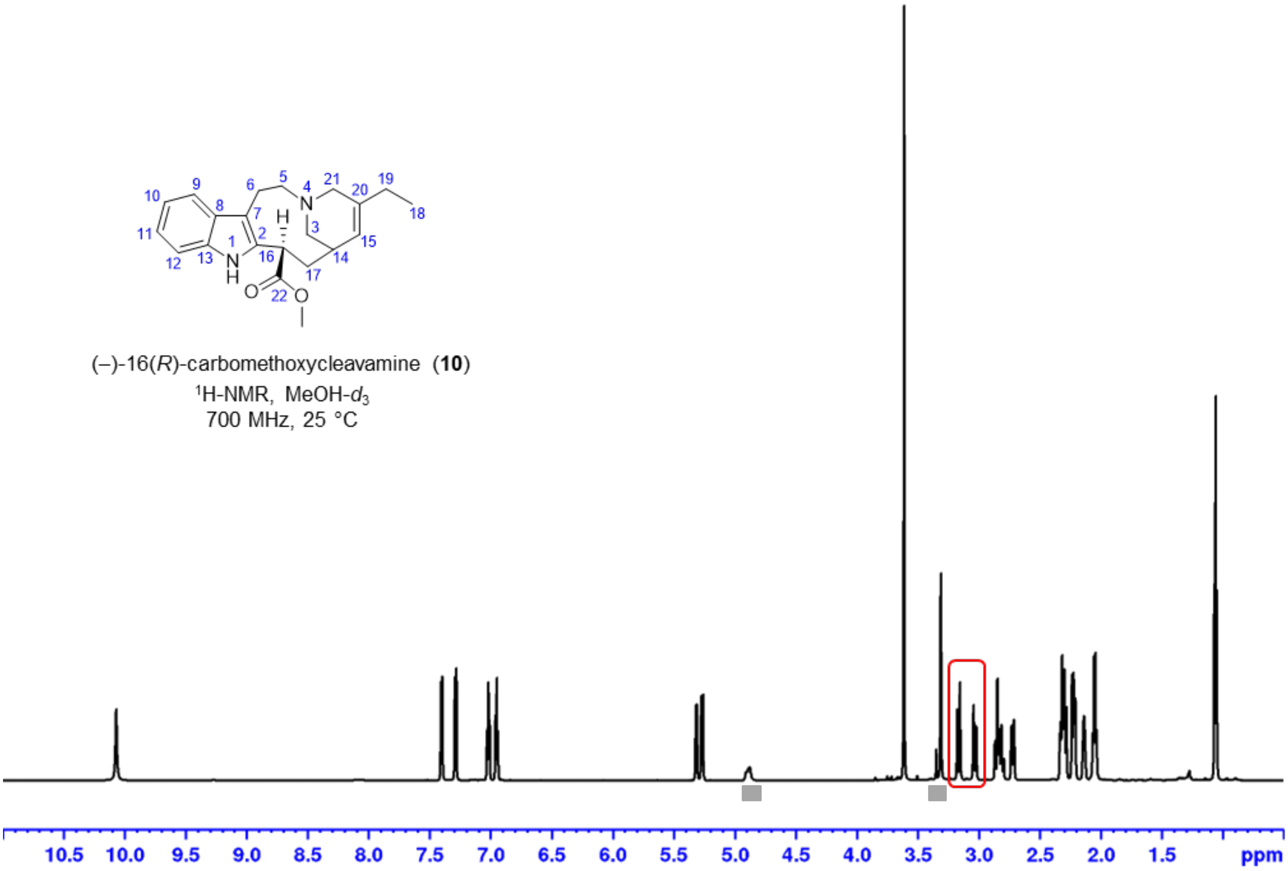

**$^1$H NMR** with water suppression (zgpr) 2.5 – 3.5 ppm in MeOH-$d_3$

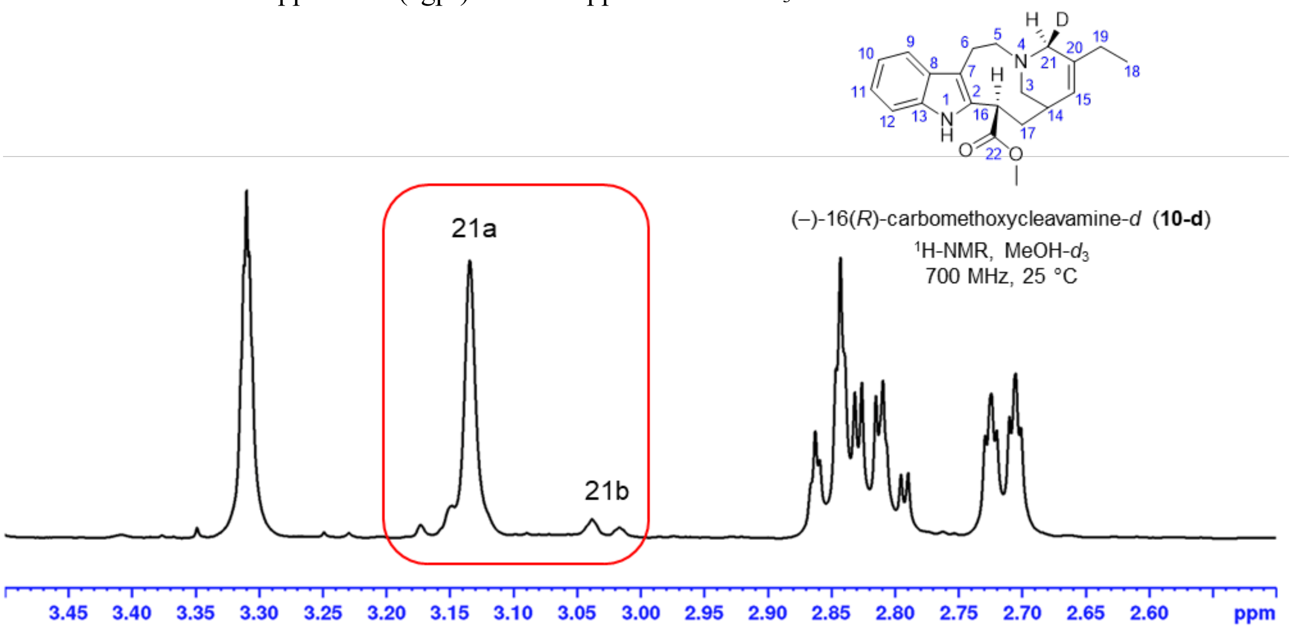


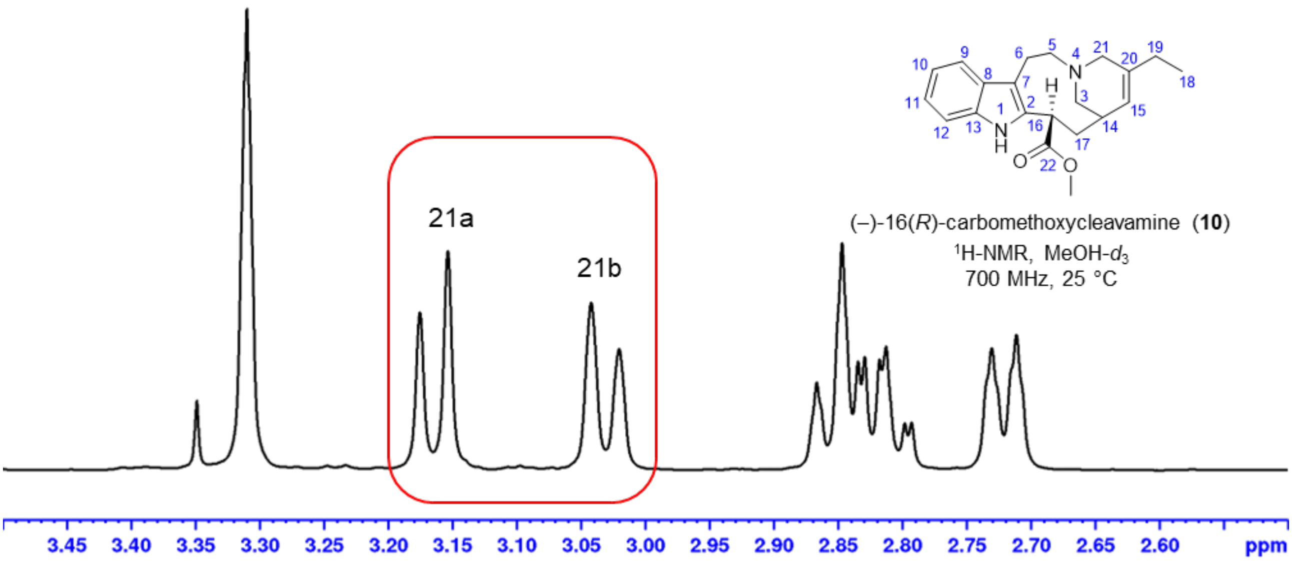

**(–)-16(*R*)-carbomethoxycleavamine-*d* (10-d): Phase sensitive HSQC**, full range in MeOH-$d_3$
Shaded areas mark impurities and solvent, red: $CH_2$, black: CH, $CH_3$

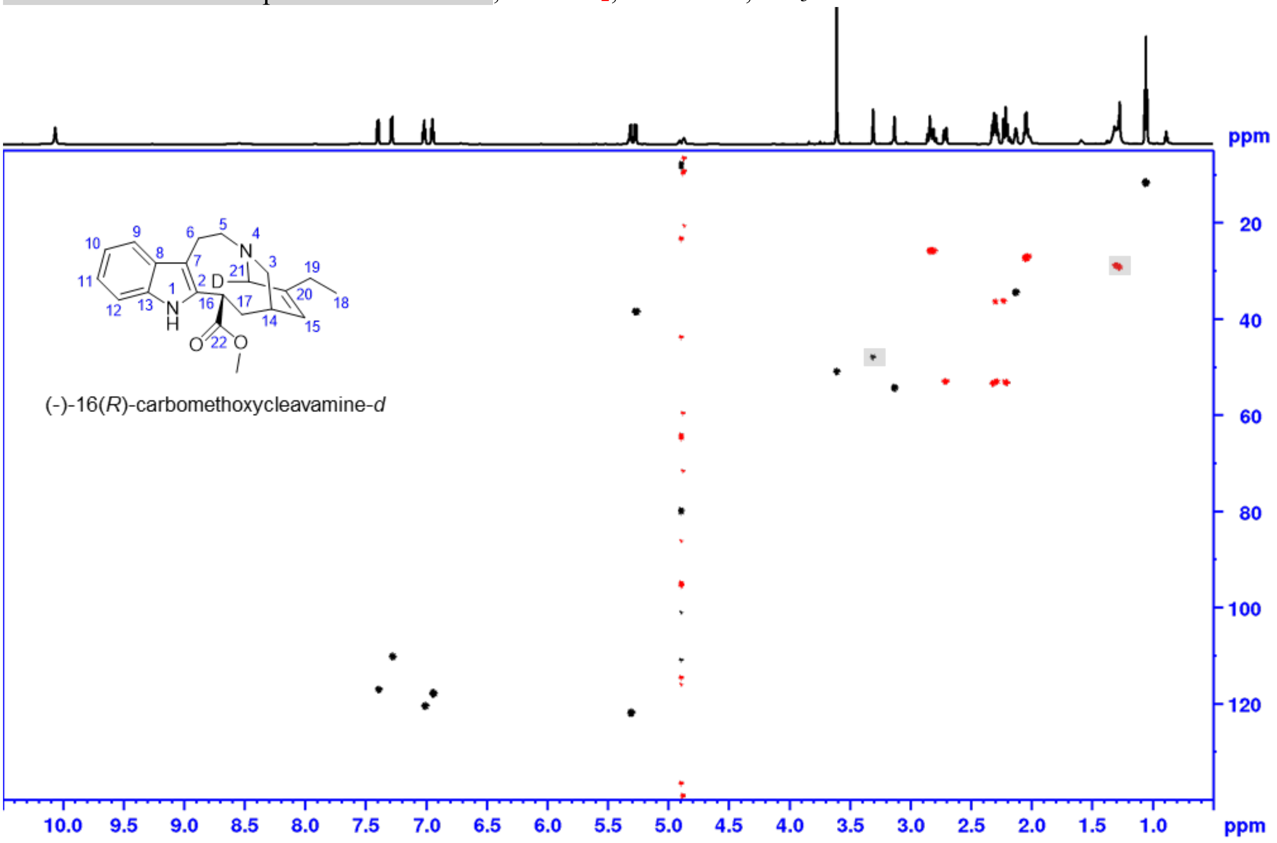


**(–)-16(*R*)-carbomethoxycleavamine-*d* (10-d): Phase sensitive HSQC**, aliphatic range in MeOH-$d_3$
Shaded areas mark impurities and solvent, red: $CH_2$, black: CH, $CH_3$

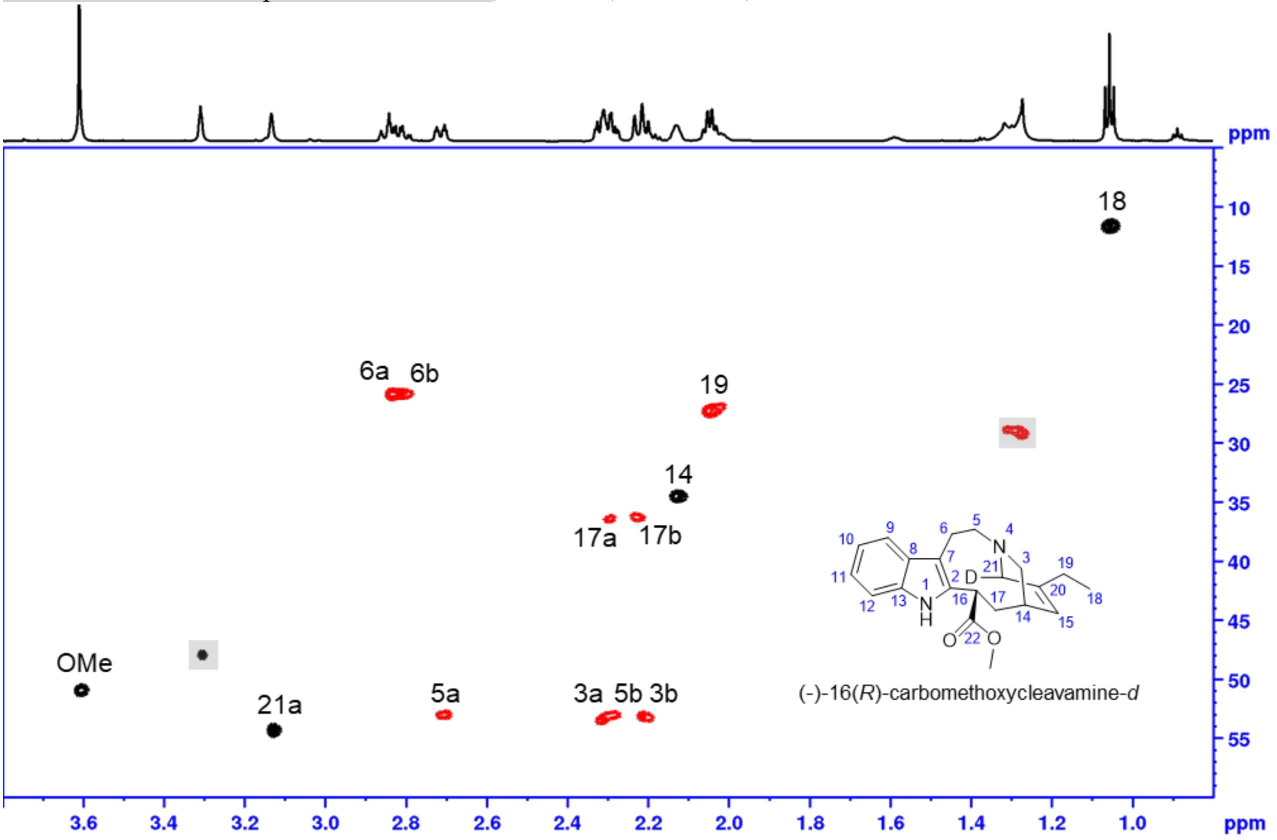

**Comparison of the NMR spectra of (–)-16(*R*)-carbomethoxycleavamine (10) and (+)-16(*S*)-carbomethoxycleavamine (10a) (synthesized from (+)-catharanthine (3)).**

$^{1}$**H NMR** with water suppression (zgpr) full range in MeOH-$d_3$

Shaded areas (grey bars) mark impurities and solvent

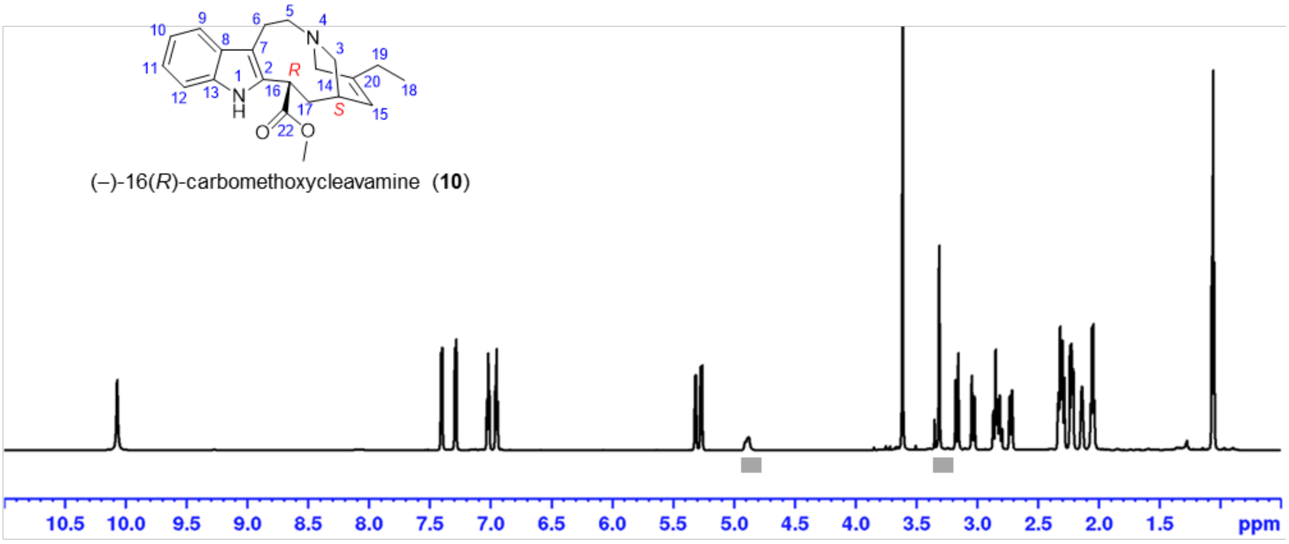


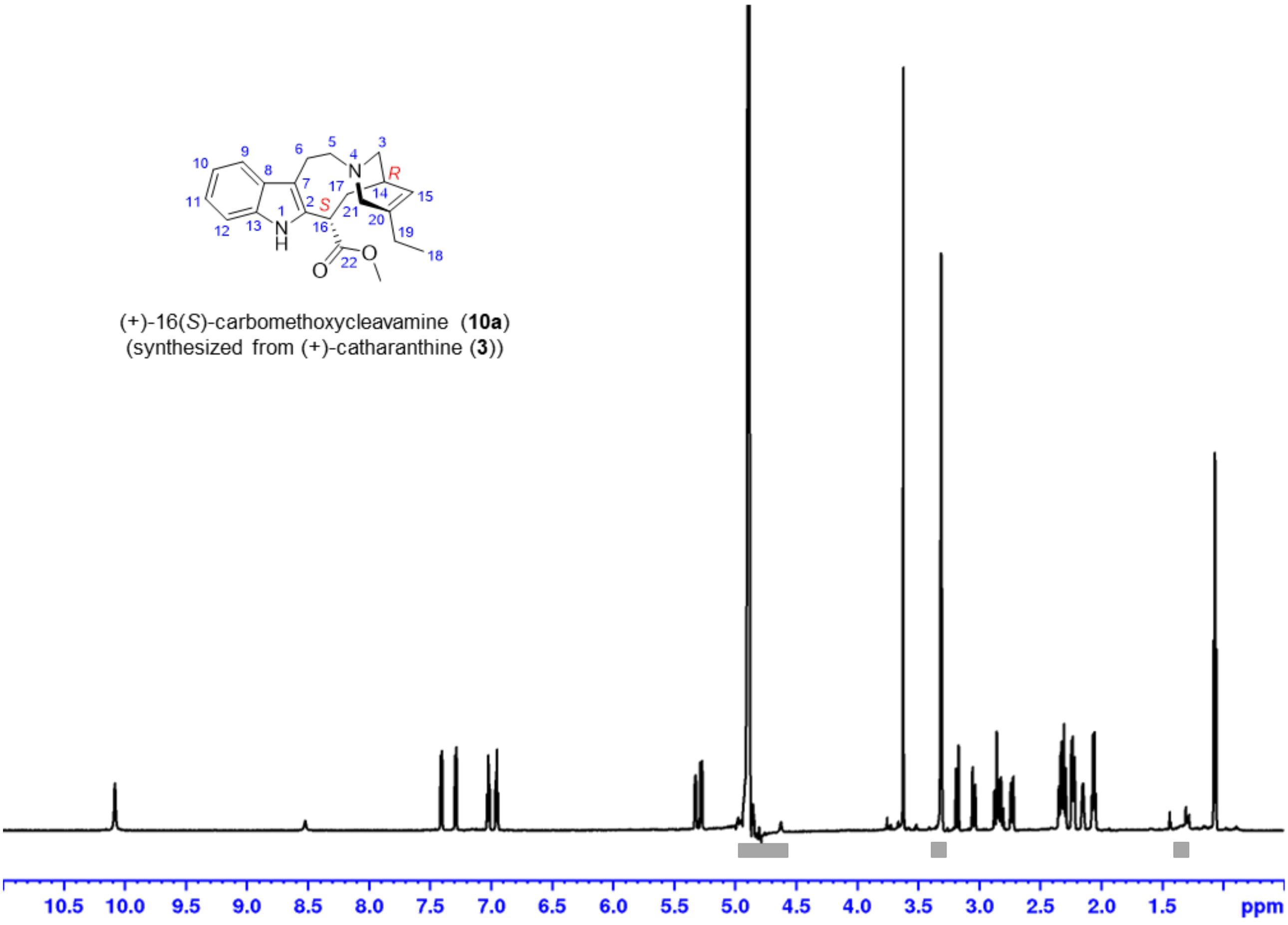
